# Are our 95% CIs only worth 45% confidence? A heterogeneity-aware method for extending inference

**DOI:** 10.1101/2025.08.05.668631

**Authors:** Ulrich Knief, Wolfgang Forstmeier

**Author notes:** Addresses for correspondence: Ulrich Knief, Evolutionary Biology and Ecology, Institute of Biology I (Zoology), University of Freiburg, Hauptstr. 1, 79104 Freiburg, Germany, Wolfgang Forstmeier, Department of Ornithology, Max Planck Institute for Biological Intelligence, Eberhard-Gwinner-Str. 7, 82319 Seewiesen, Germany.

## Abstract

1. When multiple studies on the same research question, or multiple analyses of the same dataset are summarized in a meta-analysis, our confidence intervals (CIs) are put to a relentless test of reliability. While we might hope that 95% of the CIs would contain the meta-analytic mean value (and therefore presumably also the true value), a recent meta-analysis of 512 meta-analyses in ecology and evolution suggests that only a sobering 45% of them do.
2. As this paradox of overconfidence continues to confuse researchers, we attempt to explain where most of the heterogeneity in findings might be coming from. Much of it is unsurprising, because conventional 95% CIs reflect only sampling noise and ignore other sources of error such as arbitrary analysis decisions (“model uncertainty”). Being aware of multiple sources of error beyond sampling noise, the replication crisis logically follows from an anticonservative statistical practice that allows for overinterpretation beyond the legitimate inference space, which in fact is narrower than commonly acknowledged.
3. We explain how to calculate extended confidence intervals (CI_ext_) that also cover other sources of biological and analytical heterogeneity, and we clarify which CI_ext_ is valid for which extended inference space. We further show how multiple versions of analysis of the same dataset can be merged into a many-analyses meta-analysis (MAMA) which yields a CI that accounts for two sources of error.
4. At the same time, we recognize that high heterogeneity estimates can be artefactual, arising from “comparing apples and oranges” (either in terms of statistical metric or biological interpretation). The estimate of only 45% deserved confidence therefore appears far too pessimistic. Overall, we argue for greater caution in interpreting both confidence intervals and heterogeneity estimates, and for clearer separation of comparable from non-comparable evidence.

## Introduction

Doing science can be a frustrating and sobering business! Imagine that we just published a study showing that factor *X* has a statistically significant effect on our variable of interest *Y*. We estimated that this effect is approximately of magnitude *m*, and we further estimated a 95% confidence interval (CI_lower_; CI_upper_), which, in 95% of all imaginary repetitions of the study, should contain the true mean effect *µ* (the true, but unknown, magnitude of the effect of *X* on *Y*). Now, here comes the frustrating part: it will not take long, and a colleague will publish another study on the same topic, finding a very different effect size that lies outside our proclaimed confidence interval, or, even worse, another colleague will publish a re-analysis of our own data, showing that our claim about magnitude *m* is no longer tenable when choosing a subtly different way of data analysis. All this is part of our daily business of doing science, and we are highly versed in finding *post-hoc* explanations for why the other study or the other way of data analysis may be invalid, and therefore does not threaten our initial findings and conclusions.

Such daily drama of attempting to verify the findings of others can also be summarized quantitatively. By doing meta-analysis of repeated research findings on the same topic, we can quantify how often the proclaimed confidence intervals actually end up containing the true mean value that emerges from all the evidence combined. A recent meta-analysis of 512 published meta-analyses in the fields of ecology and evolution (Yang *et al*. 2025) reported high levels of heterogeneity in research findings. As we will show, this suggests that, on average, only 45% of the time do the proclaimed confidence intervals contain the true value.

In the present study, we will explain how heterogeneity in research findings is typically quantified (using the statistical measure of *I*^*2*^) and how this measure is related to the proportion of CIs that should contain the true mean (here, the 45% mentioned above). We then explain how the discrepancy between 95% and 45% confidence can be understood and interpreted as the very expectable result of components of error that we hitherto have ignored in our statistical calculations. Conventional statistical practice, which systematically ignores certain sources of error (Simmonds *et al*. 2024), is bound to yield overconfident results that will often fail to replicate (Yarkoni 2022). Our review aims at not just raising awareness of neglected components of error, but also at enabling researchers to quantify, partly eliminate, and account for these components of error. While a range of sophisticated modelling tools have been developed that allow the specialist to incorporate particular additional source of error (see Simmonds *et al*. 2024 for an excellent review), we aim for a simple and easy-to-implement approach that, if desired, can cover *all* sources of error that may cause heterogeneity in study results. We explain how to calculate extended confidence intervals (CI_ext_) that should be able to keep what they promise, namely 95% confidence for a reasonably large inference space (*sensu* Anderson & McLean 1974; McLean *et al*. 1991). We also scrutinize the reasons why multiple analysts of the same dataset often disagree in their conclusions (Botvinik-Nezer *et al*. 2020; Breznau *et al*. 2022; Gould *et al*. 2025; Huntington-Klein *et al*. 2021; Schweinsberg *et al*. 2021; Silberzahn *et al*. 2018) and we show how to summarize multiple analyses of the same data set. We recommend the tools of “directed acyclic graphs” (DAGs, Borger & Ramesh 2025; Cinelli *et al*. 2024; Wysocki *et al*. 2022) and “specification-curve analysis” (Simonsohn *et al*. 2020) for settling disagreement on analysis decisions, and we highlight the risk of “comparing apples and oranges” in meta-analysis. We hence also call for caution in estimating and interpreting heterogeneity (*I*^*2*^).

### Three sources of error in individual studies

When carrying out a study, we typically go through the three phases of (A) design, (B) data collection, and (C) data analysis (Holzmeister *et al*. 2024). The decisions that are taken during each of these phases introduce some amount of uncertainty about our study’s conclusions (see **Fig. 1**). This uncertainty is about whether the choices that we took are actually the most representative choices for the broader research question at hand (Yarkoni 2022). Conventionally, yet problematically, only one of these three components of error in **Fig. 1**, namely the estimated sampling noise, is being used for the calculation of *P*-values and confidence intervals, while implicitly treating the other two factors as invariant. Variation that lies in these latter sources of uncertainty is, at best, discussed in the context of a sensitivity analysis (Thabane *et al*. 2013; Young & Holsteen 2017), but these sources of error are not (yet) part of conventional statistical analysis — a situation that could be changed.

**Figure 1.**
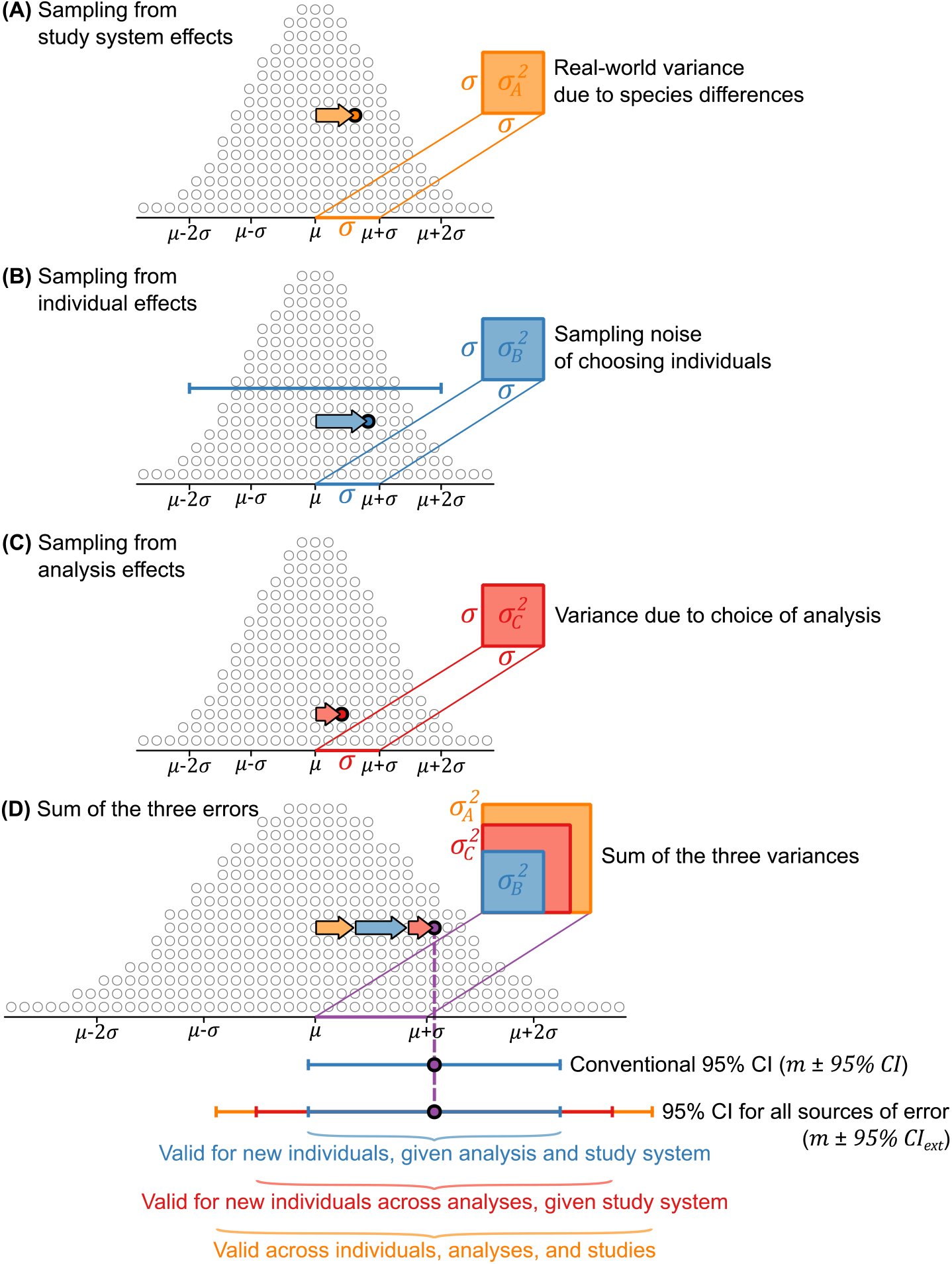
An illustration of the three sources of variation contributing to the total variation between studies. When choosing a study system (**A**), a set of study individuals (**B**), and a way of data analysis (**C**) we haphazardly sample from distributions of possible effects on the outcome variable. These populations of possible effects at each step are represented by piles of open circles, one of which is chosen in a particular study. The three sampling effects (orange, blue, and red arrows) will add up to a sum of effects (in **D**) that varies more widely than each of the three single components. In our example, all three components of sampling error have the same direction (which need not be the case), adding up to an empirical estimate *m* (dashed purple line). The orange, blue, and red squares illustrate the sampling variances (the square of the respective standard deviation, *σ*), which add up to the total sampling variance (three-coloured square), which determines the width of the distribution in (**D**). The conventional 95% CI (in blue) is based on the distribution in (**B**) only, and, in (**D**), is not sufficiently wide to ensure that the true population mean of *µ* has a 95% probability of falling within the blue interval. In contrast, the extended CI (orange) in (**D**) will contain *µ* in 95% of all cases. As indicated, such extended CIs have a wider range of interpretational validity.

A. During study design, we may decide to study a certain population of a certain species (**Fig. 1A**) in order to address a more general question (e.g. “does treatment *X* prolong the lifespan of animals?”). By taking this choice (rather than picking a different population or a different species), we have basically drawn a sample from an imaginary (and idealized) Gaussian distribution of true population-specific effects, which we use as a simple approximation for illustrative purposes. For instance, populations in which the treatment strongly prolongs lifespan will be on the far-right end of this distribution, while populations in which the treatment has little or even opposite effect on lifespan will be in the centre or on the left end, respectively. Hence, we sample from a pool of true effects that are intrinsic to each study system, and we are uncertain whether our choice of population is representative for animals in general (close to the mean of the distribution). There may be additional choices of study design (e.g. the implementation of treatment *X*), that feed into the same pool of error variance in design (but for simplicity, we here refrain from splitting this variance into several subcomponents). The magnitude of this variance in study system (or study design) effects 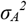 (i.e. the width of the Gaussian distribution in **Fig. 1A**) is initially unknown to us until multiple estimates become available for meta-analysis.
B. During data collection, we choose a subset of subjects (e.g. individuals) from the whole population in order to examine them (**Fig. 1B**). By quantifying the variability between subjects, we estimate the sampling noise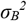, i.e. the uncertainty that comes from drawing these rather than an alternative set of subjects from the population. In practice, conventional 95% CIs are calculated based on this sampling noise (Brunswik 1949). Again, we may happen to draw a set of individuals whose sensitivity to the treatment lies close to the population mean *µ* or further away from it. Note that the same logic of assessing sampling noise applies in the situation where a given set of individuals is randomly assigned to treatment groups.
C. During data analysis, we typically select one particular way of estimating the effect of interest (effect of treatment on lifespan), from an entire pool of alternative options (aka “the garden of forking paths”; Gelman & Loken 2013) that would yield similar or different values for the effect of interest (**Fig. 1C**). This variation in analysis outcomes 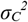 can be investigated using specification curves (Simonsohn *et al*. 2020) or multiverse analyses (Cantone & Tomaselli 2024; Steegen *et al*. 2016), or it may come to light when the statistical analyses are repeated independently by different researchers (Botvinik-Nezer *et al*. 2020; Breznau *et al*. 2022; Gould *et al*. 2025; Huntington-Klein *et al*. 2021; Schweinsberg *et al*. 2021; Silberzahn *et al*. 2018), who inevitably make different choices in the process of analysing the data (e.g. treatment of outliers, data transformation, inclusion of fixed and random effects, model selection procedures). This variance in outcomes arising from analysis decisions is also widely known as “model uncertainty” (Young & Holsteen 2017). The variance 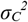 could be further subdivided (variance arising from alternative analyses that are fully legitimate versus from analyses that are simply incorrect and not defendable; Del Giudice & Gangestad 2021), but for now we treat “legitimate uncertainty” and “statistical incompetence” jointly as one source of variance.

Taking all three sources of variation together (choice of study system and methods, individuals, way of analysis), we end up with a much broader expected distribution of outcomes (see **Fig. 1D**) compared to the sampling noise alone (**Fig. 1B**) that we conventionally use for the calculation of the uncertainty in our study outcome (i.e. the classical 95% CI). Hence, while we report the conventional 95% CIs derived from our statistical model (**Fig. 1B**), we underestimate the genuine real-world 95% CIs, which would also include 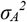 and 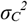and thus be derived from the sum of all three sources: 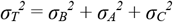. In this traditional way (**Fig. 1B**), we arrive at conclusions that are overconfident, if we think that they readily apply to a broader inference space — the real world that we are seeking to understand (Borm *et al*. 2009; Yarkoni 2022).

In our graphical example in **Fig. 1**, we chose all three sources of variance to be of equal size (red, blue, and orange squares), implying that the three idealized Gaussian distributions are of the same width. In that special case, and assuming the independence of the three errors, the sum of all three variances 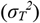 is exactly three times as large (the area of the three-coloured square in **Fig. 1D**), and the width of the distribution of the sum of the three errors in **Fig. 1D** is wider by a factor of 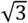 (i.e. the edge length of the three-coloured square, compared to the edge length of the blue square). With this in mind, we can now draw a range of confidence intervals around the study outcome *m*, that take the various sources of error into account. In blue (B), we indicate the conventional 95% CI, which refers only to the expected noise of sampling individuals from the study population. This 95% CI is hence valid for sampling new individuals from the same population and for analysing the data in exactly the same way (Gurevitch & Hedges 1999). If we then widen this 95% CI (here by a factor of 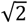 or 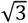, if we want to consider two (B + C, in red) or all three sources of variance (A + B + C, in orange)), we can obtain extended CIs that would also be valid across multiple ways of analysis (C), or even across multiple study system effects (A).

To implement this concept of extended CIs in practice, we require information on the width of the distributions included in the chosen inference space (e.g. 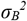 alone, 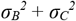 or 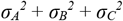), which we can obtain from meta-analyses once multiple estimates are available from several study systems 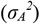or from multiple analyses of a given dataset 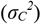. For most empirical studies it will be sufficient to account for the latter (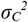, besides 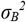 , of course), but keeping the former 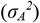 in mind is helpful for being clear about the legitimate inference space (*sensu* Anderson & McLean 1974; McLean *et al*. 1991).

Before we proceed with quantification of error components and confidence intervals, a brief note of justification may be required. We acknowledge that the three listed sources of error are of such different nature, that our approach of summing them up may be somewhat bewildering for a serious statistician. For instance, variation caused by analysis decisions has been described as “possibilistic” rather than as “probabilistic” (Hall *et al*. 2022), and we agree that calculating the average outcome of a large number of badly done analyses will typically not help with getting closer to the truth (Del Giudice & Gangestad 2021). However, from the point of view of an empirical researcher, there is a need for pragmatism. After discarding all bad choices of methods, it appears sensible to trust in the central tendency of results averaged across a large number of sensible choices, and to settle for the most robustly supported average conclusion. Our approach is hence meant to serve those who are looking for a simple and pragmatic solution to dealing with multiple sources of error.

### Quantifying these errors with meta-analysis

A so-called random-effects meta-analysis can be used to learn more about the magnitude of these errors. For those readers who have never conducted such an analysis, we encourage them to gain some hands-on experience using our most basic tutorial in the **Supplementary Materials (Part 4)** and to explore the work by Harrison (2011). Whenever one has multiple estimates of an effect (*m*_*i*_, where *i* refers to the *i*th study), and each estimate comes with a measure of uncertainty (standard error SE or confidence interval CI), one can not only derive a meta-analytic mean estimate with its own CI, but one can also test to what extent the entire set of estimates is in agreement or disagreement. The latter is done by quantifying heterogeneity. Speaking simplistically, the analysis examines whether the effects (*m*_*i*_) actually vary more or less than expected from their standard errors. As explained above, these standard errors are based on conventional statistics and solely reflect the uncertainty due to sampling noise in **Fig. 1B**. If we now find that the scatter in estimated effects (*m*_*i*_) exceeds the expectations from sampling noise B alone, we infer that there is additional variance from A and/or C. This amount of additional variance (the sum of the red and orange squares in **Fig. 1**) is commonly referred to as heterogeneity variance τ (Olsson-Collentine *et al*. 2020). As the magnitude of τ depends on the scale on which the effect sizes *m*_*i*_ are measured, it is more common to quantify heterogeneity in the form of *I*^*2*^, which describes the proportion of the total variation in study estimates (i.e. the three-coloured square in **Fig. 1**) that is due to heterogeneity (red plus orange squares) (Higgins & Thompson 2002; Higgins *et al*. 2003). Specifically, 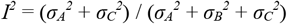 which is equivalent to subtracting the sampling noise from the total variance and dividing this by the total variance 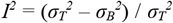. Note that, at this point, there is no way of distinguishing between the variances 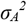 and 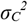, i.e. they can only be assessed jointly relative to the magnitude of variance 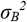.

The magnitude of *I*^*2*^ is not directly influenced by the number of studies included in a meta-analysis, but it strongly depends on the sample sizes used in the included studies (Rücker *et al*. 2008). This is because large samples help with averaging out the error, such that the sampling noise of each included study (blue square in **Fig. 1B**) strongly declines with increasing sample size. And when the sampling noise can be kept small, the heterogeneity will appear to be relatively larger (i.e. the smaller the blue square, the greater the relative contributions of the orange and red squares).

To illustrate this with some useful examples, an *I*^*2*^ = 0.67 would correspond to the situation in **Fig. 1** (with 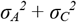 jointly making up two thirds of the total error), and *I*^*2*^ = 0.9 would imply that variances 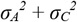 jointly are nine times larger than variance 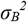. If the test for heterogeneity yields *I*^*2*^ = 0, then there appears to be no variance due to study system effects 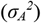 and the choice of analysis 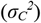, because the spread in the effect estimates *m*_*i*_ happens to be equal or less than expected from their standard errors. While *I*^*2*^ is bound between 0 and 1, the special situation of *I*^*2*^ = 0 can be further examined using tests for the statistical significance of *I*^*2*^. If one obtains *I*^*2*^ = 0 and *P* = 0.50, then the spread in estimates *m*_*i*_ is exactly equal to the expectation from sampling noise, while *P*-values above 0.5 indicate less spread in estimates than expected by chance. In an extreme case, if one obtains *I*^*2*^ = 0 and *P* = 0.99, then the spread in estimates *m*_*i*_ is apparently much less than expected by chance alone, a situation that might sometimes be suggestive of data fabrication (e.g. when the agreement in results all coming from the same author appears “too good to be true”; Francis 2012). Most frequently, however, we are confronted with a situation where *I*^*2*^ is rather large, for which there may be three possible explanations, that we will go through next.

### Main reasons for high levels of heterogeneity

#### (1) Heterogeneity due to differences in study systems

Whenever meta-analyses summarize studies that come from a wide range of study systems, it may seem most plausible to explain heterogeneity in effect sizes (*I*^*2*^ > 0) as a result of study system differences (e.g. Gurevitch & Hedges 1999), such as species differences in their sensitivity to the treatment (**Fig. 1A**). This often-made proposition, however, can also be tested systematically, for instance by adding species identity as a random effect (if there are repeated effect size estimates for the same species). In some cases, species identity is indeed an important factor (e.g. Rios Moura *et al*. 2021), and it may be interesting to examine the general role of phylogenetic inertia in such cases (for a tutorial see Mizuno *et al*. 2026). In other studies, species identity does not explain much of the observed variation in effect sizes (e.g. Cally *et al*. 2019; de Boer *et al*. 2021; Dougherty 2021; Pollo *et al*. 2025; Wang *et al*. 2019). Some authors have interpreted such lack of consistent differences between species optimistically as implying that the result of the meta-analysis is universally valid for all species (e.g. Pollo *et al*. 2025), yet one might also interpret this more pessimistically as implying a near-zero repeatability of research findings at the species level across studies. If species differences are not the main reason for heterogeneity between studies, then this may suggest a larger role for hard-to-tackle environmental conditions or methods of treatment that may differ between studies.

#### (2) Heterogeneity due to the choice of analysis

Recent “multi-team studies” have quantified the variance due to choice-of-analysis 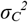 by having several teams of analysts independently analyse the same dataset and by quantifying the heterogeneity between their analysis outcomes (Botvinik-Nezer *et al*. 2020; Gould *et al*. 2025; Huntington-Klein *et al*. 2021; Schweinsberg *et al*. 2021; Silberzahn *et al*. 2018), which then mirrors our uncertainty about which model represents the correct way of analysis (Young & Holsteen 2017). Most of these studies have revealed an astonishingly large amount of heterogeneity originating from the “garden of forking paths” that consists of supposedly arbitrary analysis decisions (Gelman & Loken 2013). These studies seem to suggest that the error due to the “choice of analysis” (**Fig. 1C**) alone can cause dramatic heterogeneity in outcomes. However, there is also the view (Auspurg & Brüderl 2024a, b; Del Giudice & Gangestad 2021) that much of this apparent heterogeneity may arise from plainly incompetent data analysis by some contributing analysts and from meta-analysts summarizing effect size measures that are simply not comparable (the problem of “comparing apples and oranges”). We will come back to this controversy towards the end of our review, explaining why we suspect that truly arbitrary analysis variants (fully legitimate alternatives) typically introduce only modest amounts of heterogeneity in outcomes.

In meta-analyses that summarize published studies, the effects of choice-of-analysis could also introduce considerable heterogeneity, if a fraction of the published studies engaged in “*P*-hacking” via a targeted search for the way-of-analysis that yielded the most-extreme and typically significant outcomes (Simmons *et al*. 2011). To illustrate that *P*-hacking can induce heterogeneity, we here use a simple simulation. In each of 10,000 simulated datasets with a true effect size of zero (see **Supplementary Material Part 2**), we selected the most significant result among 128 alternative analyses (7 binary decisions allowing for 2^7^ = 128 variants) and compared it to a randomly chosen model from the same set. This increased heterogeneity in effect sizes from *I*^*2*^ = 0.02 to *I*^*2*^ = 0.61 (with 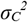 increasing nearly 50-fold). The choice to not publish non-significant findings (Rosenthal 1979) may also contribute to heterogeneity, especially in research fields where studies with strong positive *and* studies with strong negative effects are easier to publish than studies with non-significant effects. To quantify these effects empirically, it may be valuable to compare heterogeneity between meta-analyses of published studies and meta-analyses of unpublished data (e.g. Wang *et al*. 2019). Where unbiased data is not available, a range of techniques exists that may help with detecting and potentially eliminating publication bias (Nakagawa *et al*. 2022).

#### (3) Spurious heterogeneity due to underestimated sampling noise

An often-overlooked possible explanation for heterogeneity is that sampling noise may have been underestimated. We diagnose heterogeneity if the multiple study outcomes disagree with each other, and they do so confidently (based on their SEs or CIs). This can also happen when these SEs or CIs were simply too small, i.e. over-confident. The main reason for why SEs or CIs may often be too narrow is that analysts overestimate the effective sample size in their dataset by not recognizing important sources of non-independence that result in pseudo-replication (e.g. random effects, spatial and temporal autocorrelation) or overdispersion (see Bryan *et al*. 2021; Colegrave & Ruxton 2018; Forstmeier *et al*. 2017; Knief & Forstmeier 2021; Schielzeth & Forstmeier 2009; Voelkl *et al*. 2020; Yarkoni 2022), and this typically leads to confidence intervals around *m*_*i*_ that are too narrow. Since meta-analysis evaluates whether the variation in study outcomes *m*_*i*_ is larger than expected from sampling noise alone, any overestimation of the sample size (and corresponding underestimation of CI width) will lead to a meta-analysis finding heterogeneity in study outcomes that is spurious because it actually is a consequence of incorrect data analysis.

Interestingly, quantifying *I*^*2*^ in meta-analysis can also be used to identify and correct underestimated confidence intervals. As an example, Maldonado-Chaparro *et al*. (2021) carried out 10 replicates of the same aviary experiment (each aviary containing 28 birds), and entered the 10 resulting effect size estimates (Pearson *r*) with their standard errors into a meta-analysis. The finding of *I*^*2*^ = 0.9865 suggested that their initial CIs based on a widely-used randomization procedure (Butts 2008; for a more general treatment see Hart *et al*. 2022) had been too narrow by a factor of 8.6, showing that much wider CIs are needed for making inferences from one replicate aviary to the next, which is the minimal inference space that a study of general interest is supposed to cover. An intuitive way to evaluate the magnitude of reported SEs (or CIs) is to ask what sample size would be required to obtain uncertainty estimates of comparable size under independent sampling. For example, the SE of a Pearson correlation coefficient *r* can be approximated as 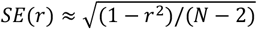, assuming that the *N* data points are independent (Sokal & Rohlf 1995). Rearranging this expression allows us to estimate the sample size implied by a given SE. In the aviary study, this calculation suggests that the narrow aviary-level CIs produced by the randomization procedure imply an effective sample size on the order of 1,740 independent observations per aviary (range: ∼1,070–2,580), which is an order of magnitude larger than even the raw input data (*N* = 196 for each aviary). In contrast, the 8.6-fold wider extended CIs (calculated as described in the next paragraph) correspond to a mean effective sample size of approximately 25.5 per aviary (range: ∼16–37), close to the actual number of 28 individuals per aviary. This suggests that, in this empirical study, the randomization procedure tended to overestimate the effective sample size by roughly 65–70-fold (pseudo-replication).

More generally, translating confidence intervals (or standard errors) into implied effective sample sizes may provide a useful diagnostic tool for evaluating whether these uncertainty estimates are compatible with the underlying sampling design. If they correspond to implausibly large effective sample sizes relative to the actual sample size, this indicates pseudo-replication or an overly narrow inference space. It may also provide a useful way for evaluating whether randomisation procedures adequately account for dependence structures and maintain appropriate type I error control (Hart *et al*. 2022; Winkler *et al*. 2015). In turn, heterogeneity-based approaches such as the extended confidence intervals proposed below may offer one possible way of correcting overconfident inference.

### Calculating extended confidence intervals

After having quantified heterogeneity in meta-analyses, there is the possibility of extending the conventional 95% CIs to cover all levels of uncertainty in individual studies. In other words, we can calculate extended CIs that are just wide enough, such that a new meta-analysis using the extended CIs would yield a heterogeneity estimate *I*^*2*^ of sharp zero (and a *P*-value = 0.50). For the extended 95% CIs we suggest the naming convention “95% CI_ext_” to clearly distinguish them from the conventional 95% CIs.

The conventional 95% CIs to our parameter estimate *m* are given by

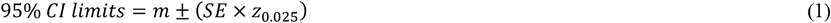

where *z*_0.025_ denotes the standard normal critical value (approximately 1.96 for a two-sided 95% interval), assuming that the estimator is approximately normally distributed (e.g. for large sample sizes). We calculate the standard error SE as 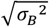 (Borenstein *et al*. 2021; see **Fig. 1** for the meaning of subscripts). We suggest to retain all conventional techniques for calculating *P*-values and CIs based on sampling noise alone, but to judge these at more stringent “heterogeneity-aware” thresholds. Thus, we need to extend the conventional 95% CIs to 95% CI_ext_ by increasing the standard error from 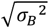 to 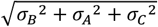 . This can be done by multiplying the conventional 95% CIs with 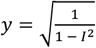 (we formally derive *y* in the **Supplementary Material Part 1**), which means that we obtain the extended 95% CIs as

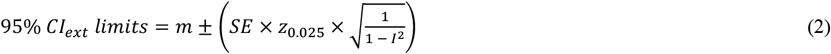

This formula was independently derived by Borm *et al*. (2009). While we assume a normal distribution and the use of a *z*-score, our calculations are generally valid for all cases where a normal approximation is appropriate (i.e. most common analyses with reasonable sample sizes and well-chosen statistics; Borm *et al*. 2009; IntHout *et al*. 2016). Throughout, variance components are estimated using a plug-in approach and treated as fixed quantities, such that the resulting intervals represent large-sample approximations.

Our proposed measure of CI_ext_ constitutes a more conservative confidence interval that acknowledges additional sources of uncertainty, quantified empirically using the *I*^*2*^ metric of heterogeneity. Note that these extended confidence intervals differ from so-called prediction intervals. While both approaches are informed by between-study heterogeneity, prediction intervals in meta-analysis leave the within-study standard errors unchanged and describe the dispersion of true effects across studies (corresponding to the distribution in **Fig. 1A**), and hence the range within which the true effect of a new study is expected to fall. They are therefore mainly determined by the between-study variance and, unlike confidence intervals for the mean effect, do not approach zero as the number of studies increases. Prediction intervals in primary studies, in turn, describe the variability of individual observations within a population, rather than the uncertainty in the population mean (Borenstein *et al*. 2021). In contrast, our extended confidence intervals quantify uncertainty in individual study estimates after accounting for additional sources of variation arising from study systems and analytical choices, and explicitly rescale the standard errors of individual studies to incorporate these additional sources of uncertainty.

Having defined extended confidence intervals, we can also calculate the corresponding *α*-thresholds (i.e. “significance”-thresholds) that would keep the type I error rate at the desired level of, say, 5%.

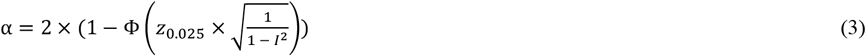

where, Φ(x) is the cumulative distribution function of the standard normal distribution.

### Limits of setting stricter *α*-thresholds for given *I*^*2*^ values

**Table 1 and Figure 2** provide an overview of how the key parameters in hypothesis testing (width of CI and critical *α*-threshold) would need to be adjusted in order to account for a given level of heterogeneity in study outcomes. For small and medium levels of heterogeneity (*I*^*2*^ < 0.6), these adjustments appear relatively minor, yet as we get to high levels of heterogeneity (e.g. *I*^*2*^ = 0.9), the adjustments become so extreme that they no longer appear sensible. For instance, Yang *et al*. (2025) reported that the median level of heterogeneity in meta-analyses in ecology and evolution was as high as *I*^*2*^ = 0.908. In this scenario, conventional 95% CIs would, on average, cover the true population parameter in only 45% of the cases — and hence the title of our paper (see **Table 1, Fig. 2A**). To bring this value up to the desired 95%, extended confidence intervals would have to be wider by a factor of 3.3-fold (**Table 1, Fig. 2B**). Accordingly, the critical *α*-threshold would have to be lowered from 0.05 to a sobering *α*_*ext*_ = 6.4 × 10^−11^ (**Table 1, Fig. 2C**). In most cases, such adjustment would do more harm than good for scientific progress. Firstly, such extreme *α*-thresholds would unlikely be helpful for overcoming the reliability crisis, because overly strict criteria might primarily incentivise bad scientific practice (Amrhein *et al*. 2019; Smaldino & McElreath 2016; Ware & Munafò 2015; Wasserstein *et al*. 2019). Secondly, and more importantly, we argue that *I*^*2*^-values of such magnitude should generally make us sceptical to begin with. As we will discuss towards the end of our review, we should check whether heterogeneity has been overestimated, and whether the meta-analysis suffers from the issue of “comparing apples and oranges”, which one could also paraphrase as the meta-analysis targeting an inference space that is larger than what is biologically meaningful and statistically sound.

**Table 1.** Derivation of the global type I error rate, the size of confidence intervals and extended *α* thresholds given heterogeneity *I*^*2*^. These derivations are identical to those in Borm *et al*. (2009). For orientation, we also provide heterogeneity estimates from published meta-analyses and multi-analyst studies. “*y*” refers to the correction factor by which the standard errors and CIs would have to be multiplied in order to be sufficiently wide to keep the type I error rate at 5%. “*z* score given *I*^*2*^” shows how the realized threshold for declaring effects as significant gets lower when *I*^*2*^ is high, leading to a high rate of false-positive findings (“Type I error rate given *I*^*2*^”). Accordingly, the proportion of CIs covering the true mean *µ* goes down, and these expected values correspond well with observed numbers of CIs that contain *µ*. “Extended *z* score” shows the critical threshold for declaring significance that should be used to keep type I errors at a 5% rate, and the “Extended *α* threshold” shows critical *P*-values that would be needed for keeping type I errors at 5%. Calculations are implemented in the **Supplementary Material Part 4.** The pnorm() function in R (R Core Team 2024) is the cumulative distribution function of the standard normal distribution Φ(x).

| Landmark | Reference | Heterogeneity<br>$I^2$ | Conventional 95% CIs | | | | | Extended 95% CIs | |
| --- | --- | --- | --- | --- | --- | --- | --- | --- | --- |
| | | | $y$ | $z$ score<br>given $I^2$ | Type I error rate<br>given $I^2$ | Expected<br>proportion of<br>conventional 95%<br>CIs cover $\mu$ given $I^2$ | Observed<br>proportion of<br>conventional 95%<br>CIs cover $\mu$ given $I^2$ | Extended<br>$z$ score | Extended $\alpha$ threshold |
| Calculation | | $\frac{\sigma_A^2 + \sigma_C^2}{\sigma_B^2 + \sigma_A^2 + \sigma_C^2}$ | $\sqrt{\frac{1}{1 - I^2}}$ | $\frac{1.96}{y}$ | $2 \times (1 - \Phi(\text{Z score}))$ | $1 - \text{Type I error rate}$ | | $1.96 \times y$ | $2 \times (1 - \Phi(\text{extended Z score}))$ |
| Zebra finch aviaries | Maldonado-Chaparro *et al.* 2021 | 0.987 | 8.607 | 0.228 | 0.820 | 0.180 | 0.200 | 16.869 | $7.6 \times 10^{-64}$ |
| Blue tits | Gould *et al.* 2025 | 0.977<sup>§</sup> | 6.545 | 0.299 | 0.765 | 0.235 | 0.198 | 12.828 | $1.1 \times 10^{-37}$ |
| Obs median Ecol & Evol: lnRR<sup>†</sup> | Yang *et al.* 2025 | 0.955 | 4.689 | 0.418 | 0.676 | 0.324 | | 9.191 | $3.9 \times 10^{-20}$ |
| Est mean Ecol & Evol | Senior *et al.* 2016 | 0.917 | 3.469 | 0.565 | 0.572 | 0.428 | | 6.799 | $1.1 \times 10^{-11}$ |
| Obs median Ecol & Evol | Yang *et al.* 2025 | 0.908 | 3.303 | 0.593 | 0.553 | 0.447 | | 6.474 | $9.5 \times 10^{-11}$ |
| Obs median Ecol & Evol: SMD<sup>†</sup> | Yang *et al.* 2025 | 0.892 | 3.037 | 0.645 | 0.519 | 0.481 | | 5.953 | $2.6 \times 10^{-9}$ |
| Obs median Ecol & Evol: $Z_r$<sup>†</sup> | Yang *et al.* 2025 | 0.868 | 2.757 | 0.711 | 0.477 | 0.523 | | 5.404 | $6.5 \times 10^{-8}$ |
| Est median Ecol & Evol | Senior *et al.* 2016 | 0.847 | 2.554 | 0.767 | 0.443 | 0.557 | | 5.006 | $5.6 \times 10^{-7}$ |
| Obs median Ecol & Evol | Senior *et al.* 2016 | 0.797 | 2.220 | 0.883 | 0.377 | 0.623 | | 4.351 | $1.4 \times 10^{-5}$ |
| *Eucalyptus* outliers removed | Gould *et al.* 2025 | 0.747<sup>§</sup> | 1.989 | 0.985 | 0.325 | 0.675 | 0.733 | 3.899 | $9.7 \times 10^{-5}$ |
| High | | 0.900 | 3.162 | 0.620 | 0.535 | 0.465 | | 6.198 | $5.7 \times 10^{-10}$ |
| High | | 0.800 | 2.236 | 0.877 | 0.381 | 0.619 | | 4.383 | $1.2 \times 10^{-5}$ |
| High | | 0.700 | 1.826 | 1.074 | 0.283 | 0.717 | | 3.578 | $3.5 \times 10^{-4}$ |
| Medium | | 0.600 | 1.581 | 1.240 | 0.215 | 0.785 | | 3.099 | $1.9 \times 10^{-3}$ |
| Medium | | 0.500 | 1.414 | 1.386 | 0.166 | 0.834 | | 2.772 | $5.6 \times 10^{-3}$ |
| Medium |  | 0.400 | 1.291 | 1.518 | 0.129 | 0.871 |  | 2.530 | 0.011 |
| Small |  | 0.300 | 1.195 | 1.640 | 0.101 | 0.899 |  | 2.343 | 0.019 |
| Small |  | 0.200 | 1.118 | 1.753 | 0.080 | 0.920 |  | 2.191 | 0.028 |
| Small |  | 0.100 | 1.054 | 1.859 | 0.063 | 0.937 |  | 2.066 | 0.039 |
| None | Magnusson 2014 | 0.000 | 1.000 | 1.960 | 0.050 | 0.950 | 0.950 | 1.960 | 0.050 |

<sup>†</sup> lnRR, SMD and Z<sub>r</sub> are different effect size measures summarized in Yang *et al.* (2025). lnRR: log response ratio, SMD: standardized mean difference, Z<sub>r</sub>: Fisher’s r-to-z transformed correlation coefficient.
<sup>§</sup> these are estimates of $I^2_{MAMA}$ calculated from values of $I^2$ \* reported in Table 2 of Gould *et al.* (2025).

**Figure 2.**
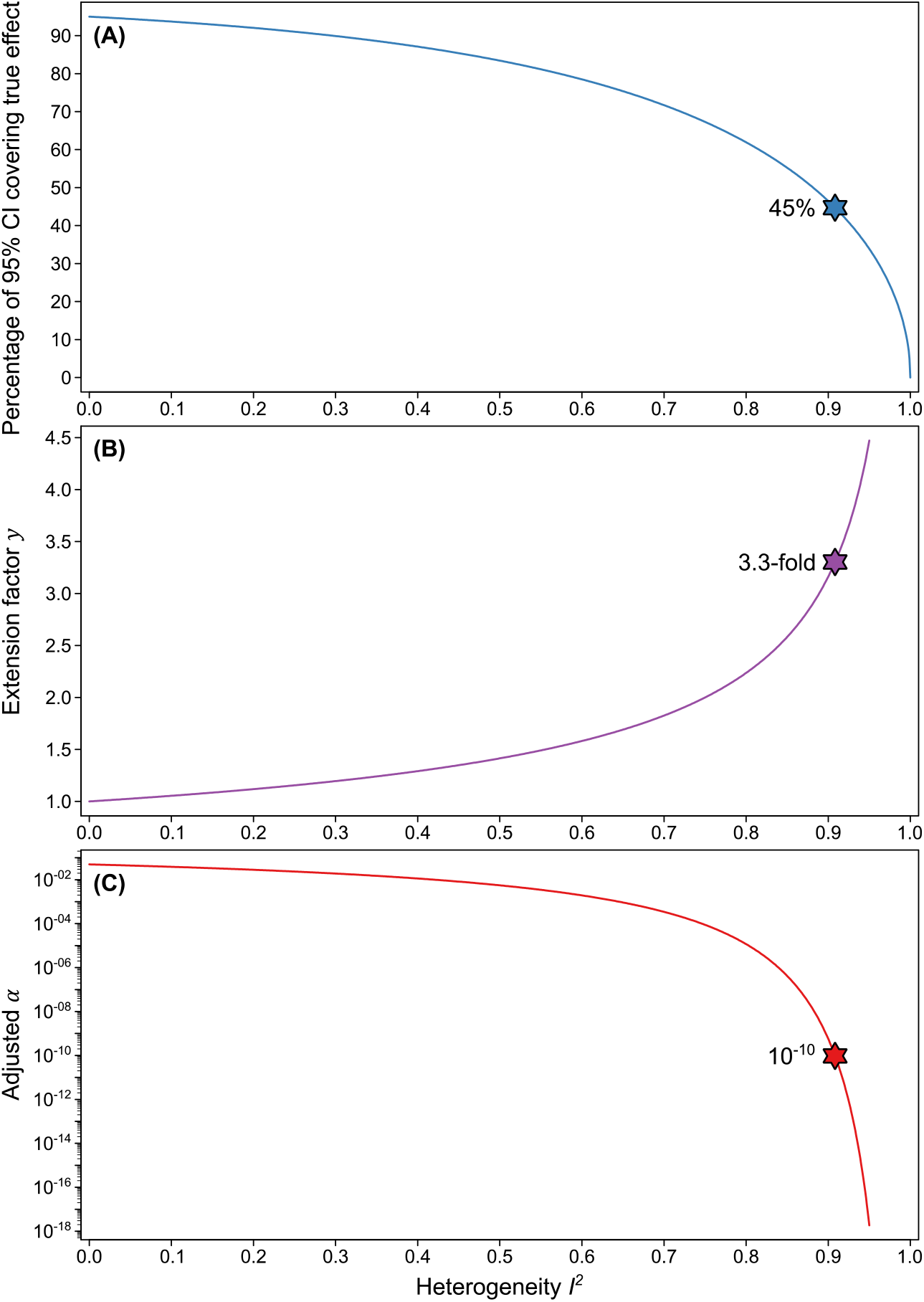
Relationship between heterogeneity *I*^*2*^ and (**A**) the percentage of conventional 95% CIs covering the true population parameter, (**B**) the extension factor *y* required to convert conventional to extended 95% CIs, and (**C**) the adjusted *α*-threshold for the extended 95% CIs. At *I*^*2*^ = 0, the conventional 95% CIs are equal to the extended 95% CIs and the *α*-threshold does not need to be adjusted (i.e., it is at 0.05). As soon as there is heterogeneity between studies due to study system effects or the choice of analysis, the confidence intervals need to be adjusted by *y* to continue to cover 95% of the true population parameter, resulting in lower *α*-thresholds. Stars indicate the conventional 95% Cis covering the true population parameter, the extension factor *y*, and the adjusted *α*-threshold given a heterogeneity *I*^*2*^ of 0.908 as reported by Yang *et al*. (2025).

### Merging multiple versions of analysis by use of MAMA

**Figure 3** illustrates how we can, in principle, eliminate most of the uncertainty about which choice of data analysis is the most representative among numerous reasonable alternative ways of analysis. We here use the “*Eucalyptus* dataset” of Gould *et al*. (2025) as an example, in which a large number of analysis teams examined the very same dataset to estimate the magnitude of a specific effect. From the 75 effect size estimates, we can see that the choices taken by some teams appear to support a significant negative effect size, the choices taken by others suggest a significant positive effect size, yet most choices yield intermediate effects that are close to zero. If we assume that all of these analyses are sensible and valid, we can summarize all these alternative choices into a meta-analytic mean value (the dashed purple line in **Fig. 3**), in the hope that this will eliminate most of the uncertainty, and will get us closer to the most-representative way of analysis (close to *µ* in **Fig. 1C**). Of course, it is fair and necessary to question on a case-to-case basis whether some of the 75 analyses are misleading and should be excluded from contributing to the meta-analytic mean value (see e.g. Del Giudice & Gangestad 2021). Here, we simply followed Gould *et al*. (2025) in excluding the four most extreme estimates from an initial set of 79 analyses.

**Figure 3.**
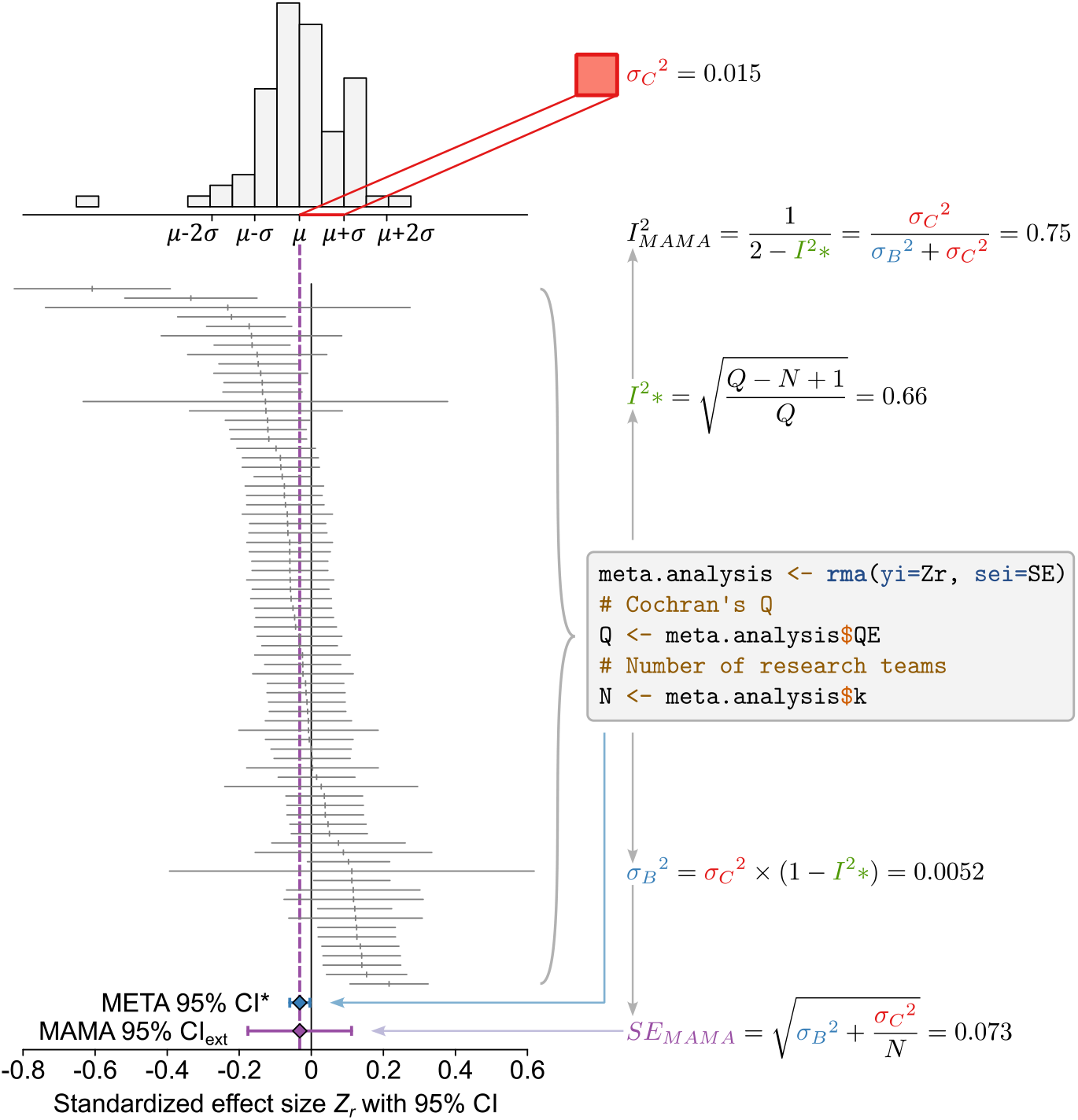
Example of a many-analyses meta-analysis (MAMA) based on the “*Eucalyptus* dataset” of Gould *et al*. (2025). The forest plot shows the 75 estimated effect sizes (and their 95% CIs) obtained from different analysts (after removal of 4 outlier results), all of which had been asked to examine the same research question by analyzing the same dataset (Gould *et al*. 2025). The histogram at the top shows the variance in outcome measures 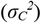 in analogy to **Figure 1.** The diversity in analysis outcomes can be summarized in a meta-analytical mean estimate (dashed purple line), accompanied by the appropriate MAMA 95% CI_ext_ (in purple), which is slightly wider than the typical (median) CI from single analysts that are based on 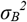 only. Note that conventional meta-analysis of such a dataset would lead to a highly misleading “META 95% CI*” (in blue), which would only be valid if these were 75 independent studies with independent data (rather than 75 analyses of the same data). Moreover, conventional meta-analysis would conclude that 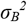 is responsible for approximately 1/3 of 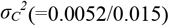, yielding the incorrect measure of *I*^*2*^*\** = 0.66, while in fact, 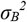 is not part of 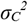, but comes on top of it (yielding *I*^*2*^_*MAMA*_ = 0.75). Note that 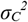 makes only a small contribution to the size of MAMA 95% CI_ext_, because the error is averaged out across the *N* = 75 analyses (dividing the error variance by *N* in the calculation of SE_MAMA_). Also note that *I*^*2*^-values reported in Gould *et al*. (2025) correspond to our *I*^*2*^*\**. They did not use this label, but explicitly recognized the need for new definitions in the present context (of summarizing many analyses of the same data set).

While sending all our datasets to a large number of colleagues for obtaining multiple versions of analysis may be impractical, we can also explore different analysis choices by undertaking a so-called “multiverse analysis” (Cantone & Tomaselli 2024; Steegen *et al*. 2016). For readers who are not yet familiar with this approach, we recommend the very intuitive tutorial by Philipp Masur (Masur 2020), besides the more in-depth treatments by Cantone & Tomaselli (2024) and Rohrer *et al*. (2026). In brief, there is software (Masur & Scharkow 2020) that allows to go through all “reasonable” versions of data analysis. The output of such analysis is summarized in a curve of effect sizes that are sorted by their magnitude (like in **Fig. 3**). If each entry to that curve gets labelled with a dot-plot indicating the underlying analysis decisions, it becomes easy to identify which choices are most influential in yielding large versus small outcome measures (aka “specification-curve analysis”; Simonsohn *et al*. 2020). In practice, it may often turn out that highly influential analysis decisions are actually those which fundamentally alter the research question that is being asked (Del Giudice & Gangestad 2021). This does not necessarily mean that such analyses are wrong. Rather, empirical research questions are often broader than any single statistical model, and different model specifications may represent different, yet biologically defensible, operationalisations of the same broad question. For example, controlling for a certain covariate, or not, can make a drastic difference to the outcome (and the biological question) depending on how the covariate is related to the predictor of interest and to the dependent variable (Cinelli *et al*. 2024; Wysocki *et al*. 2022). Typically, for understanding the total causal effect of a treatment on a dependent variable, you want to control for covariates that are confounders (or confound-blockers; considered as “good controls”), but you must not control for mediators, proxies, or colliders (considered as “bad controls”; Cinelli *et al*. 2024; Wysocki *et al*. 2022). Models that contain such “bad controls”, in most cases, address a very peculiar question and not the relevant question of the total effect of treatment on the dependent variable. Averaging across such different choices (e.g. controlling for a mediator or not), and hence across different research questions, may end up being the “comparison of apples and oranges”, which we will illustrate with a specific example further below. We therefore fully agree with Del Giudice & Gangestad (2021) in recommending to limit the multiverse analysis to only those analysis choices that do not alter the underlying research question. Yet, one may also argue for including numerous or essentially all sensible ways of analysis, such that we can rightfully generalize across a broad inference space.

### Calculating an appropriate MAMA CI_ext_

If we want to summarize the outcomes of multiple analyses of the same dataset using meta-analysis (like in **Fig. 3**), we have to realize that it is not appropriate to simply use the software-inbuilt formulas that refer to a different context, namely the meta-analysis of independent studies that differ in study design, data collection, and data analysis. In the following, we label all those parameters with an asterisk (*) that were calculated from an inappropriate formula, in order to make them distinguishable from the true values. First, the CI* of the meta-analytic mean value (shown in blue in **Fig. 3**) is much too small, because it is based on the wrong assumption that **Fig. 3** summarizes 75 independent studies (each of which would represent an independent draw from 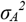 and 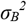 and 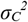), rather than 75 analyses of the same dataset (75 draws from 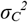 only). Second, the inbuilt formulas yield a measure of *I*^*2*^*\**, which incorrectly assumes that part of the variation in analysis outcomes 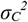 was caused by sampling noise 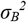, while in fact all analyses shared almost the exact same sampling noise. In a conventional meta-analysis, the total variance in study outcomes 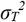 is the sum of three sources of error 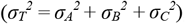, while here, all the variance between study outcomes originates from 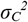 only (see the histogram in **Fig. 3**), 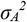 equals zero, and the sampling noise 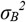 (being a matter of the sample size of the dataset, and reflected by the CIs provided by analysts) still requires to be considered in addition. With the examples of 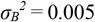 and 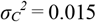 (from **Fig. 3**), conventional analysis assumes that 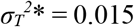, being the sum of two components, namely 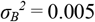 and 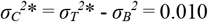, yielding 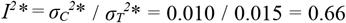. However, in fact, 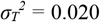, being the sum of 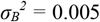 and 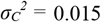, yielding 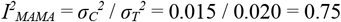

Beyond this example, we can generally compute *I*^*2*^_*MAMA*_ as

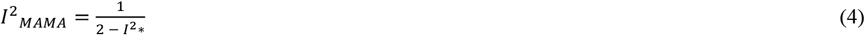

(for a formal derivation see the **Supplementary Material Part 1**). If we use the software-inbuilt *I*^*2*^*\** for calculating *I*^*2*^_*MAMA*_, this would restrict *I*^*2*^_*MAMA*_ to the interval between 0.5 and 1 (which we can see by plugging in the lower (*I*^*2*^*\** = 0) and upper (*I*^*2*^*\** = 1) bound into equation (4). Yet, *I*^*2*^_*MAMA*_ should reach from 0 to 1, because 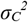can be smaller than 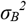. Theoretically it can even approach zero, if all analysts followed the same analysis path, in which case *I*^*2*^_*MAMA*_ = 0 (equivalent to *I*^*2*^*\** = 0, *P* → 1). We can make use of the fact that *I*^*2*^*\** can also be derived from Cochran’s *Q* (Higgins & Thompson 2002), and in fact it commonly is (Higgins *et al*. 2003). In that case *I*^*2*^*\** can also take negative values, leading to *I*^*2*^_*MAMA*_ approaching zero, if the differences between analyses is small (see the **Supplementary Material Part 1** for details and **Fig. S1**). We provide R code for calculating *I*^*2*^_*MAMA*_ in the **Supplementary Material Part 4**.

Using a similar line of reasoning, which is formally laid out in the **Supplementary Material Part 1**, we calculate the 95% CI_ext_ for a many-analyses meta-analysis from SE_MAMA_ as

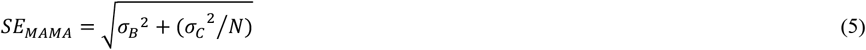

where *N* is the number of analyses summarized in the MAMA. Accordingly,

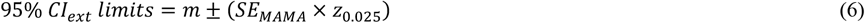

This formulation treats 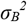 as the typical within-analysis sampling variance across analyses. We show how to derive 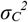 and 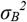 , empirically validate the procedure of dividing 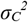by *N*, and provide R code in the **Supplementary Material Part 4**.

While all this may seem like an elegant solution to a major problem, the upcoming final part of our article actually suggests that there is often not much variance due to arbitrary analysis decisions 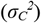 that can and should be eliminated. Instead, it appears that much of the variance in 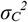 arises from analysis decisions that shift the underlying biological question. A different question will often yield a different answer, and how precisely the research question is formulated should usually not be glossed over, as it risks averaging away the essential nuances.

In the following final section, our perspective will take a rather dramatic turn. We raise the question of whether meta-analyses might frequently overestimate heterogeneity, partly for technical and partly for biological reasons. Using a specific example (the “blue tit dataset” from Gould *et al*. 2025), we also challenge the notion that arbitrary analysis decisions often exert a dramatic effect on the conclusions. What this example instead illustrates is that (1) there are many ways of making analytical mistakes, and that (2) two distinct experimental approaches to the same research question can yield drastically different answers.

### Do meta-analyses frequently overestimate the magnitude of the problem?

We raised the question of whether conventional 95% CIs may only be worth 45% confidence, which was based on the median *I*^*2*^ of 0.908 reported by Yang *et al*. (2025). However, the estimate of only 45% deserved confidence might be overly pessimistic. Some meta-analyses may have overestimated heterogeneity by overestimating the effective sample size of empirical studies (for instance, by basing study CIs on the number of individuals rather than the true number of independent replicates; e.g. Cally *et al*. 2019). In such cases, a meta-analysis might work with smaller CIs than those reported in the original literature. Further, heterogeneity in outcomes may be inflated, if the conversion of parameter estimates to effect sizes fails, so that they are not on the same scale (Auspurg & Brüderl 2024a, b). Similarly, heterogeneity may be inflated when the meta-analysis includes effect sizes expressed as (log) odds ratios, instead of collapsible effect size estimates like risk ratios or the average marginal effects from logistic regression (Greenland *et al*. 1999; Mood 2009). Finally, if meta-analysts regularly decide to summarize outcome measures that are biologically not comparable (“comparing apples and oranges”, see the example below), then this would also contribute to the high levels of heterogeneity observed by Yang *et al*. (2025). A systematic analysis of these issues would be much needed before making any calls on whether heterogeneity is often over-estimated.

### How big is the effect of truly arbitrary analysis decisions?

To explore the issue of choice-of-analysis more systematically, we carried out analyses on simulated datasets which allowed for 7 different binary analysis decisions, meaning that any given dataset could be analysed in 2^7^ = 128 different ways (see **Supplementary Materials Part 2**). We found that the amount of variance introduced by these arbitrary analysis decisions was usually quite modest (median 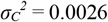 and 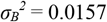), but in rare cases could reach considerably high levels (**Fig. S2**). Such a median level of variance due to analysis decisions would lead to heterogeneity of *I*^*2*^_*MAMA*_ = 0.14 and would call for CI_ext_ being wider by a factor of 1.08 than the conventional CI based on sampling noise alone, and for a heterogeneity-aware critical *α* of 0.034. More extensive simulation studies are needed to assess the generality of this result, extending it also to non-normally distributed data. There is also a need to find out whether such a rule of thumb (making CIs 8% wider) is usually sufficient to account for uncertainty in modelling decisions, or whether each study should do its own assessment of the consequences of uncertainty in the models that are being used. In any case, our result that arbitrary analysis decisions cause only little heterogeneity appears to stand in stark contrast to what many of us concluded from multi-team studies (Botvinik-Nezer *et al*. 2020; Breznau *et al*. 2022; Gould *et al*. 2025; Huntington-Klein *et al*. 2021; Schweinsberg *et al*. 2021; Silberzahn *et al*. 2018), which mostly found that analyst teams differed drastically in their conclusions (i.e. in their effect size estimates). We therefore examine a specific example below.

### Revisiting the blue tit dataset — a case of remarkable disagreement among analysts

According to **Table 1**, one of the most spectacular cases of heterogeneity due to “choice of analysis” is the “blue tit dataset” of Gould *et al*. (2025). In this multi-team study, 131 different analyses of the same dataset yielded results that were discordant to such an extent (*I*^*2*^_*MAMA*_ = 0.977), that our approach would recommend extending the CIs of single analyses by a mind-blowing factor of 6.5 (see **Table 1**) in order to encompass all the uncertainty about how to correctly analyse this data. This would imply that the very large and highly significant effect size estimate that one of us contributed to this study (Z_r_ = -0.637 ± 0.052 SE) and reported as falling |*z*| = 12.3 standard errors away from zero, would now fall just short of the extended critical *z*-score (|*z*| = 12.8, **Table 1**) and therefore be no longer considered significantly different from zero, simply because many colleagues apparently disagree with there being an effect.

To better understand what can lie behind such major disagreement, we here present some targeted versions of analysis that may help clarify part of the debate. We do this by following the advice of Del Giudice & Gangestad (2021), namely to distinguish between equivalent (E) and non-equivalent (N) analysis decisions. In case of type (E) decisions, we have no *a priori* conceptual reasons that would clearly favour one version of analysis over the other, we can hence speak of truly arbitrary choices between equally valid options (“principled equivalence”). In contrast, principled non-equivalence means that we know before-hand that one analysis variant is inferior to the other (e.g. low vs. high validity of a trait), or that the two variants of analysis are actually testing two conceptually different hypotheses (e.g. inclusion of a covariate that results in overcontrol-bias; Arif & MacNeil 2023). In between those extremes of (E) and (N), will be cases of uncertainty (U) regarding equivalence.

For our re-analysis of the blue-tit data set we chose 6 models that differ in type (E) decisions regarding the treatment of “good controls”, missing data, outlier removal, and data transformation. As a drastic contrast, we chose 8 models that differ in type (N) decisions, namely the inclusion of covariates which can be referred to as “bad controls” (Cinelli *et al*. 2024). These include conditioning on a mediator (or descendants of a mediator; causing “overcontrol bias”), conditioning on alternative outcome measures, and conditioning on components of the predictor variable. We also created two models with type (U) decisions, for which equivalence may be debated (see **Supplementary Materials Part 3** for details).

Gould *et al*. (2025) provided data from a brood-size manipulation experiment in wild blue tits (*Cyanistes caeruleus*) breeding in nest boxes to multiple independent analysis teams. Each team was asked to answer the question: “To what extent is the growth of nestling blue tits influenced by competition with siblings?” This question was deliberately defined loosely, thereby allowing for multiple analytical pathways. The dataset contained information on 3,720 nestlings from 497 clutches. As an experimental treatment, nestlings had been partly swapped between broods, leading to net brood size reductions and enlargements, and roughly a quarter of the broods remained “unmanipulated”. Information was provided on nestling growth (per nestling: body mass at day 14, tarsus length at day 14) and nestling survival (per brood: post-treatment brood size, day 14 brood size, number of fledglings). In the present analysis, we narrow the focus and define *Net Manipulation* (the net change in number of chicks) as our continuous treatment variable (ranging from -4 to +4). We believe that in an experimental context, the primary goal is to estimate the effect of the experimental treatment, rather than natural variation. We consider *Net Manipulation* therefore the most direct operationalization of what is meant by “competition with siblings” in the original question. We further use *Day 14 Chick Mass* as our primary outcome variable. Based on these decisions, we constructed a directed acyclic graph (DAG; see **Fig. S3**), which illustrates the causal pathways from brood size manipulation to chick body mass at day 14 (Borger & Ramesh 2025; Pearl 2009). We do acknowledge that slightly different versions of DAGs could be drawn and that this example represents a comparatively clear-cut case; in many empirical settings, distinctions between equivalent and non-equivalent analyses will be less clear, and part of the observed heterogeneity may reflect genuinely different but defensible ways of addressing a broader research question (see also Gould *et al*. 2025).

### Who is to blame for heterogeneity? The legitimate garden, incompetence, or diversity of fruits?

When focussing on the entire blue tit dataset (all broods, including those where brood size was enlarged, reduced, or left unmanipulated; left panel of **Fig. 4**) we see that models that differ only in type (E) decisions yield nearly identical effect size estimates (effect of the net brood-size manipulation on offspring body mass). In contrast, when changing the model non-equivalently (N), such that it deliberately contains a problematic mediator variable (e.g. the post-manipulation clutch size; see **Fig. S3**) or alternative outcome measures as a covariate, it is possible to weaken the otherwise clear treatment effect. Note that we designed the latter kind of models such that they blur the targeted hypothesis, in order to demonstrate possible effects of poorly designed models.

**Figure 4.**
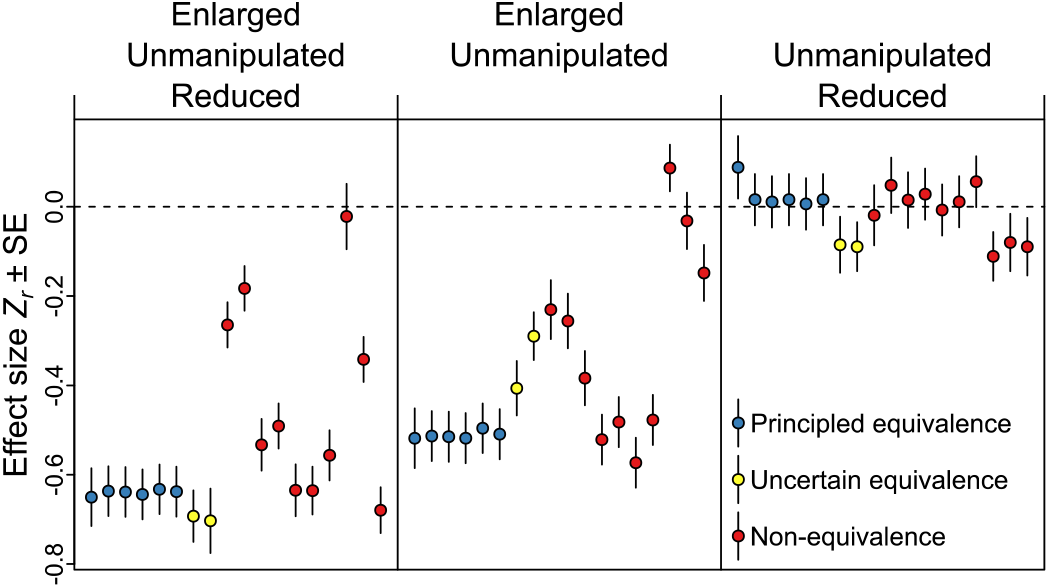
Effects of analytical decisions on estimated treatment effects in the “blue tit dataset” of Gould *et al*. (2025). Effect size estimates Z_r_ are Fisher-transformed effect sizes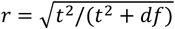of the net brood size manipulation (number of nestlings removed from or added to each nest) on offspring size (mass or tarsus length). The three panels illustrate the effect of different data selection decisions: left, all nests included; centre, enlarged and unmanipulated broods only; right, reduced and unmanipulated broods only. In each panel, blue dots represent six models based on analytical decisions that we consider truly arbitrary (reflecting principled equivalence *sensu* Del Giudice & Gangestad (2021)). The two yellow dots represent models where equivalence may be debated. The ten red dots correspond to models based on principled non-equivalence, where conditioning on a mediator, descendant of a mediator or alternative outcome measures alters the biological hypothesis being tested. Such models are intended to be non-sensible. For details on each model see **Supplementary Materials Part 3**, which also contains the results for using an alternative measure of effect size, which is better suited for biological interpretation (**Fig. S4**), as well as robustness checks based on alternative operationalizations of the outcome and treatment variables (**Figs. S5, S6**).

We then applied the same set of 18 models to two subsets of the data: one containing only enlarged and unmanipulated broods (centre panel of **Fig. 4**), and one containing only nests where brood size had been either reduced or left unmanipulated (right panel of **Fig. 4**). The most striking (post-hoc) observation here is that brood-size reductions appear to have no effect on offspring mass, while brood-size enlargements have clear negative effects on offspring mass.

In this example, we can see three clear patterns:

1. Analysis decisions that are truly arbitrary (within the legitimate garden of forking paths; “principled equivalence”) appear to have only minor effects on the analysis outcome (little scatter of blue dots within each of the three panels of **Fig. 4**), thereby confirming our earlier conclusion (**Fig. S2**) also for this particular data set.
2. Analyses that are purposefully incompetently done can make even large true effects disappear, and thereby induce much heterogeneity in outcomes (wide scatter of red dots in the left and central panels in **Fig. 4**).
3. Speaking with the benefit of hindsight, when meta-analysts jointly evaluate the effects of brood-size enlargements and reductions within one analysis, they may be “comparing apples and oranges” (large difference between panels in **Fig. 4**).

To confirm the robustness of these patterns, we repeated this series of analyses also for a different effect size measure (**Supplementary Materials Part 3 Fig. S4**), for a different outcome variable (tarsus length; **Fig. S5**), and for an alternative classification of the treatment variable (as two-level factor; **Fig. S6**), and we obtained qualitatively the same results.

While this extreme example demonstrates that analyst incompetence and the comparison of “apples and oranges” can have very large effects, there is probably also a role for decisions of uncertain equivalence (U) and a need to discuss such analysis decisions on a case-to-case basis. In the end, it always comes down to defining an intended inference space, which might sometimes also comprise a wider diversity of fruits (apples and oranges) (Yarkoni 2022).

### Practical recommendations

In order to substantiate the main conclusions of an empirical study more reliably, we encourage data analysts to explore the (dataset-specific) garden of forking paths of arbitrary analysis decisions by means of multiverse analysis (Cantone & Tomaselli 2024; Simonsohn *et al*. 2020; Steegen *et al*. 2016). As a first step, one has to reach clarity about which model specifications can be considered as being about equally “reasonable” alternatives (e.g. concerning data transformation, error structure, outlier removal, inclusion of covariates and interactions, outcome measures). This clarity can be reached by constructing a directed acyclic graph (DAG) that illustrates the causal pathways between treatment and outcome, along with potential confounders, mediators, and colliders — each of which may influence the *a priori* study hypothesis if included in statistical models (Arif & MacNeil 2023; Borger & Ramesh 2025; Lundberg *et al*. 2021; Pearl 2009). We particularly recommend the excellent tutorials by Wysocki *et al*. (2022) and Cinelli *et al*. (2024) for deciding which covariates must and which must not be included in a sensible model. Decisions of “principled equivalence” (Del Giudice & Gangestad 2021) should be identified in advance based on the DAG. Once consensus on the decision framework is reached, all planned analyses (e.g., all combinations of 5 binary analysis decisions, yielding 2^5^ = 32 versions of the model) can be performed using tools of multiverse analysis (Masur & Scharkow 2020). A specification-curve plot can then reveal which analysis decisions turn out to be most influential for the study outcome and should therefore receive particular attention regarding whether they represent genuine decisions of “principled equivalence”. Parameter estimates and their CIs can be summarized using the described MAMA procedure (as shown in **Fig. 3**) to obtain CI_ext_ (shown in purple in **Fig. 3**). The resulting CI_ext_ is expected to be slightly wider than the typical (i.e., median) width of the CIs across the input models, with the degree of widening depending on the number of models included in the averaging. When interpreting these intervals, researchers should remain explicit about the inference space to which they apply. They are valid for the specific study system, unless heterogeneity in study system effects (**Fig. 1A**) has also been incorporated into the calculation of extended CIs (according to our equation (2)).

Finally, the most optimistic takeaway from our otherwise perhaps sobering review is that *I*^*2*^ estimates exceeding 0.9, and the associated concern that individual research findings may have severely reduced reliability, with nominal 95% CIs effectively capturing only 45%, may in fact be overly pessimistic. A substantial proportion of heterogeneity may instead stem from the use of inappropriate statistical procedures, either at the level of individual studies or during meta-analytic synthesis. At the individual level, greater theoretical grounding and clearly formulated hypotheses are needed; at the synthesis level, care must be taken to avoid pooling fundamentally heterogeneous studies (i.e. “apples and oranges”). Inappropriate statistical procedures can take many forms: conceptual naivety (e.g. conditioning on colliders, using of predictive models for causal inference; Arif & MacNeil 2022; Borger & Ramesh 2025), insufficient methodological rigor (e.g. failing to account for dependence between data points or overdispersion), or questionable research practices such as *P*-hacking or undisclosed *post-hoc* hypothesizing. Meta-analyses (and many-analyst syntheses) are by no means immune to analogous problems, and are further complicated by selecting studies that are comparable in terms of the hypothesis being tested, by choosing an appropriate effect size measure, and by methodological challenges associated with standardizing effect sizes across studies (Auspurg & Brüderl 2024a, b). Greater awareness of these issues may help reducing the magnitude of *I*^*2*^ and may help increasing our confidence in the underlying data and analysis. Finally, meta-analyses may particularly advance our thinking about biological processes if they succeed at explaining what makes apples differently affected from oranges. When different operationalisations of a broad biological question lead to high heterogeneity, this heterogeneity can be informative: understanding its sources helps refine the underlying biological hypotheses rather than merely signalling a lack of reproducibility.

### Summary of conceptual insights and concluding remarks

- Empirical studies typically have multiple sources of error beyond sampling noise (**Fig. 1**).
- The legitimate inference space is limited by the sources of error that are accounted for (**Fig. 1**), hence, typically it is very narrow, covering just a single way of data analysis.
- The replication crisis logically follows from extrapolation to larger inference spaces than legitimate.
- Confidence intervals and the *α* threshold can be adjusted to account for multiple sources of error and to expand the legitimate inference space (**Fig. 2**).
- Multiverse analysis should become a standard procedure for empirical studies with complex data.
- Many-analyses meta-analysis (MAMA) is a unifying tool for incorporating the multiverse into extended CIs and for looking beyond the usual inference space (**Fig. 3**).
- Meta-analysis is also useful for verifying that CIs derived from randomization procedures correspond to the intended inference space (checking validity beyond a single dataset).
- Published estimates of heterogeneity (*I*^*2*^) should be regarded with caution, because meta-analysts may have been unaware of the many factors that can lead to inflated heterogeneity estimates.
- Truly arbitrary analysis decisions (of principled equivalence) seem to induce much less heterogeneity than most of the recent literature seems to suggest (**Fig. 4**).
- Meta-analysis, and even MAMA, should avoid pooling fundamentally heterogeneous studies or approaches (**Fig. 4**), as doing so fosters the misleading impression of low confidence in individual research findings.

We use statistical models to distinguish meaningful patterns from noise in our data, and we are tempted to generalise our findings from specific populations of individuals or situations to broader research questions (Yarkoni 2022). However, conventional 95% CIs only account for sampling noise 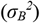, and thus limit us to the picture of the real world that we capture in our specific experimental settings and analyses. As a result, even when we control for all dependencies between datapoints, conventional 95% CIs do not include the real-world variation that we intentionally remove by holding certain experimental design factors constant (Brunswik 1949; Gelman 2015, 2018). Our approach of using extended confidence intervals raises awareness about how limited the legitimate inference space of many published research findings actually is, and it provides a tool for looking beyond that narrow horizon.

## Acknowledgements

We are grateful to Shinichi Nakagawa, Markus Neuhäuser, Julia Rohrer and Holger Schielzeth for their thoughtful and valuable feedback on an earlier version of this manuscript. We further thank colleagues at the Max Planck Institute for Biological Intelligence for constructive feedback and discussions.

## Author contributions

Conceptualization: WF and UK. Simulations and mathematical derivations: UK. Writing: UK and WF.

