## Supplementary Material for "Are our 95% CIs only worth 45% confidence? A heterogeneity-aware method for extending inference"

### Are 95% CIs only worth 45% confidence?

May 7, 2026

### 1. Formal derivations

#### 1.1 Formal derivation of the extension factor $y$

The conventional 95% CIs to our parameter estimate $m$ are given by:

$$95\%\,\text{CI limits} = m \pm (SE \times z_{0.025}) \tag{1}$$

where the quantile $z_{0.025}$ is usually approximated as 1.96 and we calculate the standard error SE as $\sqrt{\sigma_B^2}$ (Borenstein *et al.* 2021; see **Fig. 1** in the main text for the meaning of subscripts).

We are thus looking for a factor $y$ that extends the conventional 95% CIs to 95% $\text{CI}_{\text{ext}}$:

$$95\%\,\text{CI}_{\text{ext}}\,\text{limits} = m \pm (SE \times z_{0.025} \times y) \tag{2}$$

We choose $y$ to include all sources of uncertainty and thus increase the standard error from $\sqrt{\sigma_B^2}$ to $\sqrt{\sigma_B^2 + \sigma_A^2 + \sigma_C^2}$. Thus:

$$y \times \sqrt{\sigma_B^2} = \sqrt{\sigma_B^2 + \sigma_A^2 + \sigma_C^2} \Leftrightarrow y = \sqrt{\frac{{\sigma_A}^2 + {\sigma_B}^2 + {\sigma_C}^2}{{\sigma_B}^2}} \tag{3}$$

$I^2$ is defined as the proportion of the total variation between studies that is due to heterogeneity, which is:

$$I^2 = \frac{{\sigma_A}^2 + {\sigma_C}^2}{{\sigma_B}^2 + {\sigma_A}^2 + {\sigma_C}^2} = 1 - \frac{{\sigma_B}^2}{{\sigma_B}^2 + {\sigma_A}^2 + {\sigma_C}^2} \Leftrightarrow \sigma_B^2 = (1 - I^2) \times (\sigma_B^2 + \sigma_A^2 + \sigma_C^2) \tag{4}$$

By substituting ${\sigma_B}^2$ in equation (3) with the value in equation (4), we obtain:

$$y = \sqrt{\frac{1}{1 - I^2}} \tag{5}$$

and finally, by substituting $y$ into equation (2):

$$95\%\,\text{CI}_{\text{ext}}\,\text{limits} = m \pm \left(SE \times z_{0.025} \times \sqrt{\frac{1}{1 - I^2}}\right) \tag{6}$$

This is equivalent to multiplying the standard errors of individual studies with the square root of the dispersion parameter $\sqrt{H^2}$ (Holzmeister *et al.* 2024), which is another measure of heterogeneity in meta-analyses. Factor $y$ has also been derived by Borm *et al.* (2009) using a slightly different line of reasoning.

---

#### 1.2 Formal derivation of $I^2_{MAMA}$

The software-inbuilt meta-analytically derived heterogeneity $I^2$ of a many-analyses study assumes that all three sources of variation are present. Since the teams of analysts receive the same dataset, there is no variance in true study system effects, which means $\sigma_A^2 = 0$. Thus, the equation for deriving $I^2$ simplifies to:

$$I^2 = \frac{{\sigma_A}^2 + {\sigma_C}^2}{{\sigma_B}^2 + {\sigma_A}^2 + {\sigma_C}^2} = \frac{{\sigma_C}^2}{{\sigma_B}^2 + {\sigma_C}^2} \tag{7}$$

However, for a many-analyses meta-analysis (MAMA), this equation is not correct and underestimates $I^2$, because it assumes that there is true sampling noise ${\sigma_B}^2$, resulting from the selection of a different subset of individuals from the total population for each of the individual analyses. This is not the case, all the variation that we see between analyses is attributable to differences in the choice of analysis made by different researchers. Consequently, the total variance is given by ${\sigma_T}^2 = {\sigma_C}^2$, and the conventional estimator, which implicitly substracts an assumed sampling variance ${\sigma_B}^2$ from the total variance, can be written as:

$$I^2* = \frac{{\sigma_T}^2 - {\sigma_B}^2}{{\sigma_T}^2} = \frac{{\sigma_C}^2 - {\sigma_B}^2}{{\sigma_C}^2} \Leftrightarrow {\sigma_B}^2 = {\sigma_C}^2 \times (1 - I^2*) \tag{8}$$

By replacing ${\sigma_B}^2$ in equation (7) with the value in equation (8), we obtain the correct MAMA-derived heterogeneity estimate $I^2_{MAMA}$:

$$I^2_{MAMA} = \frac{{\sigma_C}^2}{{\sigma_C}^2 \times (1 - I^2*) + {\sigma_C}^2} = \frac{{\sigma_C}^2}{{\sigma_C}^2 \times (1 - I^2 * +1)} = \frac{1}{2 - I^2*} \tag{9}$$

Thus, depending on the (incorrect) size of $I^2*$ in the MAMA, $I^2_{MAMA}$ can be substantially larger (**Fig. S1**). At the lower bound of $I^2* = 0$, the MAMA assumes that all variation between studies ${\sigma_T}^2$ is due to sampling noise ${\sigma_B}^2$ (i.e. ${\sigma_T}^2 = {\sigma_B}^2$) and not to the choice of analyses ${\sigma_C}^2$, which is seriously flawed because in reality it is the opposite. No matter how small the variation between studies, it should all be due to the choice of analysis (${\sigma_T}^2 = {\sigma_C}^2$). Now, if each researcher in a many-analyses study is given a new dataset sampled from the total population and performs exactly the same analysis as before, we can derive $I^2$ as in equation (1) and obtain ${\sigma_C}^2/({\sigma_B}^2 + {\sigma_C}^2) = {\sigma_T}^2/({\sigma_T}^2 + {\sigma_T}^2) = 0.5$, which is our corrected many-analyses heterogeneity $I^2_{MAMA}$. On the other hand, when there is a substantial amount of heterogeneity between the many-analyses estimates ($I^2* > 0.9$), the $I^2_{MAMA}$ is less than 1% larger.

Cochran's $Q$ is a test statistic used to assess the homogeneity of parameter estimates from individual studies. It is calculated as the sum of the squared deviations of each study's estimate $m_i$ from the overall meta-analytic estimate $\bar{m}$, weighting the contribution of each study in the same way as in the meta-analysis (Cochran 1954).

$$Q = \sum_{i=1}^{k} \frac{(m_i - \bar{m})^2}{w_i}$$

where $w_i$ is the weighting factor, which is usually the squared standard error of the study estimate $m_i$, i.e. its variance and where $\bar{m} = \sum \frac{m_i}{w_i} / \sum \frac{1}{w_i}$. The $Q$-statistic follows a $\chi^2$ distribution with $k - 1$ degrees of freedom, where $k$ is the number of studies included in the meta-analysis. Testing for the presence of statistical heterogeneity with Cochran's $Q$ has low power when the sample size is small, and excessive power when there are many and large studies (Hardy & Thompson 1998). The important aspect for us here is that $I^2$ can be derived from Cochran's $Q$ (Higgins & Thompson

2002; Higgins *et al.* 2003) as follows:

$$I^2 = \frac{Q - k + 1}{Q}$$

We can easily see two things:

1. $I^2$ is not defined when $Q = 0$, which would be the case if all study estimates $m_i$ were equal and there was no variation between studies.
2. $I^2$ can take negative values if $Q$ was small and $k$ was large.

Normally, negative $I^2$ values are set to zero (Higgins *et al.* 2003), but to obtain the correct MAMA-derived heterogeneity estimate $I^2_{MAMA}$, we also use the negative $I^2$ estimates so that $I^2_{MAMA}$ varies between 0 and 1.

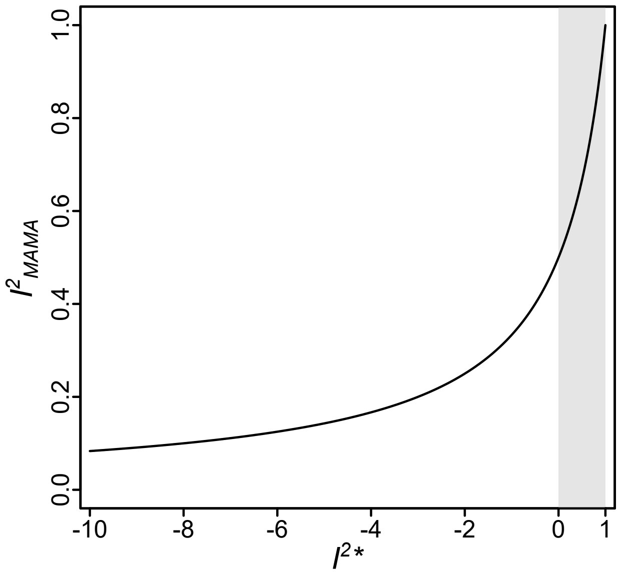

***Fig. S1*** | *Relationship between the meta-analytically derived heterogeneity $I^2*$ and the corrected many-analyses heterogeneity $I^2_{MAMA}$. The grey shaded area represents the normal range of $I^2$, i.e. negative $I^2$ values are set to zero. When negative $I^2*$ values are used to represent underdispersion, the $I^2_{MAMA}$ values vary between 0 and 1.*

---

#### 1.3 Formal derivation of $\mathrm{SE}_{\mathrm{MAMA}}$

The following derivations use a plug-in approach, where quantities estimated from the data are subsequently treated as fixed. Accordingly, the formulas should be interpreted as large-sample approximations.

In a conventional random-effects meta-analysis, the standard error of the meta-analytic mean effect ($SE_{META}$) is calculated as follows (Borenstein *et al.* 2021):

$$SE_{META} = \sqrt{\frac{1}{\sum 1/({\sigma_A}^2 + {\sigma_B}^2 + {\sigma_C}^2)}}$$

If we assume that all individual studies have the same sample size, then after applying Fisher's $z$-transformation (which is variance-stabilizing; Fisher 1921), the within-study variance ${\sigma_B}^2$ (and, consequently, the standard errors) of all individual studies are approximately the same. Since ${\sigma_A}^2$ and ${\sigma_C}^2$ are treated as common between-study variance components derived from $\tau$, this equation simplifies to:

$$SE_{META} = \sqrt{\frac{{\sigma_A}^2 + {\sigma_B}^2 + {\sigma_C}^2}{N}}$$

where $N$ is the number of individual studies included in the meta-analysis.

In a many-analyses meta-analysis (MAMA), it would be incorrect to calculate the standard error in the same way, because a regular meta-analysis assumes that all studies are based on independent data. However, in a MAMA, all research teams analyze the same dataset, and any variation between analyses arises solely from the choice of analysis made by different researchers (i.e. ${\sigma_T}^2 = {\sigma_C}^2$). As a result, the variance due to the choice of analysis ${\sigma_C}^2$ is reduced (and ${\sigma_A}^2 = 0$), while the sampling noise ${\sigma_B}^2$ is treated as a representative within-study sampling variance.

The correct standard error for the MAMA-derived average effect size ($SE_{MAMA}$) is therefore given by the following formula:

$$SE_{MAMA} = \sqrt{{\sigma_B}^2 + ({\sigma_C}^2/N)} \tag{10}$$

where $N$ is the number of teams of analysts in the MAMA. Thus, to calculate $SE_{MAMA}$, we need estimates of ${\sigma_C}^2$ and ${\sigma_B}^2$. In a MAMA, the variance in effect sizes reported by the teams of analysts (${\sigma_T}^2$) is an estimate of ${\sigma_C}^2$. Using the meta-analytically derived heterogeneity $I^2*$ and equation (8), we can derive ${\sigma_B}^2$ as:

$${\sigma_B}^2 = {\sigma_C}^2 \times (1 - I^2*) \Leftrightarrow {\sigma_B}^2 = {\sigma_T}^2 \times (1 - I^2*)$$

Finally, by substituting our estimates of ${\sigma_C}^2$, ${\sigma_B}^2$ and the number of analysts $N$ into equation (10), we calculate the standard error $SE_{MAMA}$ of a MAMA-derived average effect.

---

### 2. Many-analyses meta-analysis (MAMA)

#### 2.1 Methods

We simulated 10,000 datasets, each containing $N = 100$ observations originating from 5 years, with 20 individuals per year. The variable *year* fitted as a random intercept explained approximately 10% of the variance in the normally distributed response variable $y$. In addition, we simulated four predictors: *weight* (continuous, gamma-distributed with shape parameter 3), *wing* (continuous, normally distributed with a mean of 55 and standard deviation of 5), *sex* (binary factor with levels *female* and *male*, with probabilities 0.15 and 0.85, respectively), and *treatment* (categorical with three levels: treatment, control 1, control 2). The *treatment* variable had no effect on $y$, while *weight* and *sex* had effects detectable with approximately 20% power, and *wing* had an effect detectable with 30% power. We then introduced 20% missing values in *wing*, which substantially reduced its power.

We simulated a collapsible model—one in which the inclusion of covariates should not influence the treatment effect. In real data, however, this may not hold, as non-collapsibility can arise from collinearity, confounding, and interactions, and is an inherent property of the logit and probit link functions used in generalized linear models with a binomial error structure (Greenland *et al.* 1999).

To mimic a many-analyst situation, we defined seven analytical decisions (D1–D7) that analysts might take:

- **D1**: Use the full dataset or remove one of the two control groups.
- **D2**: Include or exclude outliers (defined as the minimum and maximum values of $y$).
- **D3**: Use $y$ or $\sqrt{y}$ as the response variable.
- **D4**: Include or exclude *weight* as a covariate.
- **D5**: Include or exclude *sex* as a covariate.
- **D6**: Remove rows with missing *wing* values or impute missing values with the mean.
- **D7**: Model *year* as either a fixed or random effect.

These decisions yield $2^7 = 128$ possible combinations. For each of these 128 combinations, we fitted linear (mixed-effects) models and derived standardized effect sizes ($Zr$; see Gould *et al.* (2025) and below for details). We then analyzed these effect sizes in a meta-analysis and obtained estimates of ${\sigma_C}^2$, ${\sigma_B}^2$, and $SE_{MAMA}$ for all 10,000 simulation runs.

---

#### 2.2 Results

By summarizing these 10,000 estimates, we quantified the amount of variation in ${\sigma_C}^2$ and ${\sigma_B}^2$, the type I error rate when cherry-picking among the 128 analytical combinations, and estimated the precision of the 95% confidence intervals derived from $SE_{MAMA}$.

We found that the sampling noise ${\sigma_B}^2$ was relatively constant across the 10,000 estimates (mean ${\sigma_B}^2 = 0.016$, standard deviation ${\sigma_B}^2 = 0.0014$, coefficient of variation ${\sigma_B}^2 = 0.086$; **Fig. S2A**), whereas the effects of analytical choices were highly variable (average ${\sigma_C}^2 = 0.0033$, standard deviation ${\sigma_C}^2 = 0.0024$, coefficient of variation ${\sigma_C}^2 = 0.738$; **Fig. S2B**).

When selecting the most significant result among the 128 combinations, the type I error rate increased to 20.7%. This is four times higher than the nominal 5% threshold, yet much lower than the rate of $1 - 0.95^{128}$ that would be expected if all models were independent.

The 95% confidence intervals derived from $SE_{MAMA}$ captured 97.4% of the meta-analytically derived summary values. This coverage rate is slightly higher than the nominal 95% and may be explained by differences in sample size across the 128 models, which result from analytical decisions D1 (removal of one control group) and D6 (handling of missing values in *wing*). In a separate simulation where sample sizes were constant across all models, the 95% confidence intervals derived from $SE_{\mathrm{MAMA}}$ captured 94.8% of the summary values, closely matching the nominal level.

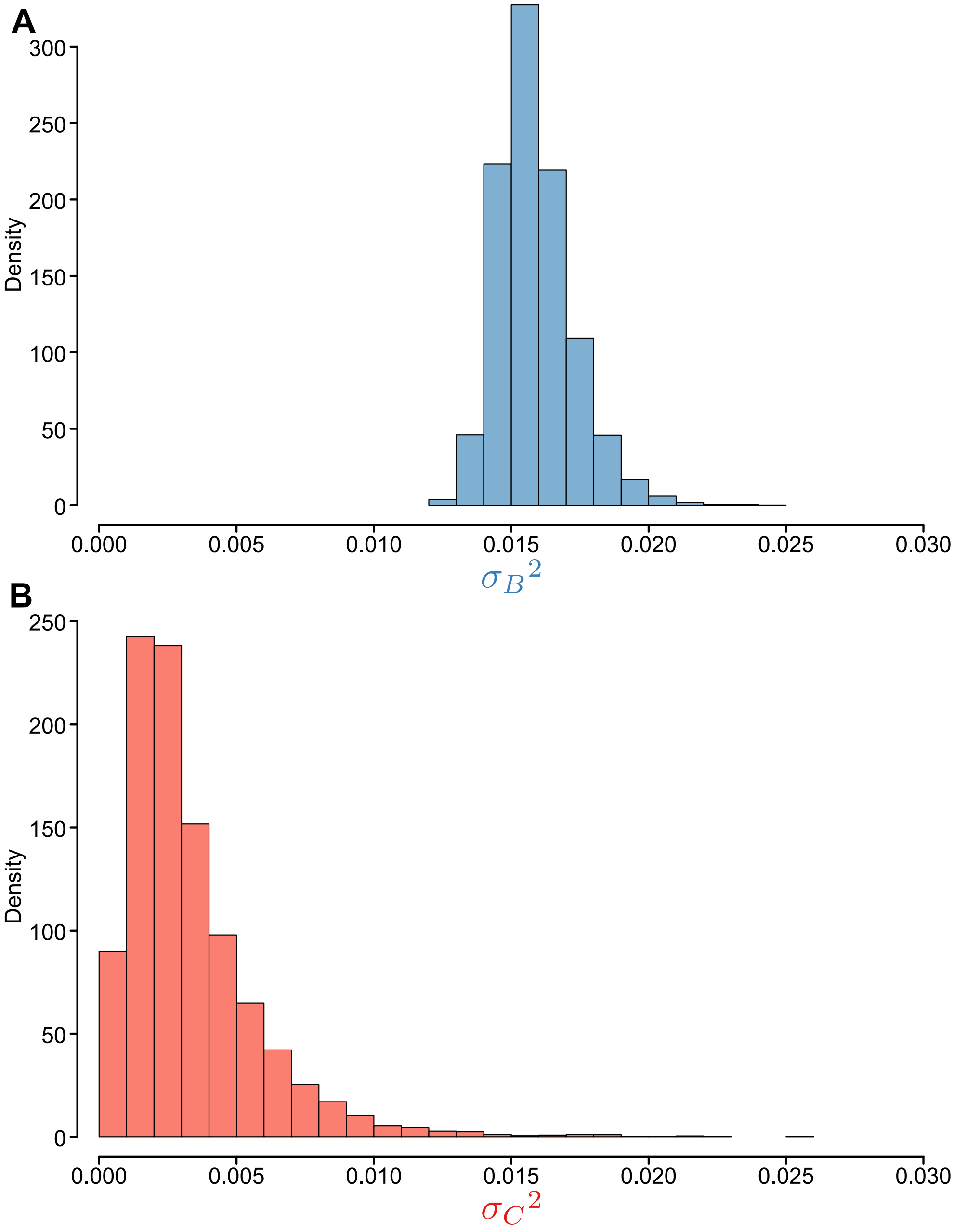

***Fig. S2** | Variation in (A) sampling noise $\sigma_B{}^2$ and (B) analytical choices $\sigma_C{}^2$ across 10,000 simulation runs. Note the much larger variation in $\sigma_C{}^2$ compared to $\sigma_B{}^2$.*

---

### 3. Reanalysis of the 'blue tit dataset'

Gould *et al.* (2025) provided data from a brood-size manipulation experiment in wild blue tits (*Cyanistes caeruleus*) breeding in nest boxes ('blue tit dataset') to multiple independent research teams. Each team was asked to answer the question: 'To what extent is the growth of nestling blue tits influenced by competition with siblings?' This question is deliberately loosely defined and allows for multiple analytical pathways.

In the present analysis, we narrow the focus and define *Net Manipulation* as our treatment variable, representing the experimentally controlled change in brood size by adding or removing 0–4 chicks. We believe that in an experimental context, the primary goal is to estimate the effect of the experimental treatment, rather than natural variation. We consider *Net Manipulation* therefore the most direct operationalization of what is meant by 'competition with siblings' in the original question. We use *Day 14 Chick Mass* as our outcome variable, as it reflects 'growth' more directly than alternative measures such as *Day 14 Tarsus Length* or *N Fledglings*.

We reuse this dataset here for illustrative purposes, not to critique previous analyses, but because one of the authors of the present study participated in the original project as a research team and is therefore familiar with the data structure and experimental design.

#### 3.1 Methods

##### Setting up a directed acyclic graph (DAG) for causal inference

Using the 'blue tit dataset', we developed models that corresponded to the three types of analytical decisions defined by Del Giudice & Gangestad (2021): (1) Type E decisions ('principled equivalence'), (2) Type U decisions ('uncertain equivalence'), and (3) Type N decisions ('non-equivalence'). Our classification of these decision types was guided by a directed acyclic graph (DAG) (**Fig. S3**), which summarizes the assumed causal relationship between *Net Manipulation* (treatment) and *Day 14 Chick Mass* (outcome).

We incorporated intermediate variables (*Post-Treatment Brood Size*), exogenous confounders (*Chick Sex*, environmental factors: *Year*, *Date*, *Area*, and genetic factors: *Nest of Origin*), and baseline covariates (*Pre-Treatment Brood Size*). In addition, we included causal paths to alternative outcome variables (*Day 14 Tarsus Length*, *N Fledglings*), with further mediators on the path to *N Fledglings* (*Day 14 Brood Size*). To reflect how researchers might conceptualize the causal structure, we also included purely deterministic variables that were derived from other nodes (*Early Mortality* = *Day 14 Brood Size* - *Post-Treatment Brood Size* and *Late Mortality* = *N Fledglings* - *Day 14 Brood Size*). These variables do not represent independent causal mechanisms but may nonetheless be included as covariates in modeling, particularly *Early Mortality*, when estimating the effect of *Net Manipulation* on *Day 14 Chick Mass*. Finally, *Net Manipulation* was decomposed into *N Chicks Added* and *N Chicks Removed* in the 'blue tit dataset', which might encourage researchers to fit these in their causal model.

##### Causal pathways in the DAG

In the DAG, the causal path from *Net Manipulation* to *Day 14 Chick Mass* was assumed to be mediated by the latent (unmeasured) variable *Feeding Rate*. A priori, the sign of the causal relationship between *Post-Treatment Brood Size* and *Day 14 Chick Mass* was ambiguous. In experimentally enlarged broods, parents might fully compensate by increasing total provisioning effort (resulting in no net effect per chick), fail to meet the increased demand (leading to a negative effect on individual chick mass), or be stimulated to provision more intensively (resulting in a positive effect). In experimentally reduced broods, parents might reduce their total provisioning effort proportionally (no effect), continue feeding at the same overall rate (positive effect), or become understimulated and reduce provisioning disproportionately (negative effect). A similar line of reasoning applies to the alternative outcome variables *Day 14 Tarsus Length* and *N Fledglings*.

#### Identifying confounders, mediators and colliders

In causal inference, we distinguish between confounders, mediators and colliders. These roles are not mutually exclusive, meaning that a single variable can function as a mediator in one causal pathway and as a collider in another. Confounders should generally be conditioned upon to remove confounding bias and reduce residual variation in the outcome. In contrast, mediators and colliders should typically not be conditioned upon if the research objective is to estimate the total effect of the experimental treatment on the outcome. Including mediators as covariates can introduce overcontrol bias, while including colliders (or their descendants) can lead to endogenous selection bias (Elwert & Winship 2014).

Along the causal path from *Net Manipulation* to *Day 14 Chick Mass*, confounders are the exogenous baseline variables related to the environment (*Year*, *Date*, *Area*) and to the genetic make-up of the chicks (*Nest of Origin*, *Chick Sex*). The sign of these effects (positive or negative) is either unknown or not of primary interest, because these variables are typically modeled as categorical or random effects. *Pre-Treatment Brood Size* is a confounder as well, but its effect on *Post-Treatment Brood Size* has an expected positive sign. Conditioning on it adjusts for natural brood size variation prior to the treatment.

Mediators are intermediate variables that transmit the effect of *Net Manipulation* on the outcome. In the causal pathways to *Day 14 Chick Mass* and *Day 14 Tarsus Length*, only *Post-Treatment Brood Size* is a mediator. In the pathway to *N Fledglings*, additional mediators are *Day 14 Brood Size* and the deterministically defined variables *Early Mortality* and *Late Mortality*.

Colliders are defined as variables with two or more incoming arrows from distinct sources. In our DAG, no variables meet this criterion; that is, there are no colliders.

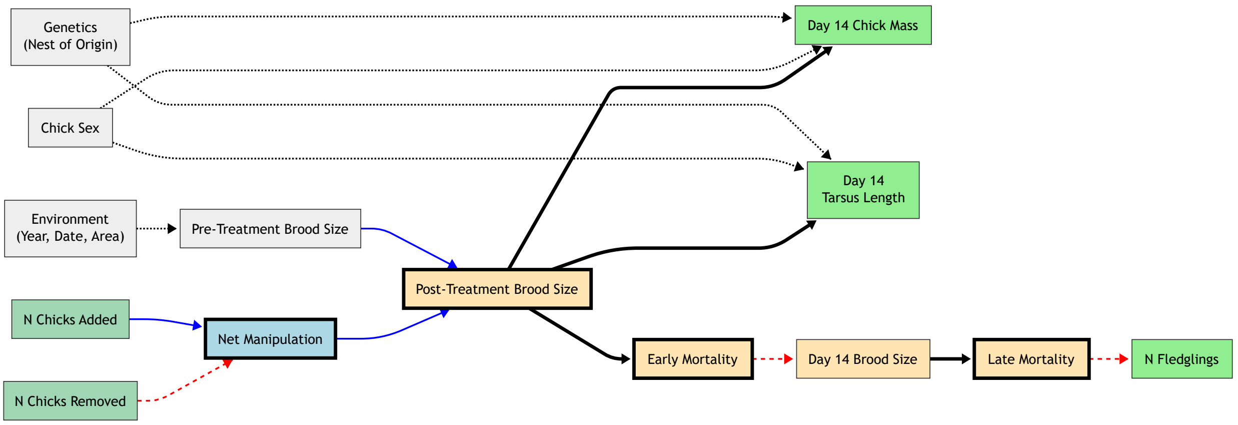

***Fig. S3** | A directed acyclic graph (DAG) illustrating the presumed causal structure in the 'blue tit dataset' of Gould et al. (2025). Node colors: blue = treatment ("Net Manipulation"); turquoise = treatment components ("Number of Chicks Added" and "Number of Chicks Removed"); green = primary outcome ("Day 14 Chick Mass"); gray = exogenous baseline covariates; orange = mediators (e.g., "Post-Treatment Brood Size"). Borders: thick = deterministic (computed) variables. Edge styles: solid blue = positive effect; dashed red = negative effect; solid black = effect of interest with a prior unknown sign; dotted black = uncertain or random effect. Note that "Day 14 Tarsus Length" and "N Fledglings" are also depicted in green because they are alternative outcome measures. We used the Mermaid software for creating the illustration (Sveidqvist 2025).*

A posteriori, we observed a non-linear relationship between *Day 14 Chick Mass* and *Net Manipulation*: while brood size reductions had no effect on *Day 14 Chick Mass*, brood size enlargements had a pronounced negative effect. Consequently, fitting *Net Manipulation* as a continuous covariate that includes both reductions and enlargements violates the linearity assumption of linear regression (Gelman & Hill 2007). To address this, we also split the dataset into two comparisons: reductions versus unmanipulated broods and enlargements versus unmanipulated broods.

##### Type E Decisions (Principled Equivalence)

1. **Handling exogenous confounders in different ways:** *Chick Sex* may affect *Day 14 Chick Mass* but is not causally linked to the treatment. We handled *Chick Sex* in three ways: (1) including it as recorded in the original data ($N = 1082$ individuals with missing sex information are hence dropped from the analysis, representing 29.8% of the full dataset; *model 1*), (2) replacing missing data with the mean sex value (coding females as 0 and males as 1 and chicks with missing sex information as the overall mean; *model 2*), or (3) excluding *Chick Sex* from the model (*model 3*).

   Similarly, *Rear Area* is a factor with 9 levels, uncorrelated with the treatment and not on the causal path between treatment and outcome. Including it as a random effect explained 1.1% of the variance in *Day 14 Chick Mass* in the full dataset. Thus, the decision to include or exclude *Rear Area* was also considered a case of principled equivalence (*model 4*).

2. **Transformation of the dependent variable:** Transforming the dependent variable (e.g., $\log_e(y)$ or $\sqrt{y}$) does not systematically bias $t$-values but may aid in model fitting. However, it alters parameter estimates, complicating direct comparisons with untransformed data. Because effect size estimates were based on the $t$-value (see below and Gould *et al.* 2025), transformation of the dependent variable is considered a case of principled equivalence (*model 5*).

3. **Outlier removal:** Removing outliers that result from processes unrelated to the treatment should not introduce systematic bias. Additionally, if the objective is to assess treatment effects on the 'average' chick, excluding severely underfed chicks may be justifiable. Thus, we also regard outlier removal as a case of principled equivalence (*model 6*).

##### Type U Decisions (Uncertain Equivalence)

1. **Fitting baseline covariates or predictors of a mediator:** Conditioning on *Pre-Treatment Brood Size* removes the natural variation in brood size that existed prior to the treatment (*model 7*). By doing so, we do not introduce bias. In fact, this adjustment can improve the precision of the estimated effect of *Net Manipulation* on *Day 14 Chick Mass* by reducing residual variation. However, it shifts the interpretation of the model: instead of estimating the average treatment effect of changing brood size on chick mass across the full range of brood sizes (marginal treatment effect), the model now estimates the effect of experimentally changing brood size conditional on initial brood size (conditional treatment effect). That is, it addresses the question: What is the average effect of experimentally altering brood size, independent of natural variation in initial brood size?

   This distinction may lead to different estimates if parents are capable of adjusting provisioning effort only up to a certain brood size (e.g. 10 chicks), but cannot compensate beyond that threshold. In such cases, the treatment effect depends on *Pre-Treatment Brood Size* (treatment effect heterogeneity), which leads to a non-linear relationship between brood size and outcome. Notably, this logic of non-homogeneous treatment effects is consistent with the patterns we observed empirically when comparing experimental brood size enlargements versus reductions.

2. **Use of alternative outcome measures of treatment that align with the research question:** In addition to *Day 14 Chick Mass*, researchers also measured *Day 14 Chick Tarsus Length*. Although we based effect size estimates on the $t$-value, it is not clear whether mass and tarsus length are equivalent outcome measures for assessing the treatment effect. For example, one might expect a stronger effect on mass, as a measure of condition, than on tarsus length (*model 8*).

##### Type N Decisions (Principled Nonequivalence)

1. **Use of alternative outcome measures of treatment that do not align closely with the research question:** As a discrete, downstream outcome, fledging success may reflect cumulative effects of multiple processes (including survival and condition), but is less directly interpretable as a response to treatment and not closely aligned with the original research question posed by Gould *et al.* (2025). It is rather a qualitative measure of offspring growth. We also acknowledge that Gould *et al.* (2025) excluded analyses that did not use outcomes interpretable as growth or size. We include it here for illustrative purposes. *N Fledglings* is partly a deterministic function of *Post-Treatment Brood Size*, and hence not independent of the treatment even under the null hypothesis. Thus, we model survival until fledging in a generalized linear mixed-effects model with binomial error structure, using *N Fledglings* and *Post-Treatment Brood Size - N Fledglings* connected with the `cbind()` function as our response variable (*model* 9).

2. **Conditioning on mediators:** Including mediators as covariates in the model can introduce overcontrol bias (Elwert & Winship 2014), as this distorts the causal path from *Net Manipulation* to *Day 14 Chick Mass*. We allowed for this bias by including *Post-Treatment Brood Size* as a covariate (*model* 10).

3. **Conditioning on descendants of a mediator (occurring before day 14):** *Early Mortality* might appear to be a relevant control variable (model 11), especially if one suspects that mortality, whether elevated in enlarged or reduced in smaller broods, moderates the treatment effect. Similarly, *Day 14 Brood Size* might be included based on a similar rationale (model 12). However, both are downstream consequences of the treatment, and conditioning on them introduces post-treatment bias.

4. **Conditioning on descendants of a mediator (occurring after day 14):** *N Fledglings* and *Late Mortality* are also downstream consequences of the treatment, but they occur after the outcome (*Day 14 Chick Mass*) has been measured. Including these variables as covariates introduces post-treatment bias and violates temporal ordering, whereby variables observed after the outcome should not be included as predictors in a causal model (*model* 13 and *model* 14) (Pearl 2009).

5. **Conditioning on alternative outcome measures:** *Day 14 Chick Tarsus Length* is highly correlated with *Day 14 Chick Mass*. By including *Day 14 Chick Tarsus Length* as a covariate, we effectively residualize mass over size, shifting the research question from the treatment's effect on growth to its effect on condition, leading to principled nonequivalence (*model* 15).

   Similarly, if we use *Day 14 Chick Tarsus Length* as the outcome variable and condition on *Day 14 Chick Mass*, the analysis is reversed, but the conceptual issue remains the same: the treatment effect is redefined (*model* 16). While this model is unlikely to be chosen by a researcher in practice, we include it for illustrative purposes.

6. **Conditioning on components of the treatment:** By including *N Chicks Added* (*model* 17) or *N Chicks Removed* (*model* 18) alongside *Net Manipulation* as covariates introduces redundancy. These variables are components of the treatment itself, and controlling for them effectively fits the treatment twice, leading to overcontrol bias and conceptual inconsistency.

##### Alternative operationalizations of outcome and treatment

In addition to the primary specification, we considered alternative definitions of both the outcome variable and the treatment, reflecting analytical decisions that may fall into the category of uncertain (Type U) or, in some cases, non-equivalent (Type N) choices.

First, we explored an alternative formulation in which the roles of *Day 14 Chick Mass* and *Day 14 Chick Tarsus Length* were systematically exchanged across the full set of models. That is, wherever one variable appeared as an outcome or predictor in the original specification, it was replaced by the other in the alternative specification.

This generates a corresponding set of models with identical structure but permuted roles of the two variables. Under this transformation, all models are altered, but some coincide with different models in the original specification. In particular, *model 2* in the alternative specification is identical to *model 8* in the original specification, and *model 15* and *model 16* exchange roles under this transformation.

This reparameterization does not introduce new types of analytical decisions, but allows us to assess whether conclusions depend on how growth is operationalized.

Second, the experimental treatment (*Net Manipulation*) can be operationalized either as a continuous variable spanning brood-size reductions, unmanipulated broods, and brood-size enlargements, or through alternative specifications that relax the assumption of a single linear effect across this range.

In the primary continuous model, *Net Manipulation* was fitted across its full range (-4 to +4 chicks), implying a single linear relationship between brood-size manipulation and the outcome. However, given the empirically observed asymmetry between brood-size reductions and enlargements, we additionally fitted separate models for the two subsets of the data (-4 to 0 and 0 to +4), thereby estimating treatment effects for reductions and enlargements independently. In these models, unmanipulated broods (value = 0) serve as a reference point within each subset, but not as strict experimental controls.

As a further specification, we also fitted the treatment as a two-level factor contrasting enlarged and reduced broods directly. This avoids using unmanipulated broods as a reference category, but changes the estimand from the effect of manipulation relative to unmanipulated broods to the difference between opposite manipulation directions. Because these formulations impose different assumptions about the functional form of the treatment effect, we treat this choice as Type U.

##### Model fitting and parameter extraction

We fitted all models in `lme4` (Bates *et al.* 2015), and either extracted the $t$-value and the degrees of freedom or the parameter estimate with its standard error for *Net manipulation.* The degrees of freedom were estimated using the `lmerTest` package (Kuznetsova *et al.* 2017) in `R`.

We transformed the $t$-value ($t$) and the degrees of freedom ($df$) to a correlation coefficient as described in Gould *et al.* (2025):

$$r = \frac{t}{\sqrt{t^2 + df}}$$

Next, we applied Fisher's $Z$-transformation:

$$Z_r = atanh(r)$$

The standard error of $Z_r$ was calculated as:

$$SE = \sqrt{\frac{1}{df - 2}}$$

Specifically, we fitted the following models:

```
rm(list=ls())
if("lme4" %in% rownames(installed.packages()) == FALSE)
  { install.packages("lme4") }
if("lmerTest" %in% rownames(installed.packages()) == FALSE)
  { install.packages("lmerTest") }
if("RColorBrewer" %in% rownames(installed.packages()) == FALSE)
  { install.packages("RColorBrewer") }

require(lme4)
require(lmerTest)
require(RColorBrewer)
require(ManyEcoEvo)

out <- data.frame(matrix(rep(NA,9*3*18), ncol=9))
colnames(out) <- c("analysis","model","type","b","SE","t","df","Zr","SEZr")

# Specify path ---
path <- "H:\\Knief\\Documents\\Stats\\MetaHeterogeneity\\"

# ----------------------------------------------------------------------------
# Data preparation -----------------------------------------------------------
Subsets <- c("CRE", "CE", "CR")
k <- 1
for(i in 1:3) {
  SubsetData <- Subsets[i]

  # Read data ---
  data(blue_tit_data)
  dat <- as.data.frame(blue_tit_data)

  # Replace "." with NA ---
  dat[] <- lapply(dat, function(x) {
    if (is.factor(x)) x <- as.character(x)
    x[x=="."] <- NA
    return(x)
  })

  # Rename columns ---
  colnames(dat)[colnames(dat)=="day_14_weight"] <- "Day14Mass"
  colnames(dat)[colnames(dat)=="day_14_tarsus_length"] <- "Day14Tarsus"
  colnames(dat)[colnames(dat)=="chick_sex_molec"] <- "ChickSex"
  colnames(dat)[colnames(dat)=="net_rearing_manipulation"] <- "NetManipulation"
  colnames(dat)[colnames(dat)=="rear_d0_rear_nest_brood_size"] <-
      "PreTreatBrSize"
  colnames(dat)[colnames(dat)=="rear_Cs_in"] <- "NChicksAdded"
  colnames(dat)[colnames(dat)=="rear_Cs_out"] <- "NChicksRemoved"
  colnames(dat)[colnames(dat)=="rear_Cs_at_start_of_rearing"] <-
      "PostTreatBrSize"
  colnames(dat)[colnames(dat)=="d14_rear_nest_brood_size"] <- "Day14BrSize"
  colnames(dat)[colnames(dat)=="number_chicks_fledged_from_rear_nest"] <-
     "NFledglings"
```

```
colnames(dat)[colnames(dat)=="hatch_year"] <- "Year"
colnames(dat)[colnames(dat)=="Date_of_day14"] <- "JulianDay14"
colnames(dat)[colnames(dat)=="rear_area"] <- "AreaID"
colnames(dat)[colnames(dat)=="hatch_nest_breed_ID"] <- "OriginNestID"
colnames(dat)[colnames(dat)=="rear_nest_breed_ID"] <- "RearNestID"
colnames(dat)[colnames(dat)=="rear_nest_trt"] <- "TreatmentCategory"
colnames(dat)[colnames(dat)=="home_or_away"] <- "ChickSwappedYN"

# Prepare data ---
dat$ChickSex <- as.numeric(dat$ChickSex)
dat$NetManipulation <- as.numeric(dat$NetManipulation)
dat$PreTreatBrSize <- as.numeric(dat$PreTreatBrSize)
dat$NChicksAdded <- as.numeric(dat$NChicksAdded)
dat$NChicksRemoved <- as.numeric(dat$NChicksRemoved)
dat$PostTreatBrSize <- as.numeric(dat$PostTreatBrSize)
dat$NFledglings <- as.numeric(dat$NFledglings)
dat$Year <- factor(dat$Year)
dat$OriginNestID <- factor(dat$OriginNestID)
dat$RearNestID <- factor(dat$RearNestID)
dat$TreatmentCategory <- factor(dat$TreatmentCategory)
dat$EarlyMortality <- dat$PostTreatBrSize - dat$Day14BrSize
dat$LateMortality <- dat$Day14BrSize - dat$NFledglings
dat$ChickSex <- ifelse(is.na(dat$ChickSex),
                       mean(dat$ChickSex,na.rm=TRUE), dat$ChickSex)

# Correct EarlyMortality=-1
subset(dat, EarlyMortality==-1)
dat$PostTreatBrSize[which(dat$EarlyMortality==-1)] <- 5
dat$EarlyMortality <- dat$PostTreatBrSize - dat$Day14BrSize

# Create subsets ---
# CE = unmanipulated and enlargements
# CR = unmanipulated and reductions
# CRE = unmanipulated, reductions, enlargements
dat.CRE <- dat
dat.CRE.SexSub <- subset(dat.CRE, ChickSex==1 | ChickSex==2)
dat.CRE.OR <- subset(dat.CRE, Day14Mass > 5.9)
dat.CE <- subset(dat.CRE, TreatmentCategory==5 | TreatmentCategory==7)
dat.CR <- subset(dat.CRE, TreatmentCategory==6 | TreatmentCategory==7)
dat.CE.SexSub <-subset(dat.CRE.SexSub,TreatmentCategory==5 |
                                      TreatmentCategory==7)
dat.CR.SexSub <-subset(dat.CRE.SexSub,TreatmentCategory==6 |
                                      TreatmentCategory==7)
dat.CE.OR <- subset(dat.CRE.OR, TreatmentCategory==5 | TreatmentCategory==7)
dat.CR.OR <- subset(dat.CRE.OR, TreatmentCategory==6 | TreatmentCategory==7)
dat.Survival <- dat[!duplicated(dat$RearNestID), ]
dat.Survival$SurvivalProp <- dat.Survival$NFledglings /
                             dat.Survival$PostTreatBrSize
dat.Survival.CRE <- dat.Survival
dat.Survival.CE <- subset(dat.Survival.CRE, TreatmentCategory==5 |
                                            TreatmentCategory==7)
```

```
dat.Survival.CR <- subset(dat.Survival.CRE, TreatmentCategory==6 |
                                            TreatmentCategory==7)
# ----------------------------------------------------------------------------
# Full data set --------------------------------------------------------------
if(SubsetData=="CRE") {
   dat <- dat.CRE
   dat.SexSub <- dat.CRE.SexSub
   dat.OR <- dat.CRE.OR
   dat.Survival <- dat.Survival.CRE
}
# ----------------------------------------------------------------------------
# Only unmanipulated and enlargements ----------------------------------------
if(SubsetData=="CE") {
   dat <- dat.CE
   dat.SexSub <- dat.CE.SexSub
   dat.OR <- dat.CE.OR
   dat.Survival <- dat.Survival.CE
}
# ----------------------------------------------------------------------------
# Only unmanipulated and reductions ------------------------------------------
if(SubsetData=="CR") {
   dat <- dat.CR
   dat.SexSub <- dat.CR.SexSub
   dat.OR <- dat.CR.OR
   dat.Survival <- dat.Survival.CR
}

# ----------------------------------------------------------------------------
# Models according to Type E Decisions (Principled Equivalence) --------------
m01 <- lmer(Day14Mass ~ NetManipulation + ChickSwappedYN + JulianDay14 +
                        Year + ChickSex + (1|OriginNestID) + (1|RearNestID),
                        data=dat.SexSub, na.action=na.exclude, REML=FALSE,
                        control = lmerControl(optimizer = "bobyqa"))
m02 <- lmer(Day14Mass ~ NetManipulation + ChickSwappedYN + JulianDay14 +
                        Year + ChickSex + (1|OriginNestID) + (1|RearNestID),
                        data=dat, na.action=na.exclude, REML=FALSE,
                        control = lmerControl(optimizer = "bobyqa"))
m03 <- lmer(Day14Mass ~ NetManipulation + ChickSwappedYN + JulianDay14 +
                        Year + (1|OriginNestID) + (1|RearNestID),
                        data=dat, na.action=na.exclude, REML=FALSE,
                        control = lmerControl(optimizer = "bobyqa"))
m04 <- lmer(Day14Mass ~ NetManipulation + ChickSwappedYN + JulianDay14 +
                        Year + ChickSex + (1|OriginNestID) + (1|RearNestID) +
                        (1|AreaID), data=dat,na.action=na.exclude,REML=FALSE,
                        control = lmerControl(optimizer = "bobyqa"))
m05 <- lmer(log(Day14Mass) ~ NetManipulation + ChickSwappedYN + Year +
                        JulianDay14 + ChickSex + (1|OriginNestID) +
                        (1|RearNestID),
                        data=dat, na.action=na.exclude, REML=FALSE,
                        control = lmerControl(optimizer = "bobyqa"))
m06 <- lmer(Day14Mass ~ NetManipulation + ChickSwappedYN + JulianDay14 +
```

```
                        Year + ChickSex + (1|OriginNestID) + (1|RearNestID),
                        data=dat.OR, na.action=na.exclude, REML=FALSE,
                        control = lmerControl(optimizer = "bobyqa"))

# ------------------------------------------------------------------------------
# Models according to Type U Decisions (Uncertain Equivalence) ---------------
m07 <- lmer(Day14Mass ~ NetManipulation + ChickSwappedYN + JulianDay14 +
                        Year + ChickSex + PreTreatBrSize + (1|OriginNestID) +
                        (1|RearNestID),
                        data=dat, na.action=na.exclude, REML=FALSE,
                        control = lmerControl(optimizer = "bobyqa"))
m08 <- lmer(Day14Tarsus ~ NetManipulation + ChickSwappedYN + JulianDay14 +
                        Year + ChickSex + (1|OriginNestID) + (1|RearNestID),
                        data=dat, na.action=na.exclude, REML=FALSE,
                        control = lmerControl(optimizer = "bobyqa"))

# ------------------------------------------------------------------------------
# Models according to Type N Decisions (Principled Nonequivalence) -----------
m09 <- lmer(SurvivalProp ~ NetManipulation + Year + JulianDay14 +
                        (1|RearNestID), data=dat.Survival,
                        weights=PostTreatBrSize, control=lmerControl(
                        check.nobs.vs.nlev='ignore',
                        check.nobs.vs.nRE='ignore', optimizer='bobyqa'))
m10 <- lmer(Day14Mass ~ NetManipulation + ChickSwappedYN + JulianDay14 +
                        Year + ChickSex + PostTreatBrSize + (1|OriginNestID) +
                        (1|RearNestID),
                        data=dat, na.action=na.exclude,REML=FALSE,
                        control = lmerControl(optimizer = "bobyqa"))
m11 <- lmer(Day14Mass ~ NetManipulation + ChickSwappedYN + JulianDay14 +
                        Year + ChickSex + EarlyMortality + (1|OriginNestID) +
                        (1|RearNestID),
                        data=dat, na.action=na.exclude, REML=FALSE,
                        control = lmerControl(optimizer = "bobyqa"))
m12 <- lmer(Day14Mass ~ NetManipulation + ChickSwappedYN + JulianDay14 +
                        Year + ChickSex + Day14BrSize + (1|OriginNestID) +
                        (1|RearNestID),
                        data=dat, na.action=na.exclude, REML=FALSE,
                        control = lmerControl(optimizer = "bobyqa"))
m13 <- lmer(Day14Mass ~ NetManipulation + ChickSwappedYN + JulianDay14 +
                        Year + ChickSex + LateMortality + (1|OriginNestID) +
                        (1|RearNestID),
                        data=dat, na.action=na.exclude, REML=FALSE,
                        control = lmerControl(optimizer = "bobyqa"))
m14 <- lmer(Day14Mass ~ NetManipulation + ChickSwappedYN + JulianDay14 +
                        Year + ChickSex + NFledglings + (1|OriginNestID) +
                        (1|RearNestID),
                        data=dat, na.action=na.exclude, REML=FALSE,
                        control = lmerControl(optimizer = "bobyqa"))
m15 <- lmer(Day14Mass ~ NetManipulation + ChickSwappedYN + JulianDay14 +
                        Year + ChickSex + Day14Tarsus + (1|OriginNestID) +
                        (1|RearNestID),
```

```
                           data=dat, na.action=na.exclude, REML=FALSE,
                           control = lmerControl(optimizer = "bobyqa"))
m16 <- lmer(Day14Tarsus ~ NetManipulation + ChickSwappedYN + JulianDay14 +
                           Year + ChickSex + Day14Mass +
                           (1|OriginNestID) + (1|RearNestID),
                           data=dat, na.action=na.exclude, REML=FALSE,
                           control = lmerControl(optimizer = "bobyqa"))
m17 <- lmer(Day14Mass ~ NetManipulation + ChickSwappedYN + JulianDay14 +
                           Year + ChickSex + NChicksAdded + (1|OriginNestID) +
                           (1|RearNestID),
                           data=dat, na.action=na.exclude, REML=FALSE,
                           control = lmerControl(optimizer = "bobyqa"))
m18 <- lmer(Day14Mass ~ NetManipulation + ChickSwappedYN + JulianDay14 +
                           Year + ChickSex + NChicksRemoved + (1|OriginNestID) +
                           (1|RearNestID),
                           data=dat, na.action=na.exclude, REML=FALSE,
                           control = lmerControl(optimizer = "bobyqa"))

# ------------------------------------------------------------------------------
summary(m01) ; summary(m02) ; summary(m03) ; summary(m04) ; summary(m05) ;
summary(m06) ; summary(m07) ; summary(m08) ; summary(m09) ; summary(m10) ;
summary(m11) ; summary(m12) ; summary(m13) ; summary(m14) ; summary(m15) ;
summary(m16) ; summary(m17) ; summary(m18)
mods <- list(m01,m02,m03,m04,m05,m06,m07,m08,m09,
             m10,m11,m12,m13,m14,m15,m16,m17,m18)
names(mods) <- c("m01","m02","m03","m04","m05","m06","m07","m08","m09",
                 "m10","m11","m12","m13","m14","m15","m16","m17","m18")

# ------------------------------------------------------------------------------
# For each model, extract Zr-values and their SE -----------------------------
for(j in 1:18) {
  mod <- mods[[j]]
  out$analysis[k] <- SubsetData
  out$model[k] <- names(mods)[j]
  out$type[k] <- ifelse(names(mods)[j] %in% c("m01","m02","m03",
                                              "m04","m05","m06"), "E",
                 ifelse(names(mods)[j] %in% c("m07","m08"), "U", "N"))
  out$b[k] <- summary(mod)$coef["NetManipulation", "Estimate"]
  out$SE[k] <- summary(mod)$coef["NetManipulation", "Std. Error"]
  out$t[k] <- summary(mod)$coef["NetManipulation", "t value"]
  out$df[k] <- summary(mod)$coef["NetManipulation", "df"]

  # Convert the t-value and the degree of freedom (df) to r
  r <- sqrt(out$t[k]^2 / (out$t[k]^2 + out$df[k]))
  r <- ifelse(summary(mod)$coef["NetManipulation", "Estimate"] < 0, -r, r)

  # Convert r and accompanying df to Zr and its sampling variance 1/(n-3)
  out$Zr[k] <- atanh(r)
  out$SEZr[k] <- sqrt(1 / (out$df[k] + 1 - 3))
  k <- k + 1
}
```

```r
}

# ------------------------------------------------------------------------------
# Plot model outputs ----------------------------------------------------------
COLS <- brewer.pal(n=9, name="Set1")
COLS <- COLS[c(1,2,6)]
out$COL <- ifelse(out$type=="E", COLS[2],
           ifelse(out$type=="U", COLS[3],
           ifelse(out$type=="N", COLS[1], NA)))

# -------------------
# Zr and their SE ---
svg(filename=paste(path,"Fig3.svg",sep=""), height=80/25.4, width=150/25.4,
    family="Arial", pointsize=10)
par(mar=c(2, 2.2, 1.5, 0.2), mgp=c(1.2, 0.2, 0))

plot(x=c(0,(nrow(out)+3)), y=range(c(out$Zr-out$SEZr, out$Zr+out$SEZr)),
     type="n", ylab="Effect size Zr + SE", xlab="", tcl=-0.25,
     xaxt="n", xaxs="i")

abline(h=0, lty=2, lwd=1)

out1 <- subset(out, analysis=="CRE")
out2 <- subset(out, analysis=="CE")
out3 <- subset(out, analysis=="CR")

abline(v=c(nrow(out1)+1,2*(nrow(out1)+1)))

k <- 1
for(i in 1:nrow(out1)) {
  lines(x=c(k,k), y=c(out1$Zr[i]-out1$SEZr[i],out1$Zr[i]+out1$SEZr[i]),
        lwd=1, lty=1)
  points(x=k, y=out1$Zr[i], pch=21, bg=out1$COL[i])
  k <- k + 1
}

k <- k + 1
for(i in 1:nrow(out2)) {
  lines(x=c(k,k), y=c(out2$Zr[i]-out2$SEZr[i],out2$Zr[i]+out2$SEZr[i]),
        lwd=1, lty=1)
  points(x=k, y=out2$Zr[i], pch=21, bg=out2$COL[i])
  k <- k + 1
}

k <- k + 1
for(i in 1:nrow(out3)) {
  lines(x=c(k,k), y=c(out3$Zr[i]-out3$SEZr[i],out3$Zr[i]+out3$SEZr[i]),
        lwd=1, lty=1)
  points(x=k, y=out3$Zr[i], pch=21, bg=out3$COL[i])
  k <- k + 1
}
```

```r
dev.off()

# --------------------------
# Estimates and their SE ---
out$b <- ifelse(out$model %in% c("m05","m08","m09","m16"), NA, out$b)
out$SE <- ifelse(out$model %in% c("m05","m08","m09","m16"), NA, out$SE)

svg(filename=paste(path,"FigS3.svg",sep=""), height=80/25.4, width=150/25.4,
    family="Arial", pointsize=10)

par(mar=c(2, 2.2, 1.5, 0.2), mgp=c(1.2, 0.2, 0))

plot(x=c(0,(nrow(out)+3)), y=range(c(out$b-out$SE, out$b+out$SE), na.rm=TRUE),
     type="n", ylab="Effect size (g / chick) + SE", xlab="", tcl=-0.25,
     xaxt="n", xaxs="i")

abline(h=0, lty=2, lwd=1)

out1 <- subset(out, analysis=="CRE")
out2 <- subset(out, analysis=="CE")
out3 <- subset(out, analysis=="CR")

abline(v=c(nrow(out1)+1,2*(nrow(out1)+1)))

k <- 1
for(i in 1:nrow(out1)) {
  if(!is.na(out1$b[i])) {
    lines(x=c(k,k), y=c(out1$b[i]-out1$SE[i],out1$b[i]+out1$SE[i]),
          lwd=1, lty=1)
    points(x=k, y=out1$b[i], pch=21, bg=out1$COL[i])
    k <- k + 1 } else { k <- k + 1 }
}
k <- k + 1
for(i in 1:nrow(out2)) {
  if(!is.na(out2$b[i])) {
    lines(x=c(k,k), y=c(out2$b[i]-out2$SE[i],out2$b[i]+out2$SE[i]),
          lwd=1, lty=1)
    points(x=k, y=out2$b[i], pch=21, bg=out2$COL[i])
    k <- k + 1 } else { k <- k + 1 }
}
k <- k + 1
for(i in 1:nrow(out3)) {
  if(!is.na(out3$b[i])) {
    lines(x=c(k,k), y=c(out3$b[i]-out3$SE[i],out3$b[i]+out3$SE[i]),
          lwd=1, lty=1)
    points(x=k, y=out3$b[i], pch=21, bg=out3$COL[i])
    k <- k + 1 } else { k <- k + 1 }
}

dev.off()
```

---

#### 3.2 Figures S4–S6

We visualized the effects of these analytical decisions using three complementary effect-size representations: treatment effects on chick mass expressed in grams per chick (**Fig. S4**), the corresponding analyses after exchanging the roles of mass and tarsus length (**Fig. S5**), and analyses in which treatment was coded as a two-level factor contrasting enlarged and reduced broods (**Fig. S6**).

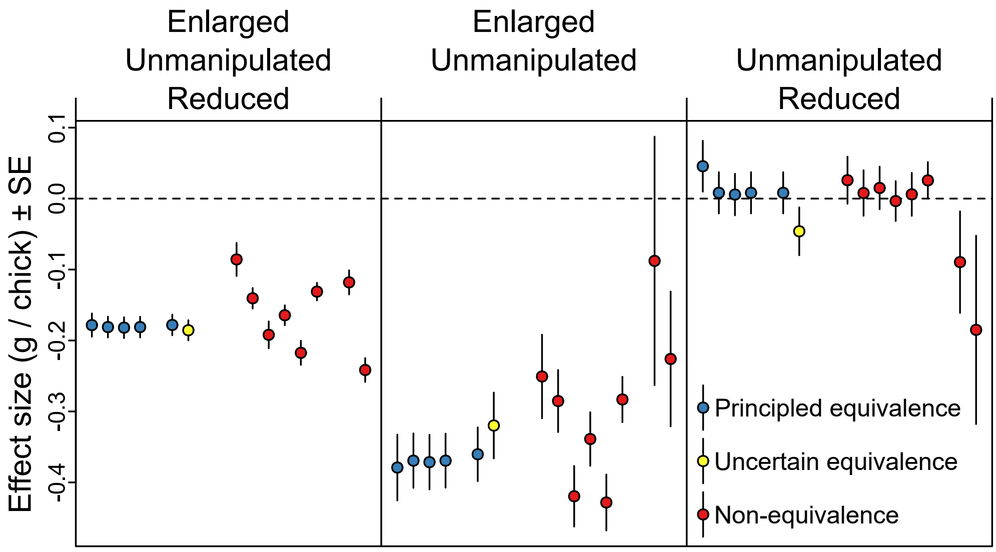

***Fig. S4*** *| Effects of analytical decisions on estimated treatment effects in the 'blue tit dataset' from Gould et al. (2025). Effect size estimates represent the impact of net brood size manipulation on chick body mass, expressed as the change in body mass (grams) per chick. The three panels show analyses based on all broods (left), enlarged and unmanipulated broods only (centre), and reduced and unmanipulated broods only (right). Colours indicate our classification of analytical decisions: blue = principled equivalence, yellow = uncertain equivalence, red = principled non-equivalence. Models using tarsus length, log-transformed mass, or fledging success as outcome variables are omitted because their coefficients are not directly interpretable as changes in body mass in grams. The full-range estimate is intermediate between the negative effect in enlarged broods and the near-absent effect in reduced broods. Note that this intuitive result looks quite different in* ***Fig. 4****, where effect sizes are assessed more in terms of statistical significance (magnitude of t-values), than in terms of an absolute effect on body mass.*

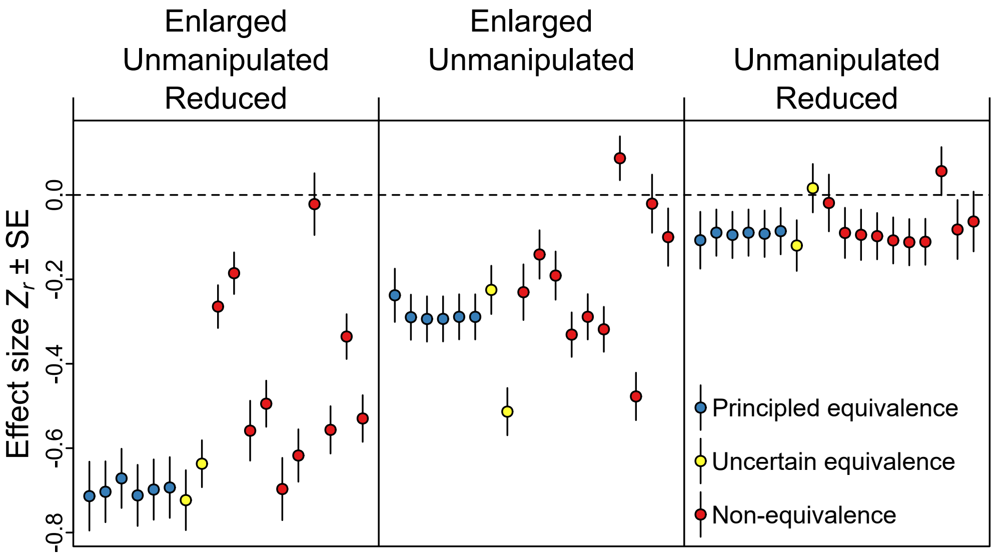

***Fig. S5*** *| Robustness of the analytical-decision framework after exchanging the roles of Day 14 chick body mass and Day 14 tarsus length. The same model set as in* ***Fig. S4*** *was refitted after systematically substituting mass for tarsus length and vice versa. Colours indicate our classification of analytical decisions: blue = principled equivalence, yellow = uncertain equivalence, red = principled non-equivalence. This analysis evaluates whether the qualitative conclusions depend on how chick growth is operationalized.*

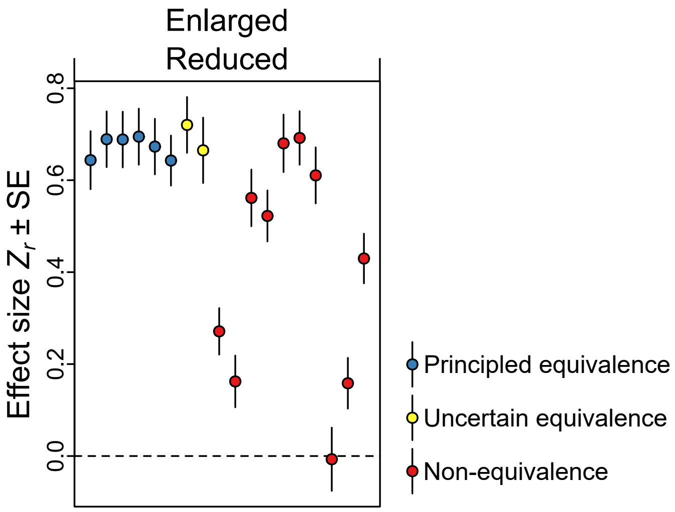

***Fig. S6*** *| Effects of analytical decisions when brood-size manipulation is operationalized as a two-level factor contrasting enlarged and reduced broods directly. This specification avoids using unmanipulated broods as a reference group, but changes the estimand to the difference between opposite manipulation directions. Colours indicate our classification of analytical decisions: blue = principled equivalence, yellow = uncertain equivalence, red = principled non-equivalence.*

---

### 4. Tutorial and code in R

#### 4.1 Preparations

We have implemented all code in R (R Core Team 2024). The R code is highlighted in grey boxes, and to run it, you must copy and paste it into the R console. Red boxes highlight the R console output that is essential for understanding of this tutorial.

We will use the metafor package in R (Viechtbauer 2010) to perform random-effects meta-analyses. If you do not have metafor installed, the following code will install the package and load it into R. If metafor is already installed, the code will simply load the package.

```
rm(list=ls())
if("metafor" %in% rownames(installed.packages()) == FALSE)
  { install.packages("metafor") }
require(metafor)
```

---

#### 4.2 Data acquisition for meta-analysis

Imagine that we have extracted **effect size estimates (`EF` in the code below) along with their standard errors (`SE`) from $N = 20$ studies**. In a later section, we will provide an `R` function to simulate such data, but for now, we will use an example dataset. The data might look like this:

```
EF <- c(-0.56626122,-0.26370324, 0.14872448,-0.05425463,0.18319571, 0.13952246,
         0.22050781,-0.22641049, 0.07724746, 0.11124586,0.15866421,-0.09323908,
        -0.17886749,-0.04861292,-0.33830514, 0.23966551,0.40206087,-0.22768412,
        -0.15233253,-0.13635335)
SE <- c(0.1622214, 0.1622214, 0.1666667, 0.1524986, 0.1666667, 0.1490712,
        0.1825742, 0.1581139, 0.1507557, 0.1825742, 0.1428571, 0.1507557,
        0.1690309, 0.1825742, 0.1666667, 0.1961161, 0.1740777, 0.1581139,
        0.1581139, 0.1924501)
```

We calculate the **lower limits of the 95% confidence intervals (`CI95.L`)**.

```
CI95.L <- EF - qnorm(0.975) * SE
```

We calculate the **upper limits of the 95% confidence intervals (`CI95.U`)**.

```
CI95.U <- EF + qnorm(0.975) * SE
```

We **create a dataframe (`meta.data`)** from the estimates of the 20 studies and take a look at it.

```
(meta.data <- data.frame(EF=EF, SE=SE, CI95.L=CI95.L, CI95.U=CI95.U))
```

**R console output**

```
            EF        SE      CI95.L      CI95.U
1  -0.56626122 0.1622214 -0.88420932 -0.24831312
2  -0.26370324 0.1622214 -0.58165134  0.05424486
3   0.14872448 0.1666667 -0.17793625  0.47538521
4  -0.05425463 0.1524986 -0.35314639  0.24463713
5   0.18319571 0.1666667 -0.14346502  0.50985644
6   0.13952246 0.1490712 -0.15265172  0.43169664
7   0.22050781 0.1825742 -0.13733105  0.57834667
8  -0.22641049 0.1581139 -0.53630804  0.08348706
9   0.07724746 0.1507557 -0.21822828  0.37272320
10  0.11124586 0.1825742 -0.24659300  0.46908472
11  0.15866421 0.1428571 -0.12133056  0.43865898
12 -0.09323908 0.1507557 -0.38871482  0.20223666
13 -0.17886749 0.1690309 -0.51016197  0.15242699
14 -0.04861292 0.1825742 -0.40645178  0.30922594
15 -0.33830514 0.1666667 -0.66496587 -0.01164441
16  0.23966551 0.1961161 -0.14471498  0.62404600
17  0.40206087 0.1740777  0.06087485  0.74324689
18 -0.22768412 0.1581139 -0.53758167  0.08221343
19 -0.15233253 0.1581139 -0.46223008  0.15756502
20 -0.13635335 0.1924501 -0.51354861  0.24084191
```

#### 4.3 Perform a random-effects meta-analysis

Using these data, we **perform a random-effects meta-analysis**. Note that the `rma()` function requires only the arguments 'yi' (effect size estimates `EF`) and 'sei' (their standard errors `SE`).

The 'method' argument specifies one of the available estimators for the amount of heterogeneity. By setting 'method="DL"', we choose the DerSimonian-Laird estimator (DerSimonian & Laird 1986, Raudenbush 2009). Setting the 'method' argument to "DL" is equivalent to using the `rmeta` package in `R` with the command `rmeta::meta.summaries(meta.data$EF, meta.data$SE, method="random")` (Lumley 2018).

```
require(metafor)
meta.out <- rma(yi=meta.data$EF, sei=meta.data$SE, method="DL")
```

We **examine the output of the meta-analysis (`meta.out`)** and **create a forest plot**.

```
meta.out
forest(meta.out)
```

**R console output**

```
Random-Effects Model (k = 20; tau^2 estimator: DL)

tau^2 (estimated amount of total heterogeneity): 0.0274 (SE = 0.0177)
tau (square root of estimated tau^2 value):      0.1656
I^2 (total heterogeneity / total variability):   50.33%
H^2 (total variability / sampling variability):  2.01

Test for Heterogeneity:
Q(df = 19) = 38.2540, p-val = 0.0055

Model Results:

estimate      se     zval     pval    ci.lb   ci.ub
 -0.0341  0.0524  -0.6512  0.5149  -0.1367  0.0685

---
Signif. codes:  0 '***' 0.001 '**' 0.01 '*' 0.05 '.' 0.1 ' ' 1
```

Here we see (1) the meta-analysis mean effect ($m = -0.0341$) along with its standard error ($se = 0.0524$) and 95% confidence interval ($-0.1367$ *to* $0.0685$). (2) We also obtain different estimates of heterogeneity: $\tau^2 = 0.0274$, $I^2 = 0.5033$, and $Q = 38.2540$. We can retrieve and **save these values into `R`'s memory**.

```
# Meta-analytic mean effect
mean.effect <- meta.out$beta[1]
# Standard error of the meta-analytic mean effect
mean.effect.se <- meta.out$se
# Lower and upper 95% confidence limit
mean.effect.ci.l <- meta.out$ci.lb
mean.effect.ci.u <- meta.out$ci.ub
# Tau2
het.tau2 <- meta.out$tau2
# I2. This is provided as a percentage,
# but we need it as a proportion for further calculations
het.i2 <- meta.out$I2 / 100
# Cochran's Q
het.q <- meta.out$QE
```

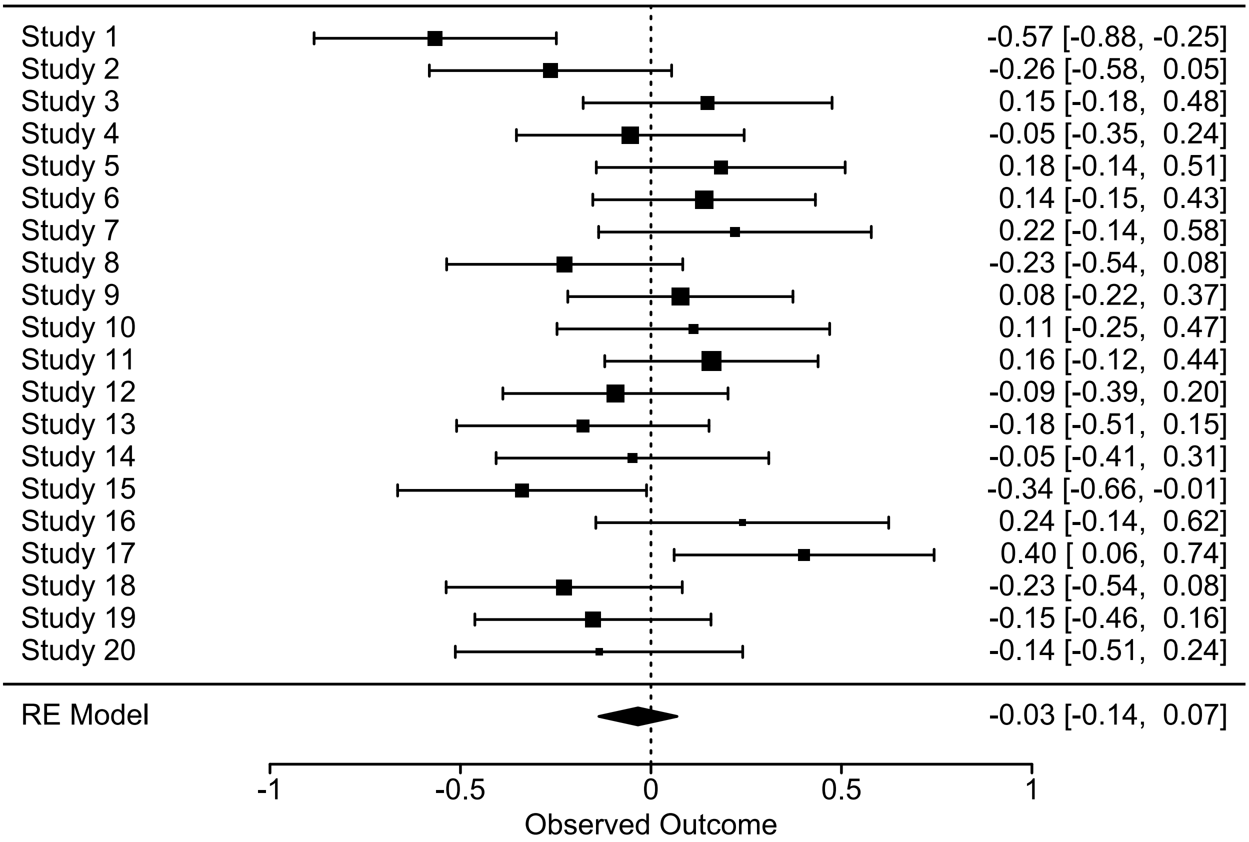

***Fig. S7*** | *Forest plot of the effect sizes from the $N = 20$ studies, including their conventional 95% confidence intervals. The mean effect from the meta-analysis is shown at the bottom. Note that the 95% confidence intervals of two individual studies do not encompass the meta-analytic mean effect. This corresponds to a type I error rate of 10%, which means that 90% of all studies cover the mean effect from the meta-analysis within their conventional 95% confidence intervals. We obtain these values using the following code.*

```
sum((meta.data$CI95.L < mean.effect & meta.data$CI95.U < mean.effect) |
    (meta.data$CI95.L > mean.effect & meta.data$CI95.U > mean.effect))
nrow(meta.data)
```

---

#### 4.4 Extend the 95% confidence intervals to 95% $\text{CI}_{\text{ext}}$

##### 4.4.1 Adjust the 95% confidence intervals

We have observed considerable heterogeneity among individual study estimates, which may suggest that the conventional 95% confidence intervals are too narrow. How much wider do they need to be to maintain a type I error rate of 5%? To answer this question, we need to calculate the **correction factor** $y$.

```
y <- sqrt(1 / (1 - het.i2))
```

**R console output**

```
> y
[1] 1.418932
```

What is the **critical** $Z$**-value given the observed heterogeneity (`Z.c`)**? Knowing the heterogeneity-aware critical $Z$-value allows us to calculate the expected type I error rate. Note that `qnorm(0.025)` and `qnorm(0.975)` provide the exact values for the 2.5th and 97.5th percentiles of the standard normal distribution, often approximated as -1.96 and 1.96, respectively. $Z$-scores less than -1.96 or greater than 1.96 are considered 'significant' when using a two-tailed test with an $\alpha$ threshold of 0.05.

```
Z.c <- qnorm(0.975) / y
```

**R console output**

```
> Z.c
[1] 1.381295
```

The **expected type I error rate (`Type.I.rate`)** is calculated as follows. The `pnorm()` function implements the cumulative distribution function of the standard normal distribution $\Phi(x)$. It returns the area under the normal distribution up to a given $Z$-score. We multiply by 2 because we are conducting a two-tailed test.

```
Type.I.rate <- 2 * (1 - pnorm(Z.c))
```

**R console output**

```
> Type.I.rate
[1] 0.1671882
```

Thus, instead of the desired type I error rate of 0.05, the expected type I error rate is 0.17. As a result, we would **expect the proportion of individual study confidence intervals that cover the meta-analytic mean effect (`Exp.Prop.CI.Beta`)** to decrease from 0.95 to a lower value.

```
Exp.Prop.CI.Beta <- 1 - Type.I.rate
```

R console output

```
> Exp.Prop.CI.Beta
[1] 0.8328118
```

We expect that 83% of the confidence intervals of the individual studies will cover the meta-analytic mean effect. This can be compared to the observed value of 90% calculated above. The discrepancy may arise from the small sample size of $N = 20$ studies and the resulting sampling noise.

To bring the type I error rate back to the desired 5%, we need to **extend the 95% confidence intervals (or the standard errors)** of the individual studies by factor $y$. We refer to these extended 95% confidence intervals as 95% $\text{CI}_{\text{ext}}$.

```
require(metafor)
meta.data$extSE <- meta.data$SE * y
meta.out.ext <- rma(yi=meta.data$EF, sei=meta.data$extSE, method="DL")
meta.out.ext
forest(meta.out.ext)
```

R console output

```
Random-Effects Model (k = 20; tau^2 estimator: DL)

tau^2 (estimated amount of total heterogeneity): 0 (SE = 0.0177)
tau (square root of estimated tau^2 value):      0
I^2 (total heterogeneity / total variability):   0.00%
H^2 (total variability / sampling variability):  1.00

Test for Heterogeneity:
Q(df = 19) = 19.0000, p-val = 0.4568

Model Results:

estimate      se     zval    pval    ci.lb   ci.ub
 -0.0364  0.0522  -0.6984  0.4849  -0.1387  0.0658

---
Signif. codes:  0 ‘***’ 0.001 ‘**’ 0.01 ‘*’ 0.05 ‘.’ 0.1 ‘ ’ 1
```

Note that the heterogeneity is $I^2 = 0$, and Cochran's $Q = 19$ (with a $P$-value as close to 0.5 as possible given this sample size). We will see below that for $k = 20$ studies, Cochran's $Q = 19$ translates exactly to $I^2 = 0$.

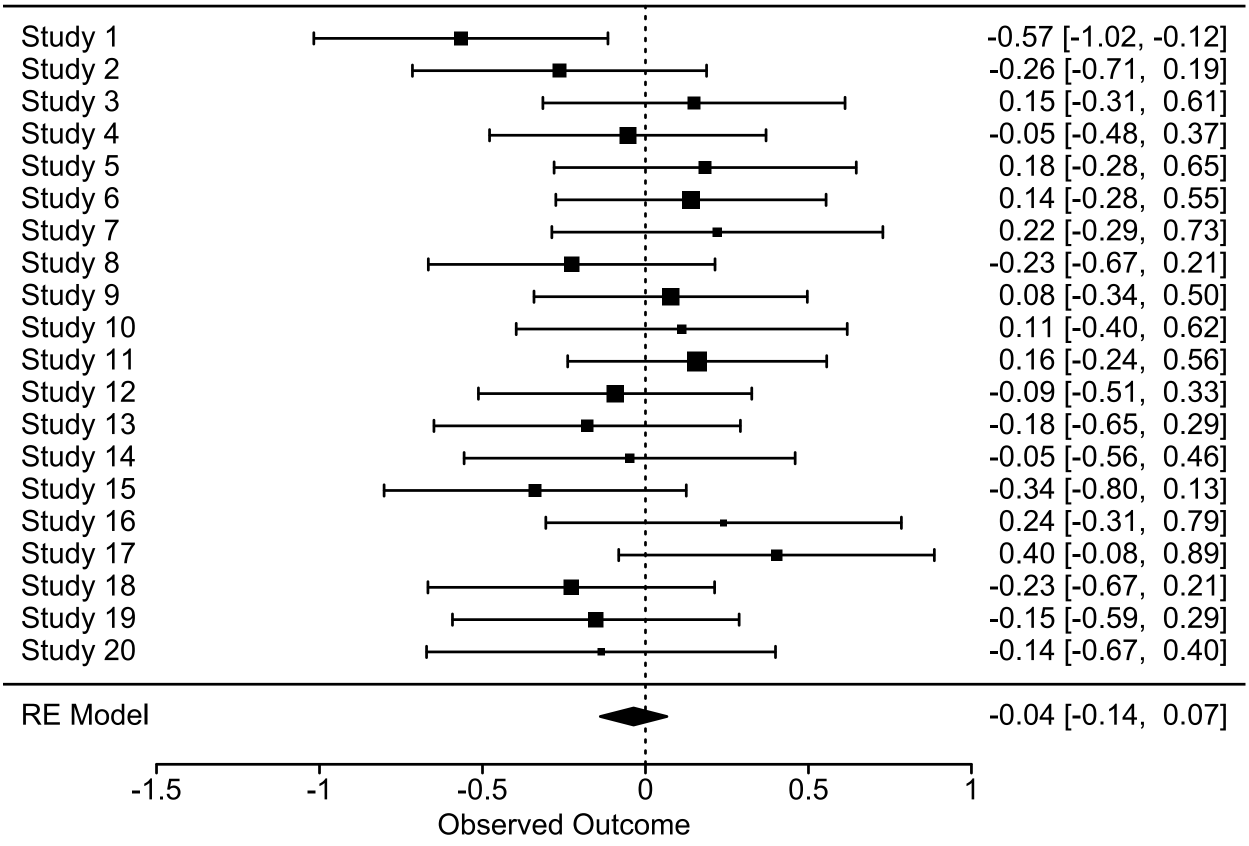

***Fig. S8*** | *Forest plot of the effect sizes from the $N = 20$ studies with their extended 95% confidence intervals. The mean effect from the meta-analysis is shown at the bottom. Note that the 95% confidence interval for a single individual study does not cover the meta-analytic mean effect. This corresponds to a type I error rate of 5%, which is the expected value we aimed for.*

##### 4.4.2 Adjust the $\alpha$ threshold

If we want to maintain the type I error rate of our individual study at 5%, we need to **increase the critical $Z$-value by factor $y$ (`Z.c.ext`)**.

```
Z.c.ext <- qnorm(0.975) * y
```

**R console output**

```
> Z.c.ext
[1] 2.781055
```

Finally, we **calculate the corresponding $\alpha$ threshold** (i.e., the threshold at which we declare a $P$-value significant). For an explanation of the formula, see the calculation of the type I error rate above.

```
alpha.ext <- 2 * (1 - pnorm(Z.c.ext))
```

**R console output**

```
> alpha.ext
[1] 0.005418251
```

Thus, the critical $P$-value needed to maintain type I errors at 5% is 0.0054, which is nearly ten times smaller than the conventional threshold of 0.05.

---

#### 4.5 Derive $I^2_{MAMA}$ from $I^2*$

Now suppose that the $N = 20$ effect size estimates do not originate from individual studies but from a **many-analyses study in which each research team analysed exactly the same dataset**. Consequently, there is no variance in true study system effects, ${\sigma_A}^2$. The conventional formula used in the `metafor` `R` package to derive $I^2$ is incorrect because it assumes the presence of true sampling noise, ${\sigma_B}^2$. Therefore, we denote the $I^2$ derived from `metafor::rma()` as $I^2*$ (and in the code below as `het.i2s`).

```
require(metafor)
meta.out <- rma(yi=meta.data$EF, sei=meta.data$SE, method="DL")
# I2*
het.i2s <- meta.out$I2 / 100
# Cochran's Q
het.q <- meta.out$QE
```

**R console output**

```
> het.i2s
[1] 0.5033197
```

We can **calculate the correct $I^2_{MAMA}$ (`het.i2MAMA`)** from $I^2*$.

```
het.i2MAMA <- 1 / (2 - het.i2s)
```

**R console output**

```
> het.i2MAMA
[1] 0.6681454
```

In this case, $I^2_{MAMA}$ is substantially higher than $I^2*$ (0.67 versus 0.50, respectively). If $I^2*$ can only take values between 0 and 1, we see that $I^2_{MAMA}$ is bounded between 0.5 and 1, which is an undesirable property. We can also **estimate $I^2*$ from Cochran's $Q$ (`het.i2s.Q`)** and the number of studies included in the meta-analysis, denoted as $k$.

```
k <- nrow(meta.data)
het.i2s.Q <- (het.q - k + 1) / het.q
# Compare both ways of estimating het.i2s (rounding to the 10th decimal)
round(het.i2s.Q,10) == round(het.i2s,10)
```

Using the alternative estimation procedure for $I^2*$ based on Cochran's $Q$, we find that $I^2*$ can now take negative values (if $Q$ is small or $k$ is large). In fact, $I^2_{MAMA}$ is now defined within the interval between 0 and 1. Thus, we can **derive $I^2_{MAMA}$ (`het.i2MAMA`) using the following code**.

```r
require(metafor)
meta.out <- rma(yi=meta.data$EF, sei=meta.data$SE, method = "DL")
# Cochran's Q
het.q <- meta.out$QE
# Number of individual research teams providing effect size estimates
k <- nrow(meta.data)
# Estimation of het.i2s through Cochran's Q
het.i2s.Q <- (het.q - k + 1) / het.q
# Calculate i2MAMA
het.i2MAMA <- 1 / (2 - het.i2s.Q)
```

---

##### 4.6 Derive $\mathrm{SE}_{\mathrm{MAMA}}$

In a conventional random-effects meta-analysis, the formula used in the `metafor` `R` package to calculate the **standard error of the meta-analytic mean effect ($\mathrm{SE}_{\mathrm{META}}$ and in the code below `SE.META`)** is given by (Borenstein *et al.* 2021):

$$SE_{META} = \sqrt{\frac{1}{\sum 1/({\sigma_A}^2+{\sigma_B}^2+{\sigma_C}^2)}}$$

If each individual study has the same sample size, applying Fisher's $z$-transformation stabilizes the variance, such that the within-study variance ${\sigma_B}^2$ (and thus the standard error) are expected to be approximately identical across individual studies. Then the equation simplifies to:

$$SE_{META} = \sqrt{\frac{{\sigma_A}^2+{\sigma_B}^2+{\sigma_C}^2}{N}}$$

where $N$ is the number of individual studies included (in this case, $N = 20$) .

```r
SE.META <- meta.out$se
```

**R console output**

```
> SE.META
[1] 0.05236626
```

In a many-analyes meta-analysis (MAMA), the standard error $\mathrm{SE}_{\mathrm{META}}$ is misleadingly small because it assumes that each study is based on independent data. However, in a MAMA, all research teams analyse the same dataset. As a result, the variance due to the choice of analysis ${\sigma_C}^2$ is reduced (and ${\sigma_A}^2 = 0$), while the sampling noise ${\sigma_B}^2$ is treated as a representative within-study sampling variance. The **correct standard error for the MAMA ($\mathrm{SE}_{\mathrm{MAMA}}$)** is therefore given by:

$$SE_{MAMA} = \sqrt{{\sigma_B}^2 + \frac{{\sigma_C}^2}{N}}$$

where $N$ is the number of research teams that provided effect size estimates (in this case, $N = 20$). We need estimates for ${\sigma_B}^2$ and ${\sigma_C}^2$. In a MAMA, we know that ${\sigma_C}^2 = {\sigma_T}^2$. Therefore, we use the variance in effect size estimates provided by the $N = 20$ research teams as our estimate for ${\sigma_C}^2$ (`sigmaC2`).

```r
sigmaC2 <- var(meta.data$EF)
```

**R console output**

```
> sigmaC2
[1] 0.05597902
```

We can estimate the representative within-study sampling variance ${\sigma_B}^2$ (`sigmaB2`) from $I^2*$ and ${\sigma_C}^2$ with the formula $I^2* = \frac{{\sigma_T}^2-{\sigma_B}^2}{{\sigma_T}^2}$.

```r
require(metafor)
meta.out <- rma(yi=meta.data$EF, sei=meta.data$SE, method="DL")
# Cochran's Q
het.q <- meta.out$QE
# Number of individual research teams providing effect size estimates
k <- nrow(meta.data)
# Estimation of het.i2s through Cochran's Q
het.i2s.Q <- (het.q - k + 1) / het.q
# Estimate sigmaB2
sigmaB2 <- sigmaC2 * (1 - het.i2s.Q)
```

**R console output**

```
> sigmaB2
[1] 0.02780368
```

Now that we have estimates for ${\sigma_B}^2$, ${\sigma_C}^2$, and we know the number of research teams $N$, we can **calculate the many-analyses standard error of the meta-analytic mean effect, $\mathrm{SE_{MAMA}}$ (`SE.MAMA`)**, using the following formula:

```r
SE.MAMA <- sqrt(sigmaB2 + sigmaC2/nrow(meta.data))
```

**R console output**

```
> SE.MAMA
[1] 0.1749361
```

We find that $\mathrm{SE_{MAMA}}$ is more than three times larger than $\mathrm{SE_{META}}$ (0.17 versus 0.052, respectively). We will simulate new effect size data below and use these simulations to demonstrate that $\mathrm{SE_{MAMA}}$ can be employed to calculate precise 95% confidence intervals for the many-analyses mean effect size.

---

#### 4.7 Function to simulate new effect size data

For the more experienced R user, we provide a **function that simulates effect size data** that can then be used to perform a meta-analysis. The function allows the user to vary:

(1) The number of studies included ('N').

(2) The sample size per study ('M').

(3) Whether all studies should have the same sample size or whether there should be variation in sample size between them ('Dist.M' = c("constant","variable exact pois","variable exact nbin")).

(4) If 'Dist.M = "variable exact nbin" ' is selected, the amount of variance in sample size between studies, specifically the shape parameter 'theta' of a negative binomial distribution.

(5) The expected meta-analytic mean effect ('MU.EF'), which should fall within the interval between -1 and 1.

(6) The amount of heterogeneity ('Het.I2').

We **define the function `sim.meta.EF(N, M, Dist.M, theta, MU.EF, Het.I2)` to simulate the effect size data**. This function returns a dataframe similar to the one described above, with additional columns: 'N', 'M', 'Het.I2'. These columns represent a running number for each study $(1, 2, \ldots, N)$, the sample size $M$ for each study, and the expected heterogeneity $I^2$, respectively.

The effect size data is simulated by applying Fisher's $z$-transformation (also known as the inverse hyperbolic tangent or the area tangens hyperbolicus) to the Pearson correlation coefficient. This transformation maps the open interval (-1, 1) onto the real number line.

Fisher's $z$-transformed Pearson correlation coefficients (using the `atanh()` function) can be back-transformed to Pearson's $r$ using the `tanh()` function. However, we follow a more precise method for deriving Fisher's $z$-transformed Pearson correlation coefficients, as outlined by Vrbik (2005). Due to this approach, direct back-transformation is more complex. We calculate their standard errors using the function provided in Vrbik (2005).

Note that the code extends over two pages and should be combined into a single block for execution.

```
sim.meta.EF <- function(N=20, M=40, Dist.M="variable exact pois",
                        theta=10, MU.EF=0, Het.I2=0) {
  # Calculate the extension factor y
  y <- sqrt(1 / (1 - Het.I2))
  # Create a dataframe for all N studies given a sample size M per study
  meta.data <- data.frame(matrix(rep(NA,N*7),ncol=7))
  colnames(meta.data) <- c("N","M","Het.I2","EF","SE","CI95.L","CI95.U")
  # Individual studies N
  meta.data$N <- 1:nrow(meta.data)
  # Individual study sample sizes M
  if(Dist.M=="constant") { meta.data$M <- rep(M,N) }
  if(Dist.M=="variable exact pois") {
    Sample.M <- c(table(sample(x=c(1:N), size=M*N, replace=TRUE)))
    Sample.M <- c(Sample.M,rep(0,N))
    meta.data$M <- Sample.M[1:N]
  }
  if(Dist.M=="variable exact nbin") {
    theta=theta; w=rgamma(N,shape=theta,scale=1/theta)
```

```r
    Sample.M <- c(table(sample(x=c(1:N), size=M*N, replace=TRUE, prob=w)))
    Sample.M <- c(Sample.M,rep(0,N))
    meta.data$M <- Sample.M[1:N]
  }
  # Expected heterogeneity I2
  meta.data$Het.I2 <- Het.I2
  # Calculate Fisher's z (inverse hyperbolic tangent) for the N studies
  # according to Vrbik (2005)
  z <- atanh(MU.EF) + (MU.EF / (2 * meta.data$M))
  # Calculate the variance of Fisher's z for the N studies
  # according to Vrbik (2005)
  var.z <- (1/meta.data$M) + (6 - MU.EF^2) / (2 * meta.data$M^2)
  sd.z <- sqrt(var.z)
  # Simulate Fisher's z, given I2
  sim.z <- data.frame(M=1, z=z, sd.z=sd.z)
  meta.data$EF <- apply(sim.z, 1,
                        function(x) rnorm(x["M"], mean=x["z"], sd=x["sd.z"]*y))
  meta.data$SE <- sd.z
  meta.data$CI95.L <- meta.data$EF - qnorm(0.975) * meta.data$SE
  meta.data$CI95.U <- meta.data$EF + qnorm(0.975) * meta.data$SE
  return(meta.data)
}
```

We can **use this function to simulate new data for a meta-analysis**. We have generated the dataframe above with the following arguments. Note that we cannot reproduce the exact numbers due to the random sampling procedure implemented in `sim.meta.EF()`.

```r
require(metafor)
meta.data <- sim.meta.EF(N=20, M=40, Dist.M="variable exact pois", theta=10,
                         MU.EF=0, Het.I2=0.5)
meta.out <- rma(yi=meta.data$EF, sei=meta.data$SE, method="DL")
meta.out
forest(meta.out)
```

---

#### 4.8 Effects of sample size on the variation in $I^2$

We can use the `sim.meta.EF()` function to perform **Monte Carlo simulations and assess how well the simulated values of heterogeneity $I^2$ and the adjusted $\alpha$ threshold align with theoretical expectations** across different sample sizes. In these simulations, we vary the parameters 'N', 'M', and 'Het.I2', generating 1,000 datasets for each combination. Note that these simulations may take several minutes to complete, and the code spans two pages, which should be combined into a single block for execution.

```
require(metafor)
# Specify where the results file is saved
path <- "H:\\Knief\\Documents\\Stats\\MetaHeterogeneity\\"
# The number of studies N being meta-analytically summarized
Ns <- c(10,20,40,80)
# The sample size M per study
Ms <- c(10,20,40,80)
# Heterogeneity I2
I2s <- c(0,0.2,0.4,0.6,0.8)
# Number of simulations per N and M
nSims <- 1000
# ---------------------------------------------------------------------------
# Initiate final outfile
meta.out.all <- data.frame(matrix(rep(NA,1*14),ncol=14))
colnames(meta.out.all) <- c("N","M","exp.Het.I2","exp.EF","obs.beta",
                            "obs.beta.CI95.L","obs.beta.CI95.U",
                            "obs.beta.CI.N","obs.tau2","obs.P","obs.sB2",
                            "obs.I2","obs.critical.Z","obs.alpha.adj")
# sB2 can be regarded as the "typical" within-study variance of the observed
# effect sizes. It is similar to the squared mean standard error, i.e.
# (mean(meta.data$zSE))^2. It is calculated from equation 9 in
# Higgins & Thompson (2002)
sB2 <- function(SE, N) {
    ((N-1) * sum(1/SE^2)) / (sum(1/SE^2)^2 - sum((1/SE^2)^2)) }
meta.out.all <- meta.out.all[-1,]
write.table(meta.out.all, paste0(path,"data_MonteCarloSim_Heterogeneity.txt"),
    append=FALSE, row.names=FALSE, col.names=TRUE, sep="\t", quote=FALSE)
# Initiate counter and timer
tTotal = Sys.time()
k <- 1
for(n in 1:length(Ns)) {
  N <- Ns[n]
  for(m in 1:length(Ms)) {
    M <- Ms[m]
    for(i in 1:length(I2s)) {
      I2 <- I2s[i]
      for(s in 1:nSims) {
        meta.out.all[1,] <- NA
        meta.data <- sim.meta.EF(N=N, M=M, Het.I2=I2)
        meta.out <- rma(yi = meta.data$EF, sei = meta.data$SE, method = "DL")
        meta.out.all[1,1:7] <- c(N, M, I2, 0, meta.out$beta, meta.out$ci.lb,
                                 meta.out$ci.ub)
```

```
        meta.out.all[1,8] <- sum(c(meta.out$beta) >= meta.data$CI95.L &
                                 c(meta.out$beta) <= meta.data$CI95.U)
        meta.out.all[1,9:12] <- c(meta.out$tau2, meta.out$QEp,
                                  sB2(meta.data$SE,N), meta.out$I2/100)
        meta.out.all[1,13] <- qnorm(0.975) * sqrt(1 / (1 - meta.out$I2/100))
        meta.out.all[1,14] <- 2 * (1 - pnorm(meta.out.all[1,13]))
        write.table(meta.out.all, append=TRUE, sep="\t", quote=FALSE,
                    file=paste0(path,"data_MonteCarloSim_Heterogeneity.txt"),
                    row.names=FALSE, col.names=FALSE)
        # Print status
        flush.console()
        if(k %% 100 == 0) { print(paste0("Taking ",
          round(as.double(difftime(Sys.time(),tTotal,units="min")),digit=2),
          " minutes to finish ",k," out of ",
          nSims*length(Ns)*length(Ms)*length(I2s)," simulations")) }
        k <- k + 1
      }
    }
  }
}
```

We **plot the proportion of conventional 95% confidence intervals that contain the true effect against** $I^2$ using the `hexbin`, `grid`, `latticeExtra` and `RColorBrewer` packages in `R` (Carr *et al.* 2024, R Core Team 2024, Sarkar & Andrews 2022, Neuwirth 2022). As before, the code spans two pages and should be combined into a single block for execution.

```
if("latticeExtra" %in% rownames(installed.packages()) == FALSE)
  { install.packages("latticeExtra") }
if("hexbin" %in% rownames(installed.packages()) == FALSE)
  { install.packages("hexbin") }
if("grid" %in% rownames(installed.packages()) == FALSE)
  { install.packages("grid") }
if("RColorBrewer" %in% rownames(installed.packages()) == FALSE)
  { install.packages("RColorBrewer") }
library(latticeExtra)
library(hexbin)
library(grid)
library(RColorBrewer)

mycols <- brewer.pal(n=3, name="Set1")

# Specify where the results file is saved
path <- "H:\\Knief\\Documents\\Stats\\MetaHeterogeneity\\"
meta.out.all <- read.table(header=TRUE, sep="\t",
                           file=paste0(path,"data_MonteCarloSim_Heterogeneity.txt"))
# --------------------------------------------------------------------------------
# Plot the proportion of confidence intervals containing the true effect vs I^2
# The number of studies N being meta-analytically summarized
Ns <- as.numeric(names(c(table(meta.out.all$N))))
# The sample size M per study
Ms <- as.numeric(names(c(table(meta.out.all$M))))
```

```r
meta.out.all$prop.obs.beta.CI.N <- meta.out.all$obs.beta.CI.N / meta.out.all$N

for(n in 1:length(Ns)) {
  dat.n <- subset(meta.out.all, N==Ns[n])
  for(m in 1:length(Ms)) {
    dat <- subset(dat.n, M==Ms[m])
    Label <- paste(Ns[n], Ms[m], sep="_")

    # Predicted line
    exp.I2 <- seq(min(dat$obs.I2),max(dat$obs.I2),0.001)
    y <- sqrt(1/(1-exp.I2))
    Z.c <- qnorm(0.975) / y
    Type.I.rate <- 2 * (1 - pnorm(Z.c))
    Exp.Prop.CI.Beta <- 1 - Type.I.rate
    RealizedAlpha <- data.frame(exp.I2=exp.I2, y=y, Z.c=Z.c,
                                Type.I.rate=Type.I.rate,
                                Exp.Prop.CI.Beta=Exp.Prop.CI.Beta)

    # Calculate loess predictions of observed data
    fnc.loess.CI = loess(dat$prop.obs.beta.CI.N ~ dat$obs.I2, span=0.4)
    fnc.loess.obs = predict(fnc.loess.CI, exp.I2)

    # Hex plot
    # ----------------------------------------------------------------------
    svg(filename=paste(path,"FigS9_",Label,".svg",sep=""), height=90/25.4,
        width=90/25.4, family="Arial", pointsize=8)
    par(mar=c(2, 2.2, 1.5, 0.2), mgp=c(1.2, 0.2, 0))
    p <- hexbinplot(dat$prop.obs.beta.CI.N ~ dat$obs.I2, xbins = 40, aspect="1",
               tck=-0.5, colorkey=FALSE, xlim=c(-0.02,1.02), ylim=c(-0.02,1.02),
               colramp=colorRampPalette(gray(10:0 / 13)),
               xlab=expression(paste(I^2)),
               ylab=expression(paste("Proportion ",mu," covered by conventional
                                     95% CIs")),
               scales=list(x=list(at=seq(0,1,by=0.2),labels=seq(0,1,by=0.2)),
                           y=list(at=seq(0,1,by=0.2),labels=seq(0,1,by=0.2)),
                           tck=0.5),
               panel=function(x,y, ...) {
                 panel.hexbinplot(x, y, ...)
                 lattice::panel.points(y=Exp.Prop.CI.Beta, x=exp.I2, type="l",
                                       col=mycols[1], lwd=1.5)
                 lattice::panel.points(y=fnc.loess.obs, x=exp.I2, type="l",
                                       col=mycols[2])
               })
    print(p)
    dev.off()
  }
}
```

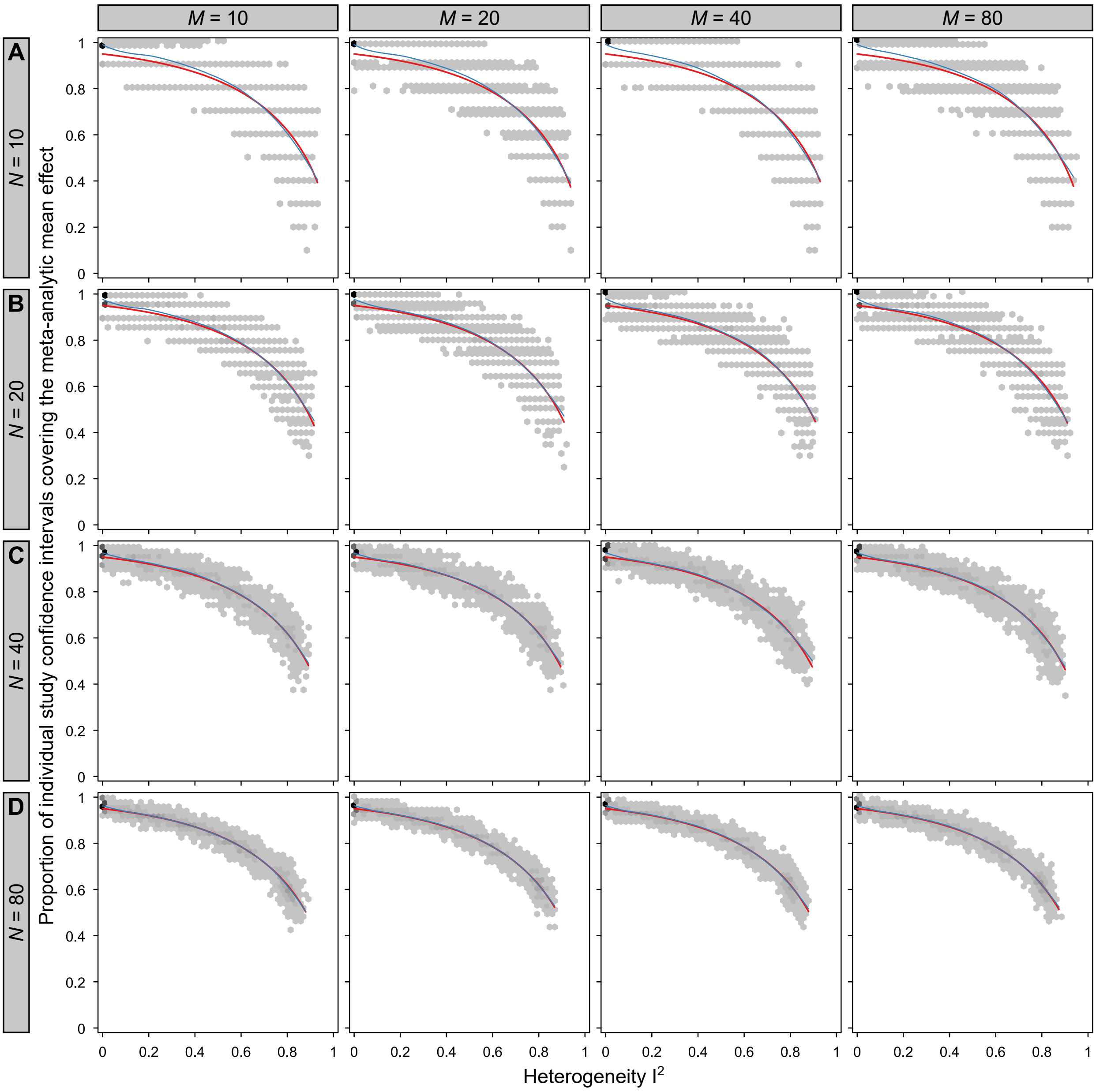

***Fig. S9*** *| Simulated and theoretical relationship between heterogeneity $I^2$ and the adjusted $\alpha$ threshold. The expected relationship is shown in red, while the relationship derived from 1,000 simulation runs is presented in blue for **(A)** 10, **(B)** 20, **(C)** 40 and **(D)** 80 studies per meta-analysis. For each of these meta-analyses, we varied the sample size per individual study from 10, 20, 40 to 80, as well as the amount of heterogeneity $I^2$ from 0, 0.2, 0.4, 0.6 to 0.8, resulting in a total of 80,000 simulation runs.*

#### 4.9 Simulation to demonstrate the precision of $SE_{MAMA}$

We can use the `sim.meta.EF()` function to perform **Monte Carlo simulations that demonstrate the precision of the $SE_{MAMA}$ derived confidence intervals in a many-analyses meta-analysis**. These simulations consider different numbers of research teams, sample sizes and varying levels of heterogeneity $I^2$. We simulate 1,000 datasets, each with $N = 20$, 50, 100, 200, or 500 research teams, sample sizes of $M = 100$, 200, or 500 and heterogeneity $I^2 = 0$, 0.5. Note that these simulations may take several minutes to complete, and as before, the code spans two pages and should be combined into a single block for execution.

```
require(metafor)
# Number of simulations
N.Sims <- 1000
# Number of research teams per MAMA
N.Analysts <- c(20,50,100,200,500)
# Sample size for individual studies in a MAMA
M.SampleSizes <- c(100,200,500)
# Heterogeneity
I2.Sims <- c(0,0.5)
# Path to where the outfile is saved
path <- "H:\\Knief\\Documents\\Stats\\MetaHeterogeneity\\"
# ----------------------------------------------------------------------------
MAMA.out <- data.frame(matrix(rep(NA,11*N.Sims*length(N.Analysts)
                                  *length(M.SampleSizes)
                                  *length(I2.Sims)), ncol=11))
colnames(MAMA.out) <- c("N","M","Het.I2","exp.EF","exp.SE","obs.EF",
                        "obs.SE","sigmaB2","sigmaC2","I2.Star","SE.MAMA")

# Initiate counter and timer
tTotal = Sys.time()
s <- 1

# Start the simulation loops
for(n in 1:length(N.Analysts)) {
  N.Analyst <- N.Analysts[n]
  for(m in 1:length(M.SampleSizes)) {
    M.SampleSize <- M.SampleSizes[m]
    for(i in 1:length(I2.Sims)) {
      I2.Sim <- I2.Sims[i]
      # Create 1000 datasets given a certain sample size per study
      sim <- sim.meta.EF(N=N.Sims, M=M.SampleSize, Dist.M="constant",
                         MU.EF=0, Het.I2=0)
      # Initiate the loop with N.Sims simulations
      for(j in 1:nrow(sim)) {
        # Create a dataset from the first of the N.Sims replicates (effect and
        # sample size) for N.Analysts analysts
        MAMA.data <- sim.meta.EF(N=N.Analyst, M=sim[j,"M"], Dist.M="constant",
                                 MU.EF=sim[j,"EF"], Het.I2=I2.Sim)
        MAMA <- rma(yi=MAMA.data$EF, sei=MAMA.data$SE, method="DL")
        # Cochran's Q
        CQ <- MAMA$QE
```

```
        k <- nrow(MAMA.data)
        # I2* and correction factor y
        I2.Star <- (CQ - k + 1) / CQ
        y.Star <- sqrt(1 / (1 - I2.Star))
        MAMA.out[s,"N"] <- N.Analyst
        MAMA.out[s,"M"] <- sim[j,"M"]
        MAMA.out[s,"Het.I2"] <- I2.Sim
        MAMA.out[s,"exp.EF"] <- sim[j,"EF"]
        MAMA.out[s,"exp.SE"] <- sim[j,"SE"]
        MAMA.out[s,"obs.EF"] <- MAMA$beta
        MAMA.out[s,"obs.SE"] <- MAMA$se
        MAMA.out[s,"sigmaC2"] <- var(MAMA.data$EF)
        MAMA.out[s,"I2.Star"] <- I2.Star
        MAMA.out[s,"sigmaB2"] <- MAMA.out[s,"sigmaC2"] * (1-I2.Star)
        MAMA.out[s,"SE.MAMA"] <- sqrt(MAMA.out[s,"sigmaB2"] +
                                      MAMA.out[s,"sigmaC2"] / N.Analyst)
        # Print status
        flush.console()
        if(s %% 100 == 0) { print(paste0("Taking ",
          round(as.double(difftime(Sys.time(),tTotal, units="min")),digit=2),
          " minutes to finish ",s," out of ",nrow(MAMA.out)," simulations")) }
        s <- s + 1
      }
    }
  }
}
write.table(MAMA.out, append=FALSE, sep="\t", quote=FALSE,
           file=paste0(path,"data_MonteCarloSim_SEMAMA.txt"),
           row.names=FALSE, col.names=TRUE)

# Test whether SE.MAMA provides a precise 95% confidence interval
MAMA.out <- read.table(header=TRUE, sep="\t",
                       file=paste0(path,"data_MonteCarloSim_SEMAMA.txt"))
rma(yi=MAMA.out$obs.EF, sei=MAMA.out$SE.MAMA, method="DL")
CI95.L <- MAMA.out$obs.EF - qnorm(0.975) * MAMA.out$SE.MAMA
CI95.U <- MAMA.out$obs.EF + qnorm(0.975) * MAMA.out$SE.MAMA
sum(0 >= CI95.L & 0 <= CI95.U) / nrow(MAMA.out)
```

We **plot the** $\mathbf{SE_{MAMA}}$**-based 95% confidence intervals and** $I^2$ **versus the number of research teams**. Please combine the three code blocks into one before execution.

```
if("metafor" %in% rownames(installed.packages()) == FALSE)
  { install.packages("metafor") }
if("RColorBrewer" %in% rownames(installed.packages()) == FALSE)
  { install.packages("RColorBrewer") }
if("Hmisc" %in% rownames(installed.packages()) == FALSE)
  { install.packages("Hmisc") }
library(metafor)
library(RColorBrewer)
library(Hmisc)
mycols <- brewer.pal(n=3, name="Set1")
```

```r
# Specify where the results file is saved
path <- "H:\\Knief\\Documents\\Stats\\MetaHeterogeneity\\"
MAMA.out <- read.table(header=TRUE, sep="\t",
                       file=paste0(path,"data_MonteCarloSim_SEMAMA.txt"))
# -----------------------------------------------------------------------------
# Plot the proportion of confidence intervals containing the true effect vs I^2
# The number of studies N being meta-analytically summarized
Ns <- as.numeric(names(c(table(MAMA.out$N))))
# The sample size M per study
Ms <- as.numeric(names(c(table(MAMA.out$M))))
# Heterogeneity per study
I2s <- as.numeric(names(c(table(MAMA.out$Het.I2))))

for(m in 1:length(Ms)) {
  dat.m <- subset(MAMA.out, M==Ms[m])
  for(i in 1:length(I2s)) {
  dat.m.i <- subset(dat.m, Het.I2==I2s[i])
  Label <- paste(Ms[m], I2s[i], sep="_")
  for.plot <- data.frame(matrix(rep(NA,length(Ns)*9),ncol=9))
  colnames(for.plot) <- c("N","I2Star","exp.I2Star","I2.L","I2.U","Prop",
                          "exp.Prop","P.L","P.U")
    for(n in 1:length(Ns)) {
      dat.m.i.n <- subset(dat.m.i, N==Ns[n])
      meta <- rma(yi=dat.m.i.n$obs.EF, sei=dat.m.i.n$SE.MAMA, method="DL")
      CQ <- meta$QE
      k <- nrow(dat.m.i.n)
      for.plot[n,"N"] <- Ns[n]
      for.plot[n,"I2Star"] <- (CQ - k + 1) / CQ
      for.plot[n,"exp.I2Star"] <- 0
      CQ.L <- qchisq(c(.025,.975),df=k-1, lower.tail=TRUE)[1]
      CQ.U <- qchisq(c(.025,.975),df=k-1, lower.tail=TRUE)[2]
      for.plot[n,"I2.L"] <- (CQ.L - k + 1) / CQ.L
      for.plot[n,"I2.U"] <- (CQ.U - k + 1) / CQ.U
      CI95.L <- dat.m.i.n$obs.EF - qnorm(0.975) * dat.m.i.n$SE.MAMA
      CI95.U <- dat.m.i.n$obs.EF + qnorm(0.975) * dat.m.i.n$SE.MAMA
      succ <- sum(0 >= CI95.L & 0 <= CI95.U)
      obs <- nrow(dat.m.i.n)
      exp.succ <- obs * 0.95
      for.plot[n,"Prop"] <- binconf(succ,obs)[1]
      for.plot[n,"exp.Prop"] <- 0.95
      for.plot[n,"P.L"] <- binconf(exp.succ,obs)[2]
      for.plot[n,"P.U"] <- binconf(exp.succ,obs)[3]
      print(confint(meta))
    }

    # Plots
    # -------------------------------------------------------------------------
    svg(filename=paste(path,"FigS10_",Label,"_Prop.svg",sep=""),
        height=70.55/25.4, width=70.55/25.4, family="Arial", pointsize=9.5)
    par(mar=c(2, 2.2, 1.5, 0.2), mgp=c(1.2, 0.2, 0), pty="s")
    p1 <- plot(for.plot$Prop ~ for.plot$N, tcl=-0.25, xlim=c(0,500),
```

```
              ylim=c(0,1.0), type="n", las=1,
              xlab=expression(paste("N Research teams")),
              ylab=expression(paste("Proportion ",mu," covered by extended
                                     95% CIs")))
    p2 <- polygon(c(for.plot$N,rev(for.plot$N)),
                  c(for.plot$P.L,rev(for.plot$P.U)), col='gray85', border=NA)
    p3 <- points(for.plot$exp.Prop ~ for.plot$N, type="l", col=mycols[1],
                 lwd=1, lty=2)
    p4 <- points(for.plot$Prop ~ for.plot$N, type="l", col=mycols[2], lwd=1)
    p5 <- points(for.plot$Prop ~ for.plot$N, type="p", pch=21, col=mycols[2],
                 lwd=1, cex=1.1, bg=mycols[2])

    print(p1) ; print(p2) ; print(p3) ; print(p4) ; print(p5)
    dev.off()

    # ---------------------------------------------------------------------------
    svg(filename=paste(path,"FigS10_",Label,"_I2.svg",sep=""), height=70.55/25.4,
        width=70.55/25.4, family="Arial", pointsize=9.5)
    par(mar=c(2, 2.2, 1.5, 0.2), mgp=c(1.2, 0.2, 0), pty="s")
    p1 <- plot(for.plot$I2Star ~ for.plot$N, tcl=-0.25, xlim=c(0,500),
              ylim=c(floor(min(for.plot$I2.L)*10)/10,1), type="n", las=1,
              xlab=expression(paste("N Analysts")),
              ylab=expression(paste("I2Star")))
    p2 <- polygon(c(for.plot$N,rev(for.plot$N)),
                  c(for.plot$I2.L,rev(for.plot$I2.U)), col='gray85', border=NA)
    p3 <- points(for.plot$exp.I2Star ~ for.plot$N, type="l", col=mycols[1],
                 lwd=1, lty=2)
    p4 <- points(for.plot$I2Star ~ for.plot$N, type="l", col=mycols[2], lwd=1)
    p5 <- points(for.plot$I2Star ~ for.plot$N, type="p", pch=21, col=mycols[2],
                 lwd=1, cex=1.1, bg=mycols[2])

    print(p1) ; print(p2) ; print(p3) ; print(p4) ; print(p5)
    dev.off()
  }
}
```

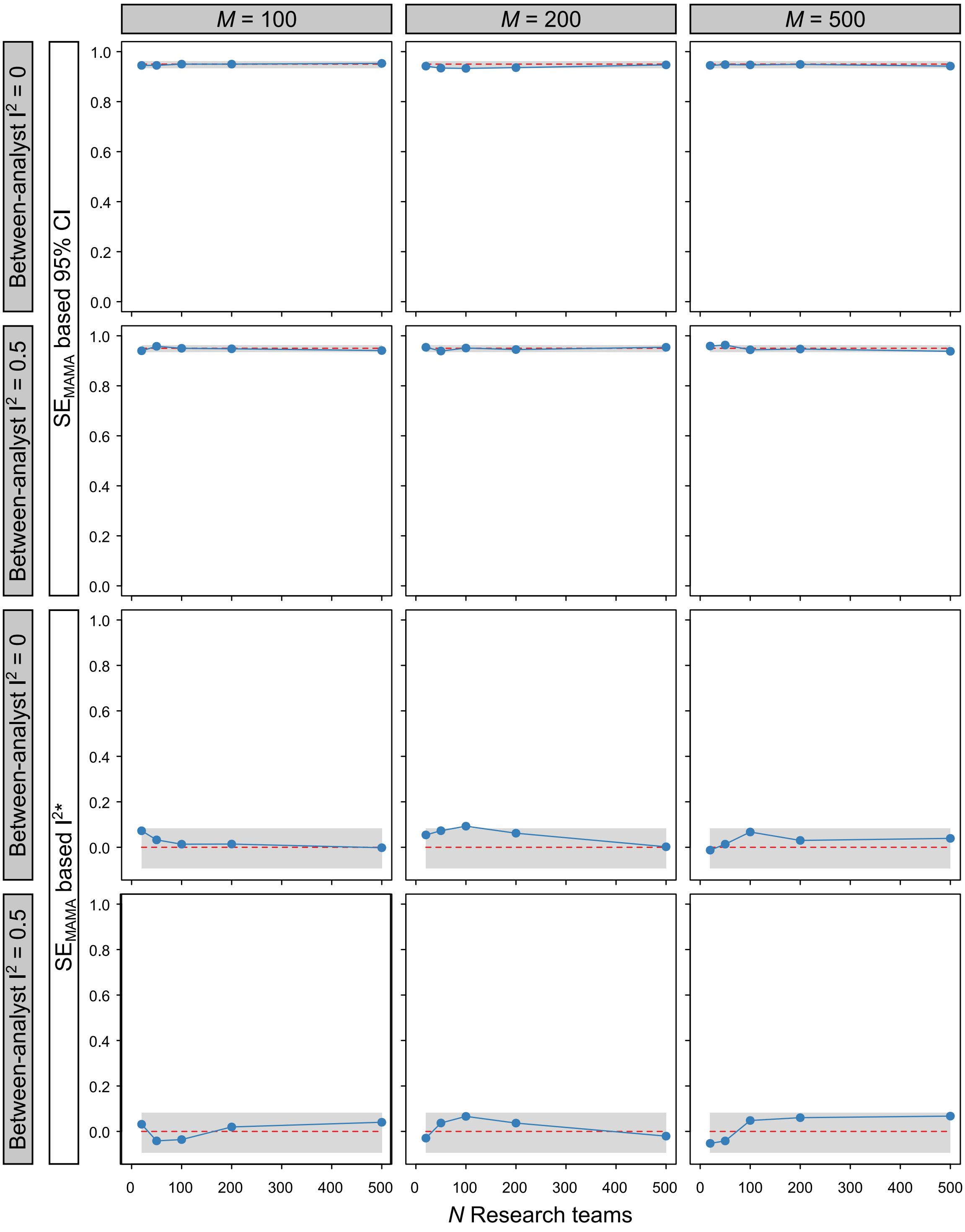

***Fig. S10*** | *Simulated and theoretical relationship between the $SE_{MAMA}$-based 95% confidence intervals and heterogeneity $I^2$ as a function of the number of research teams. The expected relationship (with the expected 95% CIs) is shown in red, while the relationship derived from 1,000 simulation runs is presented in blue. The expected 95% CIs for the $SE_{MAMA}$-based 95% CIs are binomial confidence intervals (Harrell Jr 2024). For the $SE_{MAMA}$-based $I^2$, we derive confidence intervals from the $\chi^2$ distribution with $k-1$ degrees of freedom, where $k$ is the number of studies included in the meta-analysis (Viechtbauer 2007). In this analysis, we varied the sample size per individual study from 100 to 200 and 500, and the amount of heterogeneity between research teams $I^2$ from 0 to 0.5, resulting in a total of 30,000 simulation runs.*

---

#### 4.10 Monte Carlo simulation of a many-analyses meta-analysis

We can simulate raw data with full control over fixed effects (e.g., their regression coefficients and collinearity), the random effect structure and the amount of random noise using the `simglm` package in `R` (LeBeau 2022). We fit mixed-effects models with the `lme4` and `lmerTest` packages (Bates *et al.* 2015, Kuznetsova *et al.* 2017), and increase the performance and parallelize the simulations using the `future.apply`, `parallel`, `progressr` and `data.table` packages (Barrett *et al.* 2024, Bengtsson 2021, Bengtsson 2023). Please combine all code blocks into one before execution.

```
rm(list=ls())
if("simglm" %in% rownames(installed.packages()) == FALSE)
  { install.packages("simglm") }
if("lme4" %in% rownames(installed.packages()) == FALSE)
  { install.packages("lme4") }
if("lmerTest" %in% rownames(installed.packages()) == FALSE)
  { install.packages("lmerTest") }
if("metafor" %in% rownames(installed.packages()) == FALSE)
  { install.packages("metafor") }
if("data.table" %in% rownames(installed.packages()) == FALSE)
  { install.packages("data.table") }
if("future.apply" %in% rownames(installed.packages()) == FALSE)
  { install.packages("future.apply") }
if("parallel" %in% rownames(installed.packages()) == FALSE)
  { install.packages("parallel") }
if("progressr" %in% rownames(installed.packages()) == FALSE)
  { install.packages("progressr") }

require(simglm)
require(lme4)
require(lmerTest)
require(metafor)
require(data.table)
require(future.apply)
require(parallel)
require(progressr)

# ---------------------------------------------------------------------------
# Settings ------------------------------------------------------------------

# Specify path and outfile names ---
path <- "H:\\Knief\\Documents\\Stats\\MetaHeterogeneity\\"
outfile <- "data_MonteCarlo_MAMA_10000"

# Define number of simulations ---
nSim <- 10000

# Set up parallel backend ---
# Uses all available cores minus 1
plan(multisession, workers = detectCores() - 1)

# Create all 2^7=128 decisions ---
```

```r
mds <- expand.grid(c(list(1:4),rep(list(1:2), 6)))
mds <- subset(mds, !(Var6 == Var7))

# -----------------------------------------------------------------------------
# Function for fitting lm() and lmer() models ---------------------------------

fit_mod <- function(dat, y, fixed, rand=NULL, reml=FALSE) {
  if(is.null(rand)) {
     pred <- c("treatment",fixed)
  } else {
     randstr <- sapply(rand, function(x) sprintf("(1|%s)",x))
     pred <- c("treatment",fixed,randstr)
  }
  form <- as.formula(paste(y," ~ ", paste(pred, collapse=" + ")))
  if(is.null(rand)) {
     lm(form, data=dat)
  } else {
     lmer(form, REML = reml, data=dat)
  }
}

# -----------------------------------------------------------------------------
# Function to check for complete factors --------------------------------------

chck_fac_lev <- function(df, predictors, factor_vars=c("treatment","sex")) {
  sub <- df[complete.cases(df[, predictors, drop=FALSE]), ]
  all(sapply(factor_vars, function(v) nlevels(droplevels(sub[[v]]))>=2))
}

# -----------------------------------------------------------------------------
# Function to run a single simulation -----------------------------------------

run_simulation <- function(j) {

  # Track progress ---
  p()

  # Use only valid datasets, i.e. those with fully represented factor levels
  # and where all 128 models converge ---
  valid_data <- FALSE
  rejected <- 0
  while (!valid_data) {
    # Generate data in a random effects model ---
    sim_arguments <- list(
      formula = y ~ 1 + treatment + weight + sex + wing + (1|year),
      fixed = list(treatment=list(var_type="factor",
                     levels=c("Control1","Control2","Treatment")),
                   weight=list(var_type="continuous",
                     dist='rgamma', shape=3),
                   sex=list(var_type="factor", levels=c("Female","Male"),
                     prob=c(0.15,0.85)),
```

```
                  wing = list(var_type="continuous", mean=55, sd=5)),
    error = list(variance=50),
    heterogeneity = list(variable="sex", variance=c(2,1)),
    randomeffect = list(int_id=list(variance=5/0.9, var_level=2)),
    sample_size = list(level1=20, level2=5),
    reg_weights = c(100,0,0, 0.48, 2.3, 0.21)
    # Control1, Control2, Treatment, Weight, Male, Wing
  )
  dat <- simulate_fixed(data=NULL, sim_arguments)
  dat <- simulate_randomeffect(dat, sim_arguments)
  dat <- simulate_error(dat, sim_arguments)
  dat <- simulate_heterogeneity(dat, sim_arguments)
  dat <- generate_response(dat, sim_arguments)
  NAPos <- sample(1:nrow(dat), nrow(dat)*0.2)
  dat$wing[NAPos] <- NA

  # Data Preparation ---
  dat$Sim <- j
  dat <- dat[, c("Sim","y","treatment","weight","sex","wing","year")]
  dat.raw <- as.data.table(dat)

  # Decision dataset preparations ---
  dat$sq.y <- sqrt(dat$y)
  dat$sex <- factor(dat$sex)
  datC <- subset(dat, treatment %in% c("Treatment","Control1"))
  datC$treatment <- ifelse(datC$treatment == "Control1",
                           "Control",datC$treatment)
  datC$treatment <- factor(datC$treatment,
                           levels=c("Control","Treatment"))
  dat$treatment <- ifelse(dat$treatment %in% c("Control1","Control2"),
                          "Control", dat$treatment)
  dat$treatment <- factor(dat$treatment, levels=c("Control","Treatment"))

  rem.min <- which(dat$y == min(dat$y))
  rem.max <- which(dat$y == max(dat$y))
  datOR <- dat[-c(rem.min, rem.max), ]
  rem.minC <- which(datC$y == min(datC$y))
  rem.maxC <- which(datC$y == max(datC$y))
  datCOR <- datC[-c(rem.minC, rem.maxC), ]

  # Regenerate dataset if it happens to be the same after outlier removal ---
  dat.cmp <- setorder(dat[complete.cases(
              dat[,c("y","treatment","weight","sex","wing","year")]),])
  datC.cmp <- setorder(datC[complete.cases(
              datC[,c("y","treatment","weight","sex","wing","year")]),])
  datOR.cmp <- setorder(datOR[complete.cases(
               datOR[,c("y","treatment","weight","sex","wing","year")]),])
  datCOR.cmp <- setorder(datCOR[complete.cases(
                datCOR[,c("y","treatment","weight","sex","wing","year")]),])
  if(identical(dat.cmp, datOR.cmp) || identical(datC.cmp, datCOR.cmp) ||
     identical(dat.cmp, datC.cmp) || identical(datOR.cmp, datCOR.cmp) ||
```

```
   identical(dat.cmp, datCOR.cmp) || identical(datC.cmp, datOR.cmp)) {
     rejected <- rejected + 1
     next # skip to next iteration of the while-loop and try a new dataset
   }

# Scale continuous variables ---
scale_data <- function(df) {
    df$weight <- scale(df$weight)
    df$wing <- scale(df$wing)
    return(df)
}

dat <- scale_data(dat)
datC <- scale_data(datC)
datOR <- scale_data(datOR)
datCOR <- scale_data(datCOR)

# Replace missing data with mean ---
R <- mean(dat$wing, na.rm=TRUE)
dat$wingR <- dat$wing
dat$wingR[is.na(dat$wing)] <- R
RC <- mean(datC$wing, na.rm=TRUE)
datC$wingR <- datC$wing
datC$wingR[is.na(datC$wing)] <- RC
ROR <- mean(datOR$wing, na.rm=TRUE)
datOR$wingR <- datOR$wing
datOR$wingR[is.na(datOR$wing)] <- ROR
RCOR <- mean(datCOR$wing, na.rm=TRUE)
datCOR$wingR <- datCOR$wing
datCOR$wingR[is.na(datCOR$wing)] <- RCOR

# Decision preparations ---
D12  <- list(dat, datC, datOR, datCOR)
D3   <- list("y", "sq.y")
D4   <- list("weight", NULL)
D5   <- list("sex", NULL)
D6   <- list("wing", "wingR")
D7.1 <- list("year", NULL)  # fixed
D7.2 <- list("year", NULL)  # random

# Store results ---
results_list <- vector("list", nrow(mds))

# Fit all 128 models ---
models_converged <- TRUE
for (i in 1:nrow(mds)) {

  # Determine dataset and variable names
  data_i <- D12[[mds[i,1]]]
  vars <- c("treatment", D3[[mds[i,2]]], D4[[mds[i,3]]],
            D5[[mds[i,4]]], D6[[mds[i,5]]])
```

```
if(!is.null(D7.1[[mds[i,6]]])) vars <- c(vars, D7.1[[mds[i,6]]])
if(!is.null(D7.2[[mds[i,7]]])) vars <- c(vars, D7.2[[mds[i,7]]])
if(!check_factor_levels(data_i, vars)) {
   models_converged <- FALSE
   break
}
options(warn=2)
mod <- tryCatch(fit_mod(dat=D12[[mds[i,1]]], y=D3[[mds[i,2]]],
                  fixed=c(D4[[mds[i,3]]], D5[[mds[i,4]]],
                          D6[[mds[i,5]]], D7.1[[mds[i,6]]]),
                  rand=D7.2[[mds[i,7]]]),
                  error=function(e) NULL
               )
options(warn=1)

# Check if any model failed (NULL means it did not converge) ---
if(inherits(mod, "merMod")) {
   # Check optimizer warnings, singularity, and Hessian definiteness
   hessian_valid <- tryCatch({
   all(eigen(mod@optinfo$derivs$Hessian, symmetric = TRUE)$values > 0)
   }, error=function(e) FALSE)

   if(!is.null(mod@optinfo$conv$lme4$messages) ||
      isSingular(mod, tol=1e-4) || !hessian_valid) {
        models_converged <- FALSE
        break
      }
   }

if(is.null(mod)) {
   models_converged <- FALSE
   break
}

coefs <- summary(mod)$coef
if(any(is.na(coefs["treatmentTreatment", c("t value", "Estimate")]))) {
   models_converged <- FALSE
   break
}

# Extract t-values and degrees of freedom from the model
# df are derived using the Satterthwaite approximation implemented in the
# lmerTest package in R. Note that we are talking about not the df of the
# whole model but specific df for that estimate
# See https://osf.io/kr2g9
t_treatment <- summary(mod)$coef["treatmentTreatment","t value"]
df_treatment <- ifelse(class(mod) == "lm", summary(mod)$df[2],
                       summary(mod)$coef["treatmentTreatment","df"])

# Convert the t-value and the degree of freedom (df) to the
# correlation coefficient r
```

```r
    r <- sqrt(t_treatment^2 / (t_treatment^2 + df_treatment))
    r <- ifelse(summary(mod)$coef["treatmentTreatment","Estimate"] < 0, -r, r)

    # Convert r and accompanying df to Zr and its sampling variance 1/(n-3)
    # where n=df+1
    Zr <- atanh(r)
    var.Zr <- 1 / (df_treatment + 1 - 3)
    sd.Zr <- sqrt(var.Zr)

    # Store decisions and Zr values
    results_list[[i]] <- list(
        Sim = j,
        D1 = ifelse(mds[i,1] %in% c(1,3), "C", "C1"),
        D2 = ifelse(mds[i,1] %in% c(1,2), "", "OR"),
        D3 = ifelse(is.null(D3[[mds[i,2]]]), "", D3[[mds[i,2]]]),
        D4 = ifelse(is.null(D4[[mds[i,3]]]), "", D4[[mds[i,3]]]),
        D5 = ifelse(is.null(D5[[mds[i,4]]]), "", D5[[mds[i,4]]]),
        D6 = ifelse(is.null(D6[[mds[i,5]]]), "", D6[[mds[i,5]]]),
        D7 = ifelse(is.null(D7.1[[mds[i,6]]]), "random", "fixed"),
        Zr = Zr,
        var.Zr = var.Zr
    )
  }

  # If any model failed, regenerate dataset ---
  if(!models_converged) {
     rejected <- rejected + 1
     next
  }

  # If everything is valid, break loop and proceed
  valid_data <- TRUE
}
cat("Simulation", j, "accepted after", rejected + 1, "attempt(s)\n")

# Combine results into a data.table ---
MAMA.data <- rbindlist(results_list)

# Multi-analyses meta-analysis (all 128 models=7 decisions):
# SEMAMA.a, SEMAMA.p, sigmaC, sigmaB, Type I Errors ---
MAMA.out <- rma(yi=MAMA.data$Zr, sei=sqrt(MAMA.data$var.Zr), method="DL")
sC <- var(MAMA.data$Zr)
CQ <- MAMA.out$QE
k <- nrow(MAMA.data)
I2STAR <- (CQ - k + 1) / CQ
sB <- sC * (1 - I2STAR)

# How many of the 128 variant Zr confidence intervals do not overlap 0?
CIL.Zr <- MAMA.data$Zr - qnorm(0.975) * sqrt(MAMA.data$var.Zr)
CIU.Zr <- MAMA.data$Zr + qnorm(0.975) * sqrt(MAMA.data$var.Zr)
No.Ov.95 <- 1 - sum(0 >= CIL.Zr & 0 <= CIU.Zr) / nrow(MAMA.data)
```

```r
  # Multi-analyses meta-analysis (a subset of 16 models = 4 decisions):
  # SEMAMA.a.4D and SEMAMA.p.4D ---
  MAMA.data.4D <- subset(MAMA.data, D3 == "y" & D4 == "weight" & D7 == "random")
  MAMA.out.4D <- rma(yi=MAMA.data.4D$Zr, sei=sqrt(MAMA.data.4D$var.Zr),
                     method="DL")
  sC.4D <- var(MAMA.data.4D$Zr)
  CQ.4D <- MAMA.out.4D$QE
  k.4D <- nrow(MAMA.data.4D)
  I2STAR.4D <- (CQ.4D - k.4D + 1) / CQ.4D
  sB.4D <- sC.4D * (1-I2STAR.4D)

  # Multi-analyses meta-analysis (a subset of 4 models = 2 decisions):
  # SEMAMA.a.2D and SEMAMA.p.2D ---
  MAMA.data.2D <- subset(MAMA.data, D3 == "sq.y" & D4 == "" & D5 == "sex" &
                         D6 == "wing" & D7 == "fixed")
  MAMA.out.2D <- rma(yi=MAMA.data.2D$Zr, sei=sqrt(MAMA.data.2D$var.Zr),
                     method="DL")
  sC.2D <- var(MAMA.data.2D$Zr)
  CQ.2D <- MAMA.out.2D$QE
  k.2D <- nrow(MAMA.data.2D)
  I2STAR.2D <- (CQ.2D - k.2D + 1) / CQ.2D
  sB.2D <- sC.2D * (1-I2STAR.2D)

  # Store multi-analyses meta-analysis results ---
  MAMA.out.res <- data.table(Sim=j, sC=sC, sB=sB, Ratio.sB.sC=sB / sC,
                             beta=as.numeric(MAMA.out$beta), SEMAMA=MAMA.out$se,
                             No.Ov.95=No.Ov.95,
                             SEMAMA.a=sqrt(sB + (sC / 7)),
                             SEMAMA.p=sqrt(sB + (sC / 128)),
                             beta.4D=as.numeric(MAMA.out.4D$beta),
                             SEMAMA.a.4D=sqrt(sB.4D + (sC.4D / 4)),
                             SEMAMA.p.4D=sqrt(sB.4D + (sC.4D / 16)),
                             beta.2D=as.numeric(MAMA.out.2D$beta),
                             SEMAMA.a.2D=sqrt(sB.2D + (sC.2D / 2)),
                             SEMAMA.p.2D=sqrt(sB.2D + (sC.2D / 4)))

  # Return the three data tables ---
  return(list(MAMA.data=MAMA.data, dat.raw=dat.raw, MAMA.out.res=MAMA.out.res))
}

# ------------------------------------------------------------------------------
# Run parallel simulations using future_lapply() -------------------------------
invisible(with_progress({
    p <- progressor(along = 1:nSim)
    all_results <- future_lapply(1:nSim, function(j) run_simulation(j),
                                 future.seed = TRUE)
}))

# Extract and combine individual results ---
dat.raw <- rbindlist(lapply(all_results, `[[`, "dat.raw"),
                     use.names=TRUE, fill=TRUE)
```

```r
dat.MA <- rbindlist(lapply(all_results, `[[`, "MAMA.data"),
                    use.names=TRUE, fill=TRUE)
dat.MAMA <- rbindlist(lapply(all_results, `[[`, "MAMA.out.res"),
                      use.names=TRUE, fill=TRUE)

# Write to disk ---
out <- list(dat.raw,dat.MA,dat.MAMA)
names(out) <- c("datRAW","datMA","datMAMA")
saveRDS(out, paste0(path,outfile,".rds"))
```
